# Loss of TREM2 exacerbates parenchymal amyloid pathology but diminishes CAA in Tg-SwDI mice

**DOI:** 10.1101/2023.11.04.565659

**Authors:** Rui Zhong, Yingzheng Xu, Jesse W. Williams, Ling Li

## Abstract

Alzheimer’s disease (AD) is a progressive neurodegenerative disease, and it is the most common cause of dementia worldwide. Recent genome-wide association studies (GWAS) identified TREM2 (triggering receptor expressed on myeloid cells 2) as one of the major risk factors for AD. TREM2 is a surface receptor expressed on microglia and largely mediates microglial functions and immune homeostasis in the brain. The functions of TREM2 in AD pathogenesis, including in the formation of the key pathology parenchymal amyloid-β (Aβ) plaques, have been investigated by introducing *Trem2* deficiency in AD mouse models. However, the role of TREM2 in cerebrovascular amyloidosis, in particular cerebral amyloid angiopathy (CAA) remains unexplored. CAA features Aβ deposition along the cerebral vessels, signifying an intersection between AD and vascular dysfunction. Using a well-characterized CAA-prone, transgenic mouse model of AD, Tg-SwDI (SwDI), we found that loss of TREM2 led to a marked increase in overall Aβ load in the brain, but a dramatic decrease in CAA in microvessel-rich regions, along with reduced microglial association with CAA. Transcriptomic analysis revealed that in the absence of *Trem2*, microglia were activated but trapped in transition to the fully reactive state. Like microglia, perivascular macrophages were activated with upregulation of cell junction related pathways in *Trem2*-deficient SwDI mice. In addition, vascular mural cells and astrocytes exhibited distinct responses to *Trem2* deficiency, contributing to the pathological changes in the brain of *Trem2*-null SwDI mice. Our study provides the first evidence that TREM2 differentially modulates parenchymal and vascular Aβ pathologies, which may have significant implications for both TREM2- and Aβ-targeting therapies for AD.

## Introduction

Triggering receptor expressed on myeloid cells 2 (TREM2) is a surface receptor from the immunoglobulin superfamily, and recent genome-wide association studies have identified *Trem2* as a major genetic risk factor for Alzheimer’s disease (AD) (Guerreiro *et al*., 2013; Jonsson *et al*., 2013; Jansen *et al*., 2019). TREM2 is expressed predominantly on microglia in the central nervous system and serves as a key regulator of microglial functions in normal aging and AD (Qu and Li, 2021). Recent studies using transgenic mouse models have shown a significant impact of *Trem2* deficiency on cerebral amyloid-β (Aβ) deposition, a neuropathological hallmark of AD. Interestingly, the impact of *Trem2*-deficiency is inconsistent, with some studies showing an increase but others a decrease in Aβ pathology upon *Trem2* deletion, although some of the discrepancies can be explained by the different stages of the development of Aβ pathology (Ulrich *et al*., 2014; Jay *et al*., 2015, 2017; Wang *et al*., 2015, 2016; Krasemann *et al*., 2017). In the meantime, it has been consistently observed that the Aβ plaque-associated microgliosis is reduced regardless of the stages of Aβ pathology development in the absence of *Trem2*, and recent transcriptomic studies have unveiled that *Trem2*-null microglia fail to switch to a disease-associated, reactive phenotype, leading to diffused amyloid plaques in the parenchyma (Ulrich *et al*., 2014; Jay *et al*., 2015, 2017; Wang *et al*., 2015, 2016; Krasemann *et al*., 2017; Sheng *et al*., 2019; Joshi *et al*., 2021). However, the role of TREM2 in cerebrovascular amyloidosis, in particular cerebral amyloid angiopathy (CAA), has not been investigated.

CAA, characterized by Aβ deposition in the cerebrovascular basement membrane, is a common pathological feature of brains with AD, present in approximately 80% of AD patients (Viswanathan and Greenberg, 2011; Charidimou, Gang and Werring, 2012; Yamada and Naiki, 2012). It triggers cerebrovascular inflammation, hemorrhages, microinfarcts, and cognitive impairment (Cisternas, Taylor and Lasagna-Reeves, 2019; Greenberg *et al*., 2020). AD subjects with CAA show a more rapid decline in cognitive test performance than those devoid of CAA (Pfeifer *et al*., 2002). In addition, CAA-related adverse effects hamper the development of Aβ-targeting immunotherapies, currently the most successful class of disease-modifying therapies for AD (Gandy, 2023). Compelling evidence indicates that CAA constitutes the underlying cause of the amyloid-related imaging abnormalities (ARIA), one of the main adverse events associated with anti-Aβ immunotherapies (Karran and De Strooper, 2022). Thus, a better understanding of cerebrovascular amyloidosis not only elucidates the pathogenesis of CAA per se but also facilitates the development of safe and efficacious therapies for AD.

Aβ is produced by the sequential cleavage of amyloid-β precursor protein (APP) by β-secretase (BACE1) followed by γ-secretase (O’Brien and Wong, 2011). Aβ40 and Aβ42 are the two most abundant products of the BACE1 cleavage and hence the two main species found in amyloid pathology in AD brains. A number of point mutations in APP have been identified to cause familial AD by elevating Aβ production, including the Swedish double mutation (K670N/M671L) (Mullan *et al*., 1992), and several other mutations cause familial CAA, including Dutch E22Q and Iowa D23N within the Aβ sequence of APP (Levy *et al*., 1990; Grabowski *et al*., 2001). Although Aβ40 and Aβ42 are the primary constituents of both CAA and parenchymal plaques, Aβ40 is more vasculotropic than Aβ42 and accumulates largely along blood vessels to form fibrillar CAA, whereas Aβ42 mainly forms dense fibrillar cores of the parenchyma plaques (Miller *et al*., 1993; Gravina *et al*., 1995). There is a growing body of evidence suggesting that cerebral vascular dysfunction precedes the buildup of amyloid plaques and tau tangles that eventually lead to cognitive deficits in the development of AD (Govindpani *et al*., 2019; Korte, Nortley and Attwell, 2020; Montagne *et al*., 2020). An early presence of CAA at the basement membrane of the vessel wall resulting from failed clearance of excess Aβ via the perivascular pathway is believed to impair the blood-brain barrier (BBB) function (Weller *et al*., 2008; Gireud-Goss *et al*., 2021; Chen *et al*., 2022). However, how vascular Aβ interacts with the neurovascular unit and other adjacent cell types remains incompletely understood. While recent studies highlight the importance of microglia in the pathogenic process of CAA (Scholtzova *et al*., 2017; Delaney *et al*., 2021; Kiani Shabestari *et al*., 2022), the specific role of TREM2 in CAA has not been explored.

Therefore, the current study was undertaken to examine the impact of *Trem2* deficiency on CAA and the neurovascular components, as well as parenchymal Aβ deposition in the brain. We used a well-characterized transgenic mouse model of CAA/AD, Tg-SwDI (Davis *et al*., 2004) (hereafter referred to as SwDI), which carries both the familial AD Swedish (Sw) and Dutch/Iowa (DI) mutations in the *APP* gene that drive the cerebrovascular accumulation of Aβ (CAA) with Aβ40 as the dominant species. SwDI mice develop diffused Aβ plaques in the parenchyma, as well as CAA, in various brain regions in an age-dependent manner (Davis *et al*., 2004). We generated the SwDI/Trem2 mouse line by breeding SwDI with *Trem2-/-* mice (Turnbull *et al*., 2006) and aged them till 16 months old when the pathologies are widespread and profound. We found that loss of *Trem2* led to a marked exacerbation of parenchymal Aβ deposition but a drastic diminishment of CAA, along with reduced CAA-associated microgliosis. Single nucleus transcriptomic analyses revealed that interactions of microglia, perivascular macrophages, and other vascular cells contributed to the shift from CAA to parenchymal plaques in *Trem2*-deficient mice. These findings demonstrate a differential role of TREM2 in modulating parenchymal and vascular amyloid pathology.

## Materials and Methods

### Animals

Tg-SwDI (SwDI) mice (C57BL/6-Tg(Thy1-APPSwDutIowa)BWevn/Mmjax) have been described previously (Davis *et al*., 2004), and *Trem2*-knockout (TKO) mice (Turnbull *et al*., 2006) were generously provided by Dr. Marco Colonna at the Washington University in St. Louis. The SwDI/Trem2 line was generated by the two-step breeding of SwDI mice with TKO mice, producing six genotypes: SwDI;Trem2-/- (SwDI/TKO), SwDI;Trem2+/− (SwDI/THet), SwDI;Trem2+/+ (SwDI/TWT), Trem2-/- (TKO), Trem2+/− (THet), and wild-type (WT). Mice carrying the SwDI transgene were used in the present study. Littermates were used whenever possible, and both males and females were included. All animal experiments were reviewed and approved by the Institutional Animal Care and Use Committee of the University of Minnesota.

### Immunohistochemical, immunofluorescent, and histochemical staining

Immunohistochemistry experiments were conducted as previously described (Cheng *et al*., 2013; Qu *et al*., 2022). Briefly, the posterior half of the mouse brains were collected and post-fixed with 4% paraformaldehyde for 48 hours. 50-μm coronal sections were obtained using a vibratome (Leica Microsystems Inc). For the immunohistochemical staining, the VECTASTAIN ABC kit (Vector Laboratories, PK-4002) was used following the manufacturer’s protocol. For immunofluorescent staining, sections were washed with PBS before blocking (5% normal donkey serum and 0.5% Triton X-100 in PBS) for one hour at room temperature followed by overnight incubation with primary antibodies. Sections were then washed and incubated with secondary antibodies for 3 hours at room temperature. After PBS washes, sections were mounted onto glass slides and sealed in Vectashield HardSet antifade mounting medium (Vector Laboratory). Methoxy-X04 (Tocris 4920) staining was conducted as previously described (Meilandt *et al*., 2020), following the incubation of secondary antibodies. The primary antibodies used include anti-Aβ antibodies 6E10 (Biolegend 803002), anti-GFAP (Dako Z0334), anti-IBA1 (Wako 019-19741), and anti-CD31 (R&D AF3628). The secondary antibodies for immunofluorescence include donkey anti-mouse IgG Alexa Fluor 568, anti-rabbit IgG Alexa Fluor 488 and 647, anti-rat IgG Alexa Fluor 488 and 647, and anti-goat IgG Alexa Fluor 488 (Invitrogen).

### Optical and fluorescent imaging and quantification

Four consecutive sections (350 μm apart) and two (300 μm apart) were selected for immunohistochemical and immunofluorescent quantifications, respectively, as described previously with some modifications (Cheng *et al*., 2013; Jeong *et al*., 2021; Qu *et al*., 2022). Briefly, for optical imaging, the images were acquired with a 4x objective lens and subsequently stitched with ImageJ/FIJI. The immunoreactivity of Aβ, IBA1, and GFAP in the cortex, hippocampus, and thalamus of the mouse brains was quantified with individual 4x images acquired with Image-Pro Plus (MediaCybernetics, Rockville, MD). These images were converted to 16-bit grayscale on Image J/FIJI and subsequently quantified with the same threshold. A minimum size of 30 μm^2^ was applied to filter out small nonspecific staining. For fluorescent imaging, sections were imaged using the Keyence all-in-one fluorescent microscope (Keyence, BZ-X810). The entire section was scanned under a 10x objective lens and a stitched image was produced in the Keyence Analysis software. Similar approaches were applied for the quantifications and representative images of CAA, CAA-associated microglia, and microvascular integrity. For quantifications, images were taken under a 20x objective with a field of view (FOV) of 720 μm x 540 μm. For representative images, a z-stack of images (8 μm ± 0.12 μm) with a z-step size of 0.2 um was taken under a 100x oil objective lens using the multi-stack module. At least six 20x images per animal from the same region in the thalamus were used for quantifications, and the staining was quantified per FOV with regions of interest (ROIs) to exclude non-target staining using ImageJ/FIJI. For the CAA-associated microglia quantification, the microglial association was defined as “within 5 μm of CAA immunostaining”, and ROIs were selected around CAA and expanded 5 μm in ImageJ/FIJI. For X04+ plaque density quantifications, fluorescent images were converted to 16-bit on grayscale, and gray values were quantified as a readout for fluorescent intensity representing plaque density. The same threshold was applied within each channel per experiment, and the rolling ball method was utilized to reduce background staining. Results were averaged across the sections for each animal.

### Whole mouse brain processing/clearing, immunostaining, and 3D imaging

Whole mouse brains were processed following the SHIELD protocol (Park *et al*., 2018). Samples were cleared for 1 day at 42°C with SmartBatch+ (LifeCanvas Technologies), a device employing stochastic electrotransport (Kim *et al*., 2015). Cleared samples were then actively immunolabeled using SmartBatch+ (LifeCanvas Technologies) based on eFLASH technology integrating stochastic electrotransport (Kim *et al*., 2015) and SWITCH (Murray *et al*., 2015). Each brain sample was stained with primary antibodies, 5 μg of mouse anti-Aβ antibody (Encor, #MCA-AB9), 5 μg of rabbit anti-IBA-1 monoclonal antibody (Cell Signaling Technologies, #17198S), and 10 μg of goat anti-CD31 (R&D Systems, # AF3628) followed by fluorescently conjugated secondary antibodies in 1:2 (primary:secondary) molar ratios (Jackson ImmunoResearch). After active labeling, samples were incubated in EasyIndex (LifeCanvas Technologies) for refractive index matching (RI=1.52) and imaged at either 3.6X or 15X with a SmartSPIM axially-swept light sheet microscope (LifeCanvas Technologies).

### Immunoblotting

The experimental procedures for protein assay and immunoblotting have been previously described (Cheng *et al*., 2013; Jeong *et al*., 2021; Qu *et al*., 2022). Briefly, protein concentrations from total brain tissue homogenates were determined by the Bradford assay (Thermo Fisher 23246). Proteins were then separated by 12% SDS-PAGE and transferred to PVDF membranes. After blocking, the membranes were incubated in primary antibodies overnight. The primary antibodies used include BACE1 (Invitrogen PA1-757), GFAP (Aves labs), IBA1 (Wako 016-20001), GAPDH (Invitrogen AM4300), and tubulin (Sigma-Aldrich T5198). Subsequently, the membranes were incubated with HRP-conjugated secondary antibodies, followed by incubation in the Clarity Western ECL substrate (Bio-rad) for signal detection using the iBright Western Blot Imaging System (Thermo Fisher). Densitometric analysis was performed using the ImageJ software.

### Aβ species-specific enzyme-linked immunosorbent assay (ELISA)

The anterior half of the cortical hemisphere was homogenized for immunoblot analysis (see above) and Aβ ELISA. For the ELISA fraction, homogenized samples were further sequentially separated into the carbonate-soluble and guanidine-soluble fractions as previously described (Cheng *et al*., 2013; Jeong *et al*., 2021; Qu *et al*., 2022). The levels of Aβ40 and Aβ42 were measured using Aβ40- and Aβ42-specific ELISA kits (Invitrogen KHB3481 and KHB3441) according to the manufacturer’s protocol.

### Nuclei Isolation from flash-frozen mouse brain tissue

Nuclei from six female mouse brain cortical tissues (three for each genotype) were isolated following the 10x Genomics Demonstrated Protocol (CG000375; Nuclei Isolation from Complex Tissues for Single Cell Multiome ATAC + Gene Expression Sequencing), with the following modifications: the NP40 Lysis Buffer was used at a 20X dilution, tissue dissociation was performed on the gentleMACS Octo Dissociator with Heaters (Miltenyi Biotec, Bergisch Gladbach, North Rhine-Westphalia, Germany) using a modified version of the 4C_nuclei_1 program, and no nuclei permeabilization was performed. Nuclei were not sorted before capture.

### Single nucleus RNA-seq library construction using the 10x Genomics Chromium platform

Single nucleus RNA-seq (snRNA-seq) libraries were prepared per the Single Cell 3’ v3.1 Reagent Kits User Guide (10x Genomics, Pleasanton, California) using the 10x Genomics Chromium Controller, X, or Connect. Barcoded sequencing libraries were quantified by quantitative PCR using the Collibri Library Quantification Kit (Thermo Fisher Scientific, Waltham, MA). Libraries were sequenced on a NovaSeq 6000 (Illumina, San Diego, CA) as per the Single Cell 3’ v3.1 Reagent Kits User Guide, with a sequencing depth of ∼40,000 reads/nucleus. Estimated number of nuclei sequenced from individual samples range from 5700 to 8100.

### Initial Data Analysis

The demultiplexed raw reads were aligned to the transcriptome using STAR (version 2.5.1) (Dobin *et al*., 2013) with default parameters, using mouse mm10 transcriptome reference from Ensembl version 84 annotation, containing all protein-coding and long non-coding RNA genes. Expression counts for each gene in all samples were collapsed and normalized to unique molecular identifier (UMI) counts using Cell Ranger software version 4.0.0 (10X Genomics). The result is a large digital expression matrix with cell barcodes as rows and gene identities as columns.

### Count matrix generation and preprocessing

Sequencing read alignment was performed using *cellranger* (v6.1.2) *count* pipeline, where FASTQ files generated from *cellranger mkfastq* were mapped to a murine reference transcriptome (https://cf.10xgenomics.com/supp/cell-exp/refdata-gex-mm10-2020-A.tar.gz). *cellranger count* output containing filtered count matrix was adopted into R by Seurat (v4.0.1) (Butler *et al*., 2018) *Read10X* function. Cells with less than 200 total feature reads or possessing more than 25% mitochondrial mRNA content were removed as potential empty or apoptotic cells. Prior to scRNAseq dataset integration, the DoubletFinder package (McGinnis, Murrow and Gartner, 2019) was used to further refine doublet detection. Filtered scRNAseq objects were then merged and followed by normalization and scaling using Seurat *NormalizeData* and *ScaleData* functions with default parameters. Calculation of the top 3000 features that exhibit significant cell-to-cell variations was done by *FindVariableFeatures* function. Overall, 23483 genes and 36000 nuclei remained after filtering.

### Integration, clustering, and cell type annotation

First, linear dimensional reduction and principal component analysis (PCA) were performed on the merged dataset that passed previous preprocessing filters. The principal components (PC) were ranked based on explained variance of each PC to determine the most meaningful number of PCs (nPCs), which was reflected by the turning point in an elbow diagram by *Elbowplot*. Collectively, the top 20 PCs were able to cover the most meaningful PCs. Then, batch effects were corrected using the Harmony package (Korsunsky *et al*., 2019) *RunHarmony* function and projected dataset in UMAP and tSNE embeddings. This integration pipeline was adopted from Seurat and Harmony vignettes. To avoid manual annotation of clustered dataset that may involve biased opinions, a computational approach employing the *map_sampling* function of *scrattch.hicat* package (Yao *et al*., 2021) was implemented. Cells in our dataset were bootstrap-mapped with a mouse cortex RNAseq dataset reported by (Tasic *et al*., 2016). The final cell type predictions were evaluated by the authors using reported marker genes. Initial clustering of all nuclei was performed using a resolution of 0.6. Further rec-lustering of vascular cells, microglia and macrophages used resolutions of 0.6, 0.3, and 0.4. Over-clustered populations were refined and merged according to the signature gene expressions.

### Differential gene expression and pathway analysis

Differential expression (DE) analysis was performed using Seurat *FindMarkers* and *FindAllMarkers* functions. Log fold change threshold was set to 0 to avoid gene filtering by the DE function. The *fgsea* function of the *fgsea* package was utilized to profile pathway analysis. The Molecular Signature Database (MSigDB) containing all known annotated gene sets was imported to be used with *fgsea*. Then, log fold change values generated from DE algorithms were added to their corresponding gene symbols. Next, the named array was rank-ordered in *fgsea*, and MSigDB gene sets were subsequently mapped onto the ranked array. Statistical significance of the pathways was determined as adjusted p-value < 0.1. The classification of significant pathways (**Table S1**) was based on keywords identified by the authors for each respective class, and was manually examined for errors.

### Statistical analysis

Statistical tests, including one-way ANOVA with Tukey’s post-hoc test and student’s t-test, were carried out with GraphPad Prism 9, and data are presented in boxplots. The box extends from the 25th to 75th percentiles and the whiskers from the minimum and to the maximum value. The p-value < 0.05 was considered significant.

## Results

### *Trem2* deletion exacerbates overall amyloid deposition and production

The impact of *Trem2* deficiency on amyloid pathology was evaluated in a cohort of 16-month-old SwDI/Trem2 mice matched by sex and genotype. Immunohistochemical (IHC) analyses showed that amyloid deposition measured by 6E10 immunoreactivity was significantly exacerbated in SwDI/TKO compared to SwDI/TWT mice in cortex (**Fig. 1A-B**), hippocampus (**Fig. 1A, C**), and thalamus (**Fig. 1D-E**). Quantifications of 6E10 immunoreactive area showed an approximately two-fold increase in the amyloid load in SwDI/TKO compared to SwDI/TWT in all three quantified regions (**Fig. 1F-H**). Sub-analysis of the plaque size distribution showed the same trend when plaques were divided into small (< 20 µm^2^), medium (20-40 μm^2^), and large (> 40 μm^2^) in cortical images (**Fig. S1A-C**). Interestingly, there was no significant difference between SwDI/TWT and SwDI/THet mice, indicating that one copy of *Trem2* is sufficient to keep the amyloid pathology at bay.

**Fig. 1.**
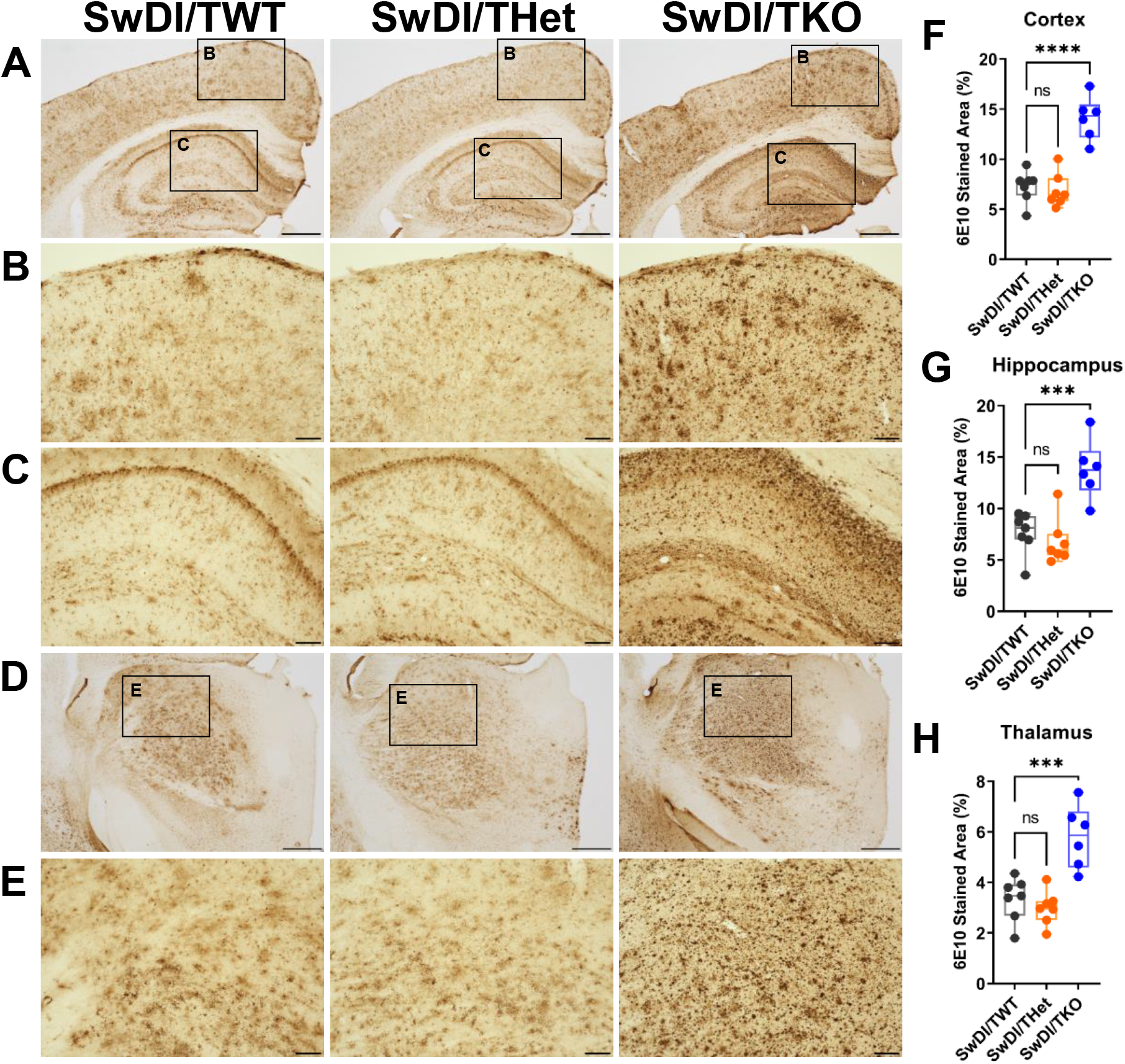
*Trem2* deletion exacerbates total amyloid deposition in SwDI mice. **A, D** Representative images of the immunohistochemical staining of amyloid by 6E10 in (**A**) cortex, hippocampus, and (**D**) thalamus in SwDI/TWT (n=7), SwDI/THet (n=7), and SwDI/TKO (n=6). Scale bars, 500 μm. **B, C, E** Selected zoomed-in areas from the representative images of (**B**) cortex, (**C**) hippocampus, and (**E**) thalamus. Scale bars, 100 μm. **F-H** Quantifications of the immunoreactive area of 6E10 in SwDI/TWT, SwDI/THet, and SwDI/TKO in (**F**) cortical, (**G**) hippocampal, and (**H**) thalamic regions, respectively. One-way ANOVA and Tukey’s post-hoc test. ***p<0.001, ****p<0.0001. ns, not significant.

To further investigate the underlying reasons for the increase in amyloid deposition, Aβ40- and Aβ42-specific quantitative ELISA were performed. In SwDI mice, Aβ40 accumulates more aggressively than Aβ42 in the brain parenchyma and cerebral vasculatures driven by the Dutch and the Iowa familial AD mutations (Davis *et al*., 2004). Therefore, Aβ40 and Aβ42 levels were evaluated in the carbonate soluble and insoluble (guanidine-soluble) fractions. As expected, Aβ40, Aβ42, and the Aβ40/Aβ42 ratio all significantly increased in SwDI/TKO mice compared with SwDI/TWT in both carbonate soluble and insoluble fractions (**Fig. 2A-D**). In addition, consistent with the immunostaining results (**Fig. 1**), there was no significant difference between SwDI/TWT and SwDI/THet mice (**Fig. 2A-D**). Thus, subsequent analyses focused on the comparisons between SwDI/TWT and SwDI/TKO groups. Additionally, immunoblot analysis showed a significant increase in BACE1 level in the SwDI/TKO group compared with SwDI/TWT (**Fig. 2E-F**), suggesting that enhanced amyloidogenic processing of APP contributes to the elevated levels of Aβ40 and Aβ42 in SwDI/TKO mice.

**Fig. 2.**
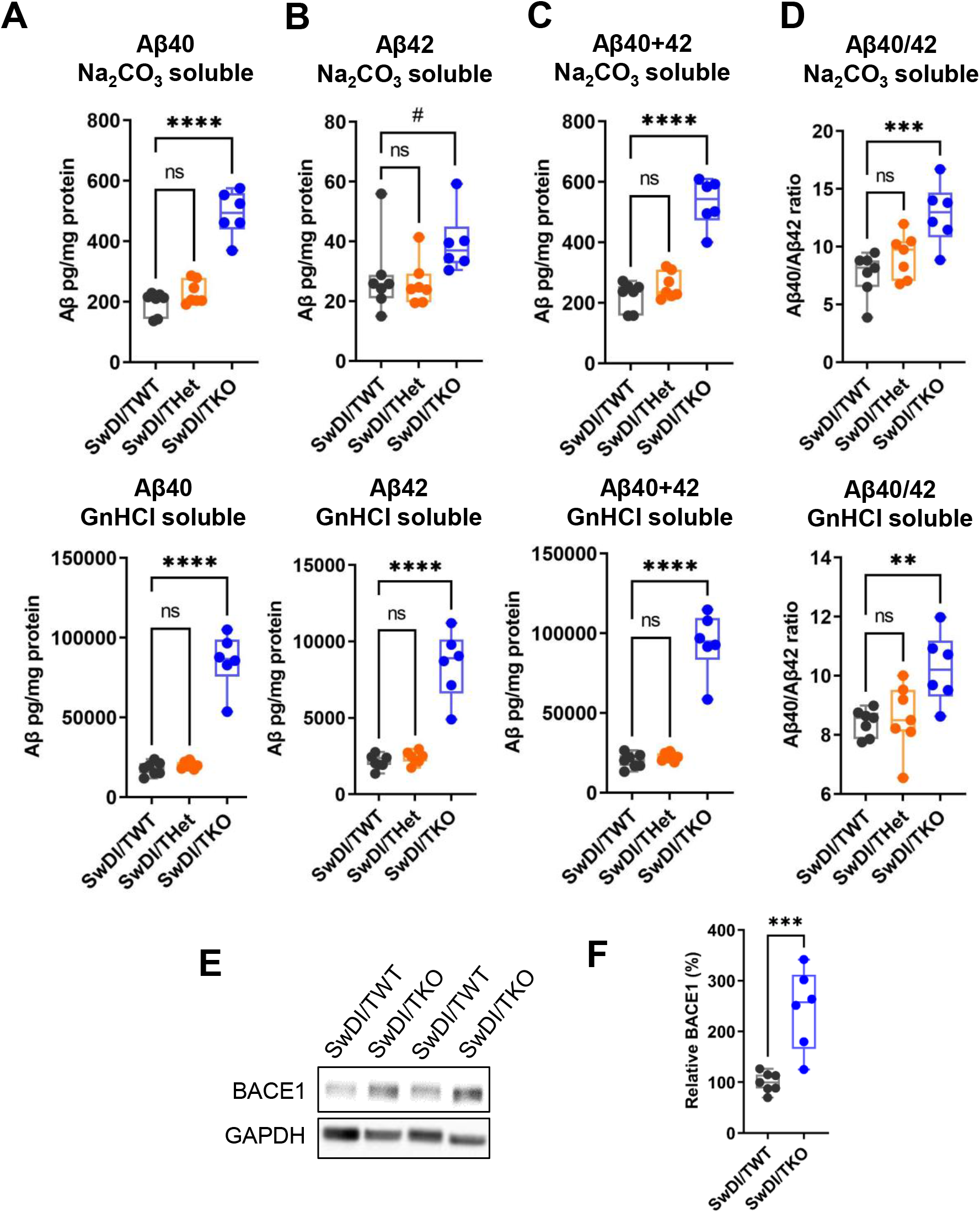
*Trem2* deletion elevates overall Aβ levels in SwDI mice and induces APP processing. **A-D** ELISA on Aβ40 and Aβ42 in SwDI/TWT (n=7), SwDI/THet (n=7), and SwDI/TKO (n=6) groups. The levels of (**A**) Aβ40 and (**B**) Aβ42, (**C**) total Aβ40 and Aβ42, and (**D**) the ratio of Aβ40/Aβ42 in both carbonate soluble and guanidine soluble fractions were all elevated in the SwDI/TKO mice whereas SwDI/THet mice showed no difference compared to SwDI/TWT. One-way ANOVA and Tukey’s post-hoc test. **E-F** Immunoblot analysis of BACE1. Representative images of the BACE1 immunoblot (**E**) and the quantification of BACE1 normalized by GAPDH (**F**) showed a significant increase in BACE1 level in SwDI/TKO mice compared with SwDI/TWT mice. Unpaired student’s t-test, two-tail. #p<0.1, **p<0.01, ***p<0.001, ****p<0.0001. ns, not significant.

### *Trem2* deletion increases total fibrillar Aβ but decreases CAA

The elevated Aβ40/Aβ42 ratio in the absence of TREM2 (**Fig. 2D**) suggested a potential increase in CAA. To quantify CAA, methoxy-X04 (X04) was used to stain fibrillary Aβ co-stained with CD31 to label the vascular endothelium to aid the visualization of CAA. The results showed that in the cortical area, the X04-stained amyloid fibrils/compact plaques were minimal in SwDI/TWT (**Fig. 3A-B**) despite the abundance of 6E10+ immunostaining (**Fig. 1**), confirming earlier reports that the parenchymal amyloid deposits are diffuse in nature resulting from the high level of vasculotropic Aβ40 in SwDI mice (Davis *et al*., 2004; Miao *et al*., 2005). The same studies also reported CAA largely present in thalamus and subiculum, although with minimal X04+ plaque load as in SwDI/TWT mice (**Fig. 3A-B, F**). In contrast, total X04^+^ amyloid fibrils were markedly increased in the cortex, hippocampus, and thalamus of SwDI/TKO mice (**Fig. 3A-E, Fig. S2A-B**), corresponding to the increase in total amyloid deposition by 6E10-immunostaining (**Fig 1**). However, intriguingly, there was a marked decrease of CAA in *Trem2*-deficient mice in the thalamus (**Fig. 3F-G**) where CAA was most abundant, indicating that the increase in Aβ40/Aβ42 or the overall increase of total Aβ (**Fig. 2**) did not aggravate but rather diminished CAA in SwDI/TKO mice. To confirm this unexpected finding, whole brain hemispheres of SwDI/TKO and SwDI/TWT littermates were subjected to brain clearing for 3D imaging. The brains were processed via SHIELD, SmartClear, immunostaining procedures to label Aβ plaques, microglia (IBA1), and blood vessels (CD31) and imaged using a SmartSPIM axially-swept light sheet microscope. The results clearly showed a dramatic decrease of CAA in the SwDI/TKO brain compared to the SWDI/TWT brain, despite significantly increased overall Aβ load in SwDI/TKO (**Fig. 3H**). To further understand if the compactness of fibrillar amyloid changes following a shift from CAA to diffused plaques when *Trem2* is deleted, the fluorescent intensity of each individual X04^+^ amyloid staining in the thalamus was analyzed. The results showed a significant decrease in the average maximal intensity and an increase in the average minimal intensity SwDI/TKO mice compared to SwDI/TWT (**Fig. S2C-D**), indicating that *Trem2* deficiency reduced the core density of amyloid fibrils.

**Fig. 3.**
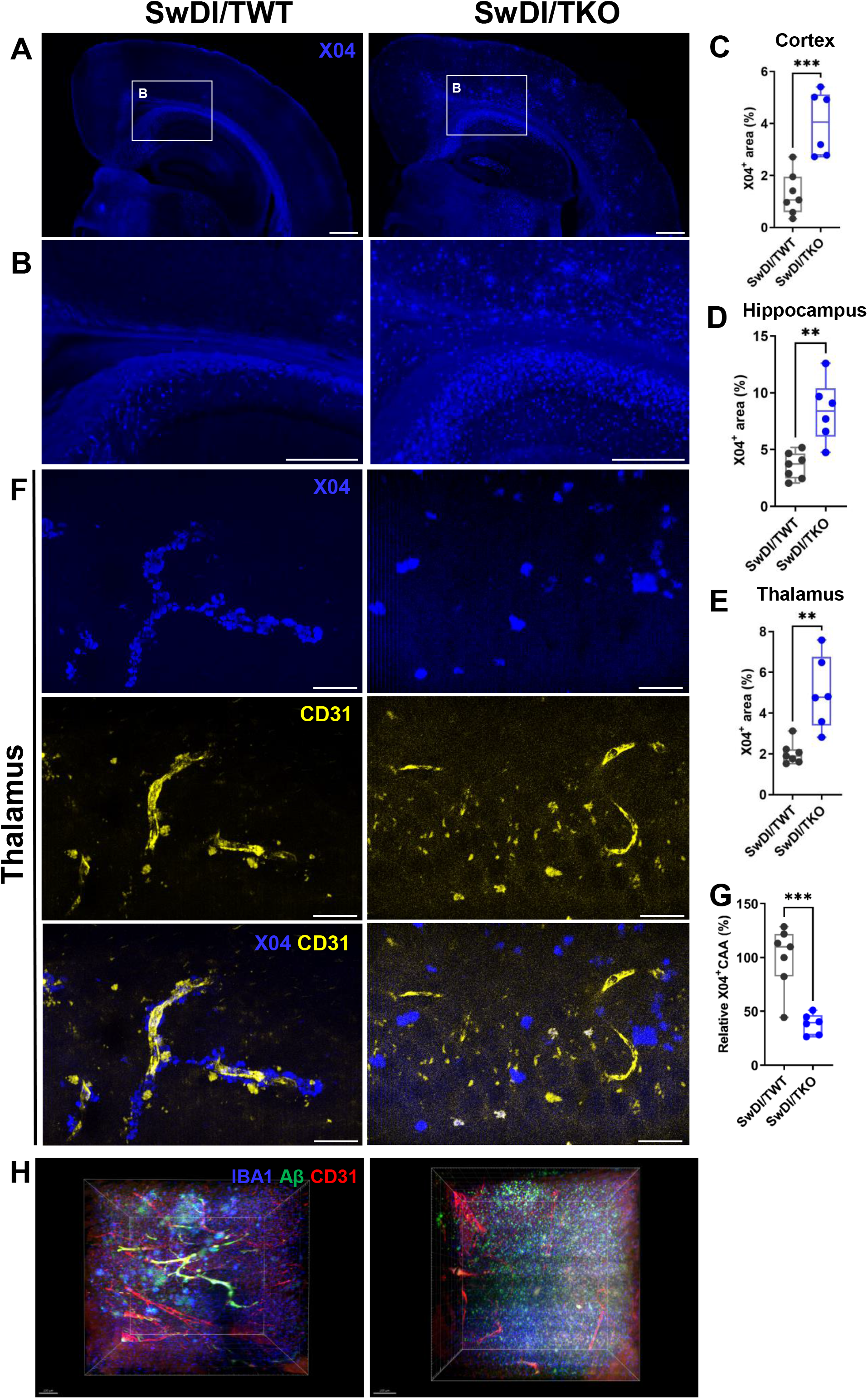
*Trem2* deletion increases total amyloid fibrils and decreases CAA in SwDI mice. **A** The representative stitched images of the fluorescent staining of methoxy-X04 under the 10x objective lens. Scale bars, 500 μm. **B** Selected zoomed-in areas of (**A**). Scale bars: 300 μm. **C-E** Quantifications of the X04+ stained area in SwDI/TWT (n=7) and SwDI/TKO (n=6) in (**C**) cortex, (**D**) hippocampus, and (**E**) thalamus, respectively. **F** Representative images of CAA co-stained with capillaries (CD31) and amyloid fibrils (X04) in the thalamus. Scale bars: 20 μm. **G** Quantification of CAA, showing significant decrease of CAA in SwDI/TKO compared to SwDI/TWT. **H** Representative 3D images of whole brain clearing and immunostaining with anti-Aβ, IBA1, and CD31 antibodies. While endothelial marker CD31 (red) and Aβ (green) co-stained CAA microvessels (yellow) are clearly seen in the thalamus image of SwDI/TWT, almost no CAA is visible in the thalamus image of SwDI/TKO, despite significantly more Aβ load (green) in SwDI/TKO. Scale bars, 200 μm. Unpaired student’s t-test, two-tail. **p<0.01, ***p<0.001.

### *Trem2* deletion reduces microgliosis in thalamus and decreases CAA-associated microglia

TREM2 plays a crucial role in regulating microglial activation and function, and loss of TREM2 modifies the response of microglia to Aβ. To assess the impact of *Trem*2 deletion on microgliosis in SwDI mice, brain sections were subjected to IBA1 immunostaining. Interestingly, IHC analysis showed no changes in global microgliosis in SwDI/TKO compared to SwDI/TWT and SwDI/THet mice in the cortex or the hippocampus (**Fig. 4A, D-E**), although microglial clustering was clearly reduced in the hippocampal CA1 stratum oriens region in SwDI/TKO mice (**Fig. S3A-B**). Intriguingly, microgliosis was significantly decreased in the thalamus of SwDI/TKO mice (**Fig. 4B-C, F**). Immunoblot analysis of the cortical homogenates also confirmed that IBA1 level was not significantly different between SwDI/TKO and SwDI/TWT mice (**Fig. 4G-H**). Consistent with Aβ-staining results (**Fig. 1**), no significant differences in IBA1 staining were found between SwDI/TWT and SwDI/THet. Therefore, SwDI/THet was not included in the following experiments assessing microglial pathology in the thalamus.

**Fig. 4.**
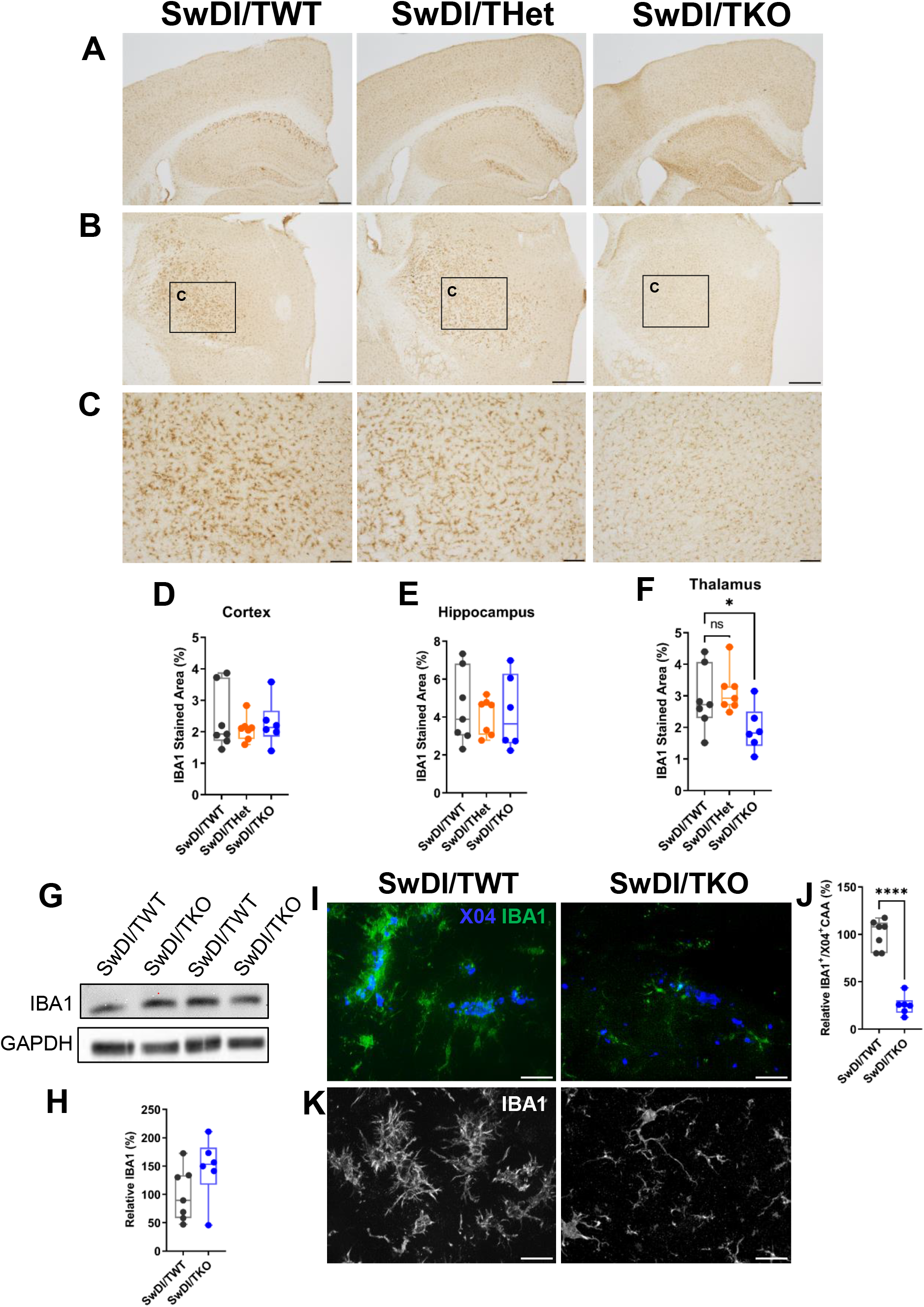
*Trem2* deletion reduces microgliosis and CAA-associated microglia in the thalamus. **A-C** Representative images of the immunohistochemical staining of microglia by IBA1 in (**A**) cortex and hippocampus, and (**B**) thalamus in SwDI/TWT (n=7), SwDI/THet (n=7), and SwDI/TKO (n=6). Scale bars, 500 μm. (**C**) Selected zoomed-in areas from (**B**). Scale bars, 100 μm. **D-F** Quantifications of the immunoreactive area of IBA1 in SwDI/TWT, SwDI/THet, and SwDI/TKO in (**D**) cortex, (**E**) hippocampus, and (**F**) thalamus, respectively. Only in thalamus was a significant decrease observed in IBA1 in SwDI/TKO mice. One-way ANOVA and Tukey’s post-hoc test. **G-H** Immunoblot analysis of IBA1. Representative images of the IBA1 immunoblot (**G**) and the quantification of IBA1 normalized by GAPDH (**H**). **I-J** CAA-associated microglia in the thalamus. (**I**) Representative images of CAA and microglia stained with X04 and IBA1, respectively, and (**J**) the quantification. **K** Representative images of microglial morphology in thalamus. Microglia adopt a less reactive morphology with longer process and smaller soma in SwDI/TKO than SwDI/TWT mice. Unpaired student’s t-test, two-tail. *p<0.05, ****p<0.0001. ns, not significant.

As Aβ deposits in the thalamus are predominantly CAA, brain sections were co-stained with X04 and IBA1 to determine if the reduction of microgliosis corresponds to the reduction of CAA in these regions. Indeed, X04 and IBA1 were completely co-localized in the thalamus of SwDI/TWT mice (**Fig. 4I**), indicating the region-specific reduction of microgliosis in SwDI/TKO corresponds to the location of diminished CAA pathology. Consistently, CAA-associated microglia were significantly reduced in the thalamus of the SwDI/TKO compared with SwDI/TWT (**Fig. 4I-J**), aligning with previous literature showing reduced plaque-associated microglia in other *Trem2*-deficient transgenic AD mouse models (Ulrich *et al*., 2014; Jay *et al*., 2015, 2017; Wang *et al*., 2015, 2016; Krasemann *et al*., 2017; Parhizkar *et al*., 2019; Meilandt *et al*., 2020). Additionally, microglia adopted a less reactive morphology with smaller soma and longer processes in SwDI/TKO than SwDI/TWT mice (**Fig. 4K**). Together, these data suggest that the lack of TREM2 prevents microglia from switching to a reactive state and impairs the ability of microglia to engage CAA.

### *Trem2* deletion aggravates cortical and hippocampal astrogliosis

Astrocytes, as well as microglia, respond to amyloidosis and regulate neuroinflammation. To assess the impact of *Trem2* deficiency on astrogliosis, brain sections of different genotypes of mice were subjected to GFAP immunostaining. The results showed that astrogliosis was significantly increased in cortical and hippocampal regions in SwDI/TKO compared to SwDI/TWT mice, whereas no change was seen in SwDI/THet mice (**Fig. 5A-C, E-F**), consistent with overall Aβ deposition in these mice (**Fig. 1**). The significant increase of GFAP in the cortex of SwDI/TKO mice was confirmed by immunoblot analysis of cortical homogenates from SwDI/TKO and SwDI/TWT mice (**Fig. 5H-I**). Notably, there was no difference in GFAP immunoreactivity between SwDI/TWT and SwDI/TKO in thalamus (**Fig. 5D, G**), where CAA and microgliosis were significantly reduced in SwDI/TKO (**Fig. 3** and **Fig. 4**). These results suggest that astrocyte activation is independent of TREM2 and responds more to parenchymal Aβ deposition than to CAA.

**Fig. 5.**
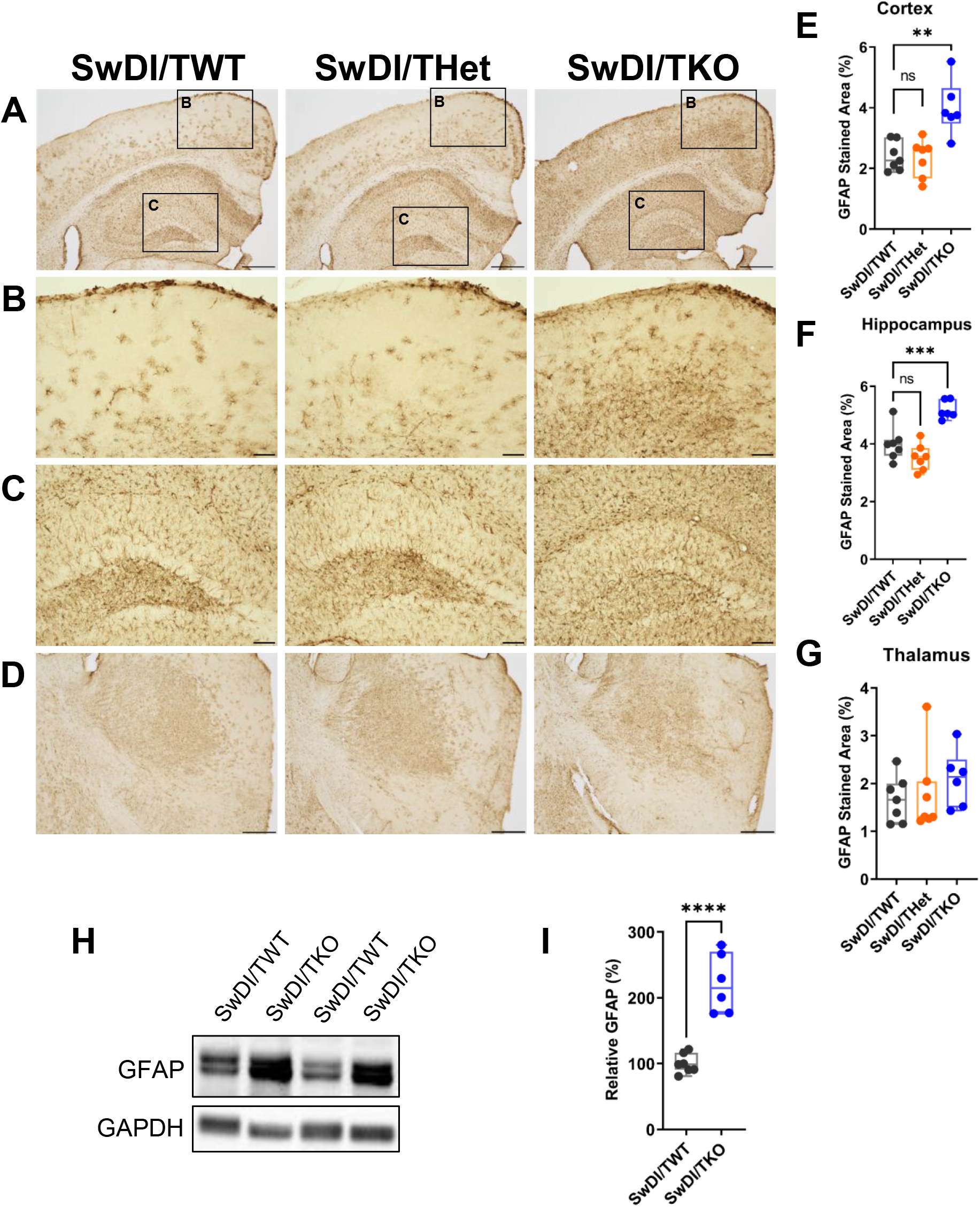
*Trem2* deletion aggravates cortical and hippocampal astrogliosis. **A, D** Representative images of the immunohistochemical staining of reactive astrocytes by GFAP in (**A**) cortex, hippocampus, and (**D**) thalamus in SwDI/TWT (n=7), SwDI/THet (n=7), and SwDI/TKO (n=6). Scale bars, 500 μm. **B-C** Selected zoomed-in areas from the representative images of (**B**) cortex and (**C**) hippocampus. Scale bars, 100 μm. **E-G** Quantifications of the immunoreactive area of GFAP in SwDI/TWT, SwDI/THet, and SwDI/TKO in (**E**) cortical, (**F**) hippocampal, and (**G**) thalamic regions, respectively. One-way ANOVA and Tukey’s post-hoc test. **H-I** Immunoblot analysis of GFAP. Representative images on the GFAP immnoblot (**H**) and the quantification of GFAP normalized by GAPDH (**I**). Unpaired student’s t-test, two-tail. **p<0.01, ***p<0.001, ****p<0.0001. ns, not significant.

### Single nucleus transcriptomic analysis reveals that microglia are trapped in transition in the absence of TREM2

To further investigate the molecular mechanisms underlying the pathological changes following *Trem2* deletion, snRNA-seq was performed using the frontal cortex tissues of SwDI/TWT and SwDI/TKO mice. After removing low-quality nuclei, transcriptional data from 36,660 nuclei were subjected to unsupervised clustering t-distributed stochastic neighbor embedding (t-SNE) in two dimensions. To avoid manual annotation of cell types, bootstrapping algorithms were applied to match cells in our dataset to mouse cortical cell types in a published dataset (Tasic *et al*., 2016) to predict the cell types, namely oligodendrocyte precursor cells (OPCs), vascular cells, oligodendrocytes (oligo), inhibitory neurons, excitatory neurons, microglia, astrocytes, and macrophages (**Fig. 6A-D**). The marker genes generated from these cell types were compared with reported marker genes (McKenzie *et al*., 2018; Zeisel *et al*., 2018; Grubman *et al*., 2019; Ximerakis *et al*., 2019) to confirm the cell type prediction (**Fig. 6B-C**). For both genotypes, although there were some individual variations in relative fractions of cell types, overall there were no significant differences in representation of different cell types between SwDI/TWT and SwDI/TKO (**Fig. 6E-F**).

**Fig. 6.**
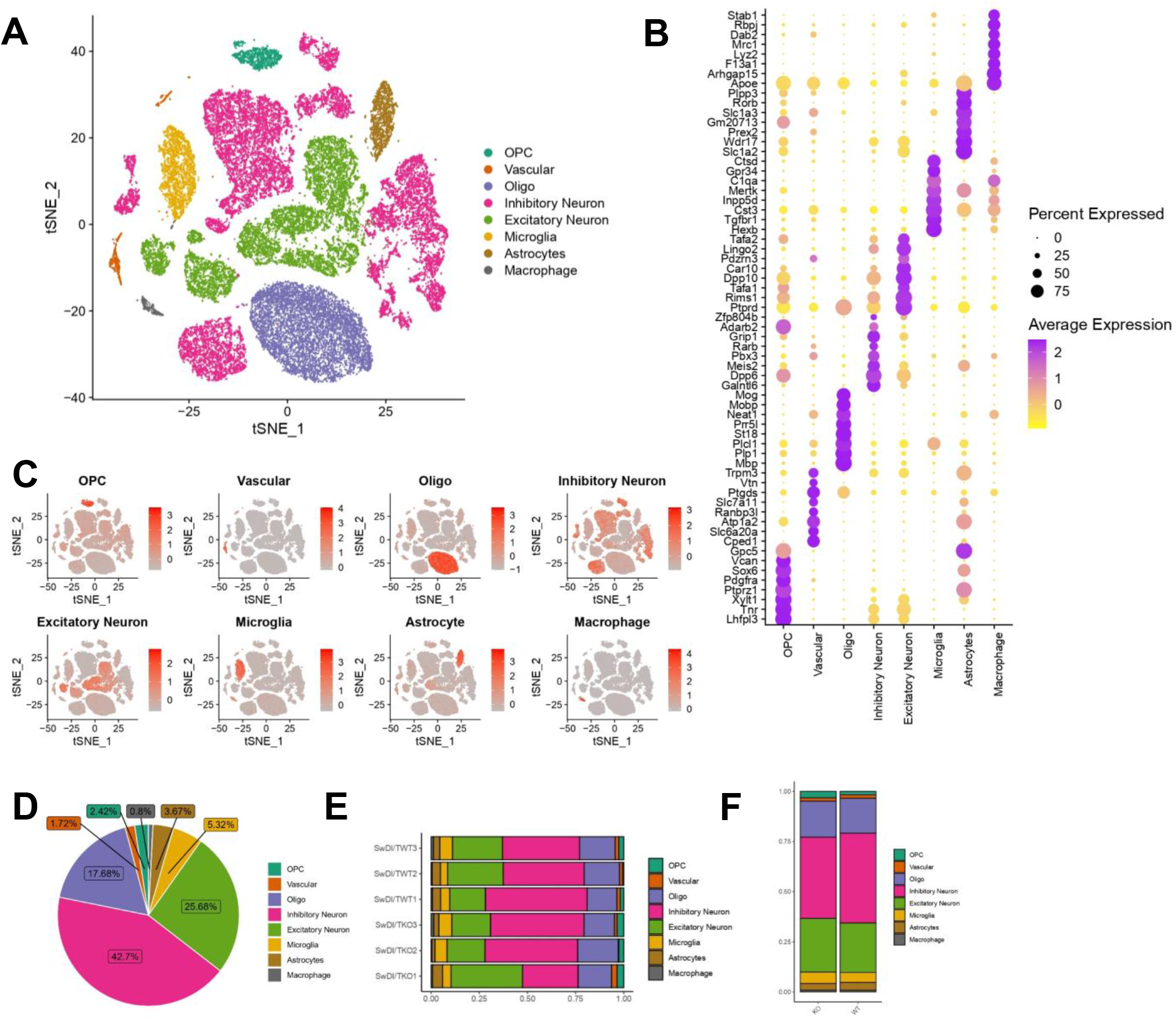
snRNA-seq analysis distinguishes major brain-cell types in SwDI/*Trem2* mouse brain cortical samples (n=6 total with n=3 mice for each genotype; 36660 total nuclei). **A** t-SNE plot showing distinguished clusters after integrating dataset from individual samples with eight distinct cell-type identities as determined by algorithm prediction and manual confirmation. **B** Expression of specific markers in every predicted cell type. **C** t-SNE visualization of all eight major cell populations showing the average expression of the representative cell type-specific marker genes. Numbers reflect the average number of UMI detected for the representative genes for each cell. **D** Pie chart showing the fraction of each cell type after integrating all six samples for both genotypes. **E-F** Bar graph showing the fractions of each cell type in **(E)** individual samples and **(F)** each genotype averaged as a group.

Multiple AD studies have used *Trem2*-deficient models and demonstrated distinct reactive transcriptomic profiles in microglia that are *Trem2*-dependent, featuring genes present in disease associated microglia (DAM). Therefore, the analysis was focused first on the microglia cluster (**Fig. 7A**) comparing SwDI/TKO to SwDI/TWT, and 71 differentially expressed genes (DEGs) were identified, of which 58 are upregulated in SwDI/TKO (**Fig. 7B; Table S2**). Among the DEGs, known disease-associated microglia (DAM) genes, including *Ctsd*, *Ctsb*, *Gnas*, and *Apoe,* were upregulated in the SwDI/TKO group, while homeostatic microglial marker genes such as *Cx3cr1* were downregulated (Keren-Shaul *et al*., 2017; Krasemann *et al*., 2017; Zhou *et al*., 2020). Notably, these DEGs have been reported to be responsible for the microglial transition from homeostatic stage to exclusively Stage 1 DAM (also known as the intermediate stage), which is *Trem2*-independent (Keren-Shaul *et al*., 2017). Correspondingly, there were no DEGs involved in the *Trem2*-dependent transition to Stage 2 DAM (**Table S2**). Pathway analysis on microglia unveiled an overwhelming upregulation of pathways mostly related to a reactive state of microglia in SwDI/TKO compared to SwDI/TWT, including protein synthesis, cell death, energy production and mitochondrial regulation, immune response, autophagy, and lipid biosynthesis and metabolism (**Fig. 7C**). These findings indicate that microglia lacking Trem2 adopted a stressed state in response to amyloidosis. However, as microglia were trapped in the intermediate Stage 1 of the DAM transition in the absence of TREM2, they were not sufficiently reactive to contain Aβ deposition, leading to exacerbated amyloid pathology in SwDI/TKO mice.

**Fig. 7.**
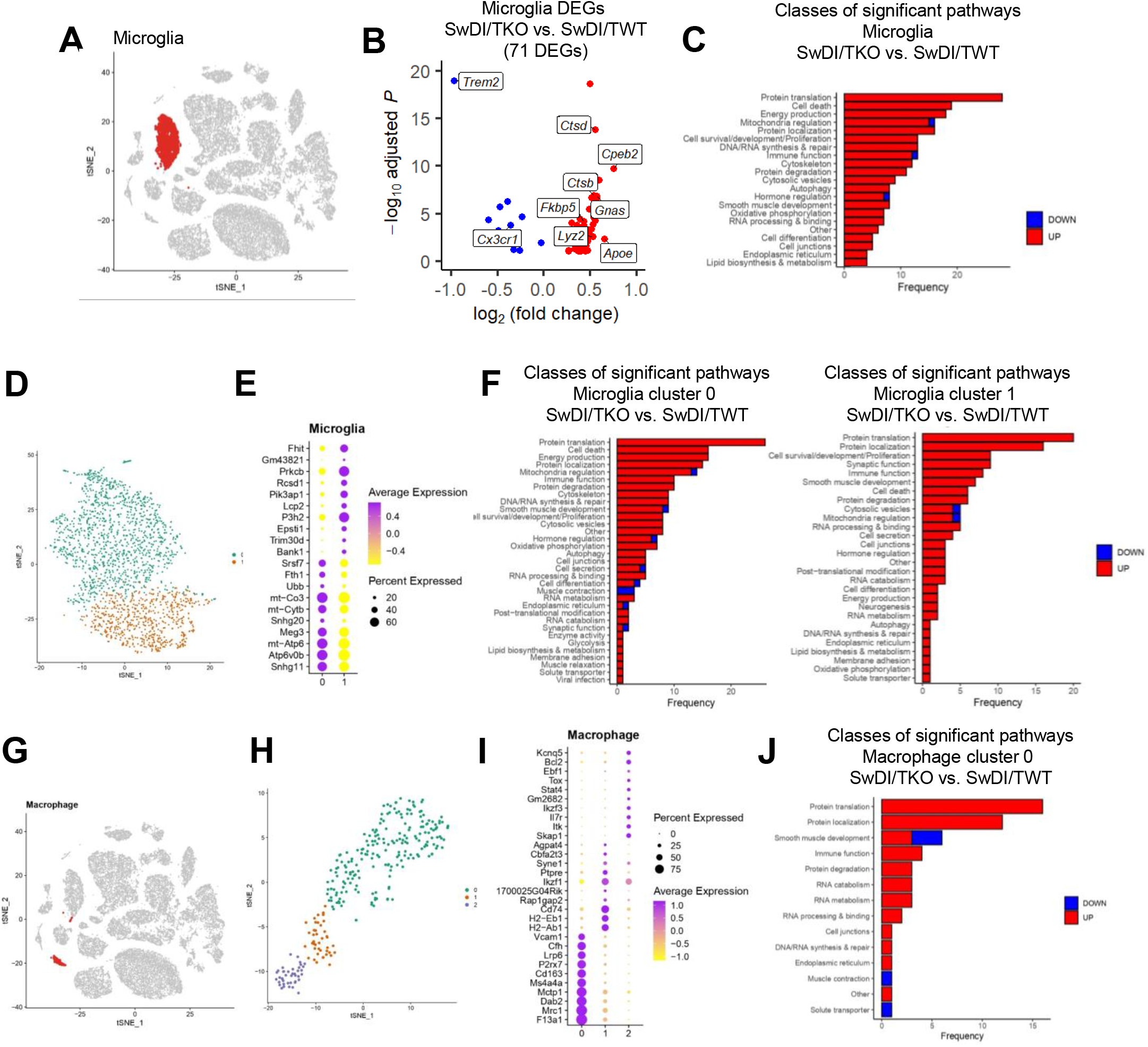
*Trem*2 deletion switches microglia and perivascular macrophages to a reactive transcriptomic profile. **A** t-SNE plot showing the microglia populations from **Fig. 6** based on the cell type prediction algorithm. **B** Volcano plots showing all 71 significant DEGs (adjusted *p* < 0.1) in microglia of SwDI/TKO vs. SwDI/TWT. Stage 1 DAM genes are elevated in SwDI/TKO. **C** Bar graph showing different classes of significant pathways (adjusted *p* < 0.1) of SwDI/TKO vs. SwDI/TWT in all microglia. Up-regulated pathways in red and down-regulated pathways in blue. **D** t-SNE plot of re-clustered microglia identifying two sub-clusters, 0 and 1. **E** Expression of specific markers in each microglial cluster. **F** Bar graphs showing different classes of significant pathways (adjusted *p* < 0.1) of SwDI/TKO vs. SwDI/TWT in microglia clusters 0 and 1. Up-regulated pathways in red and down-regulated pathways in blue. **G** t-SNE plot showing the macrophage populations from **Fig. 6** based on the cell type prediction algorithm. **H** t-SNE plot of re-clustered macrophage identifying three sub-clusters, 0, 1, and 2. **I** Expression of specific markers in each macrophage cluster. **J** Bar graph showing different classes of significant pathways (adjusted *p* < 0.1) of SwDI/TKO vs. SwDI/TWT in macrophage cluster 0. Up-regulated pathways in red and down-regulated pathways in blue.

Interestingly, further re-clustering of microglia uncovered two subclusters (0 and 1) (**Fig. 7D-E**), although the distribution of microglia between the two clusters was similar in SwDI/TWT and SwDI/TKO (**Fig. S4A**). Pathway analysis showed that the two subclusters of microglia have distinct functions. In microglia cluster 0, protein synthesis, mitochondria functions, and energy metabolism were among the most significantly enriched Gene Ontology (GO) pathways, while microglia cluster 1 was characterized mainly by immune functions including immune cell activation, phagocytosis, and cytokine production (**Fig. S4B**). Compared to SwDI/TWT, *Trem2* deficiency enhanced the respective signature functions characterizing the microglial subclusters 0 and 1, demonstrating divergent contributions from microglia clusters 0 and 1 to the stressed microglial state in SwDI/TKO (**Fig. 7F**).

### Perivascular macrophages are differentially activated in the absence of TREM2

In addition to microglia, TREM2 is also expressed in macrophages (Li *et al*., 2022). To understand the impact of TREM2 deletion on macrophages in the context of amyloidosis in SwDI mice, further analysis was conducted for a small population of macrophages identified from cell type prediction (**Fig. 7G**). The macrophages were re-clustered into three sub-populations (0, 1, and 2) (**Fig. 7H**). Cluster 0 covered the largest proportion of cells, characterized by perivascular macrophage (PVM) marker CD163 and Mrc1 (CD206) (Fabriek *et al*., 2005; Kim *et al*., 2006; Holder *et al*., 2014; Yang, Guo and Zhang, 2019) (**Fig. 7I**). PVMs are a distinct population of resident brain macrophages characterized by close association with the cerebral vasculature. In AD, these vessel-associated macrophages are shown to exacerbate AD pathologies including CAA and neuroinflammation and contribute to the detrimental effect of Aβ in affecting cerebral blood flow (Thanopoulou *et al*., 2010; Park *et al*., 2017). There were two significant DEGs in SwDI/TKO vs. SwDI/TWT: *Fkbp5*, crucial in AKT pathway regulation and NF-κB activation (Wang, 2011; Zannas *et al*., 2019), was up-regulated; whereas *Malat1*, a non-coding RNA, was downregulated, as observed in microglia carrying the dysfunctional *Trem2*-R47H mutation (Sayed *et al*., 2021). Pathway analysis of cluster 0 (PVM) revealed that pathways related to protein synthesis and immune reaction are more enriched in SwDI/TKO (**Fig. 7J**). In addition, several pathways pertaining to cell junction maintenance were upregulated in the absence of *Trem2* (**Fig. 7J**). Cluster 1 macrophages are featured by high expression of classic MHC class II genes such as Cd74, H2-Eb1, and H2-Ab1 (**Fig. 7I**), indicating an immune reactive status. Although no significant DEGs or pathways were found in SwDI/TKO vs. SwDI/TWT, there was a trend of increase in cluster 1 macrophages in SwDI/TKO (**Fig. S4C-D**). Together, these findings showed that perivascular macrophages were differentially activated in the absence of TREM2, likely contributing to pathological changes in SwDI/TKO mice.

### Vascular cell-type analysis uncovers distinct responses of mural cells and astrocytes to *Trem2* deficiency

To investigate the impact of *Trem*2 deletion on cerebrovascular cells, the snRNA-seq dataset was further analyzed to identify DEGs in vascular cell types (**Fig. 8A**). Comparing SwDI/TKO vs. SwDI/TWT mice, five upregulated and six downregulated protein-coding DEGs were identified (**Fig. 8B; Table S2**), and pathway analysis indicated their crucial roles in cytoskeleton maintenance (**Fig. 8C**). Furthermore, using recently published subtype-specific vascular marker genes (Vanlandewijck *et al*., 2018; Barisano *et al*., 2022; Lee *et al*., 2022), distinct cell clusters were identified, including smooth muscle cells (SMC), vascular leptomeningeal cells, pericytes, endothelial cells, and astrocytes (**Fig. 8D-E**). Pathway analysis of SwDI/TKO vs. SwDI/TWT unveiled divergent responses of vascular cells to *Trem2* deletion. In vascular leptomeningeal and endothelial cells, no significant changes were detected. For SMC, pericytes, and astrocytes, their respective significant pathways (adjusted *p*-value < 0.1) were shown in **Fig. 8F**. In both SMC and pericytes, an overall increased cell activity was observed in SwDI/TKO, including cytoskeleton, cell survival/development/proliferation, and immune function pathways. In addition, in SMC, pathways relevant to smooth muscle contraction were upregulated, and the extracellular matrix pathway was downregulated. As increased microvascular extracellular matrix has been associated with CAA (Damodarasamy *et al*., 2020), downregulation of the extracellular matrix was consistent with reduced CAA in SwDI/TKO. Interestingly, while the majority of significant pathways identified in SMC and pericytes were upregulated, almost all significant pathways in vascular-associated astrocytes were downregulated in SwDI/TKO mice. These findings indicate that various types of cerebrovascular cells exhibit differential responses to *Trem2* deletion, likely contributing to the pathological phenotypes in SwDI/TKO mice.

**Fig. 8.**
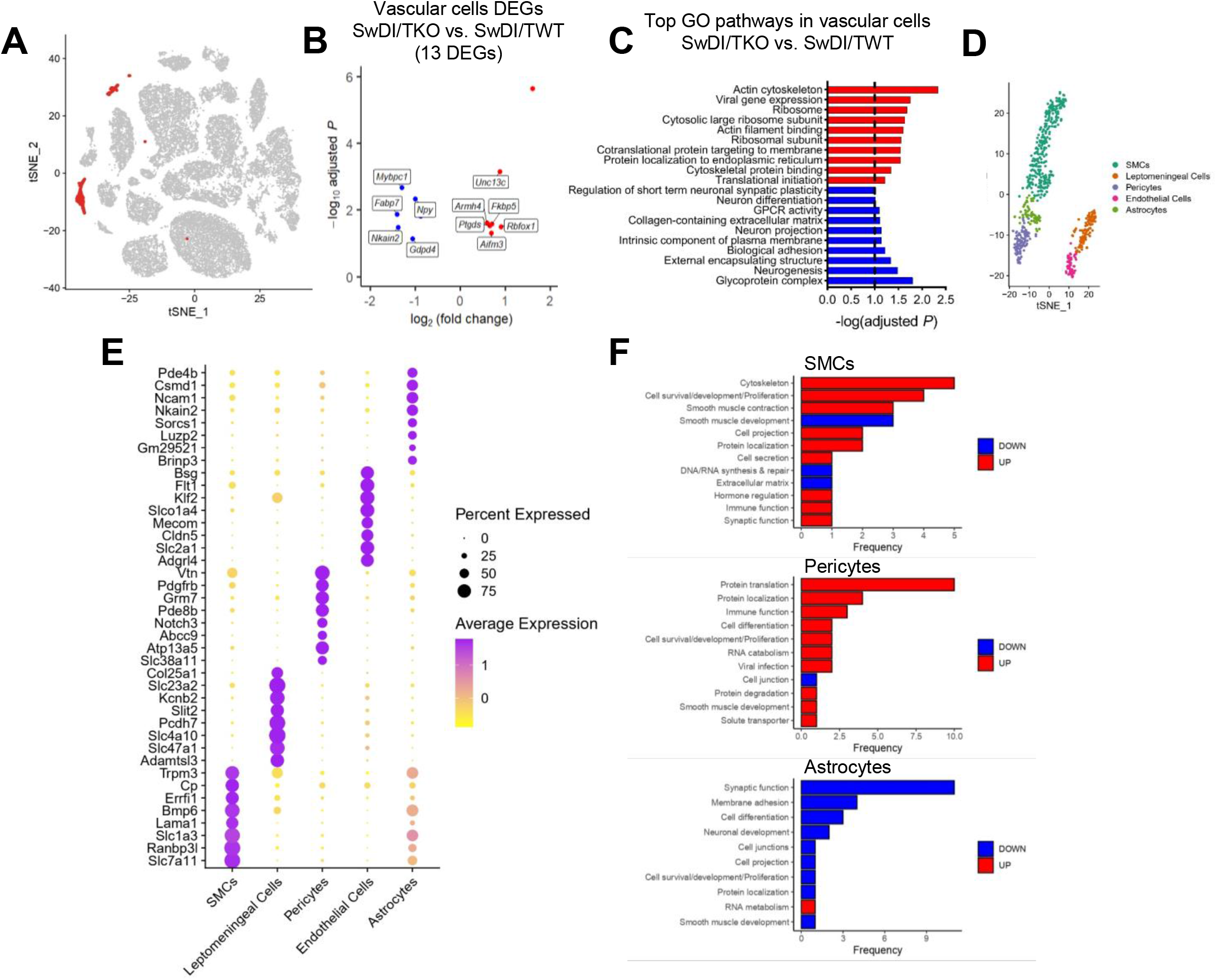
Vascular cell-type analysis uncovers distinct responses of mural cells and astrocytes to *Trem2* deficiency. **A** t-SNE plot showing the vascular cell populations from **Fig. 6** based on the cell type prediction algorithm. **B** Volcano plots showing all 13 significant DEGs (adjusted *p* < 0.1) in vascular cells of SwDI/TKO vs. SwDI/TWT. **C** Bar graph showing top 10 up- and down-regulated Gene Ontology (GO) significant pathways (adjusted *p* < 0.1) of SwDI/TKO vs. SwDI/TWT in all vascular cells. Up-regulated pathways in red and down-regulated pathways in blue. **D** t-SNE plot of re-clustered vascular cells showing five distinct cell types manually identified by reported marker genes. **E** Expression of specific markers in each vascular subtype. **F** Bar graph showing different classes of significant pathways (adjusted *p* < 0.1) of SwDI/TKO vs. SwDI/TWT in smooth muscle cells (SMC), pericytes, and astrocytes. No significant differential pathways were found in vascular leptomeningeal cells and endothelial cells.

## Discussion

The present study aimed at filling the gap in knowledge of the role of TREM2 in the pathogenesis of CAA. To achieve this goal, we introduced *Trem2*-deficiency on the background of SwDI, an established CAA/AD model (Davis *et al*., 2004; Miao *et al*., 2005), by breeding SwDI mice with the well-characterized *Trem2* KO mice (Turnbull *et al*., 2006). We performed a comprehensive characterization of the pathologies and molecular signatures of the new SwDI/Trem2 line using immunohistochemical, biochemical, and transcriptomic approaches, and we identified a previously unknown differential effect of TREM2 on modulating parenchymal and vascular amyloid pathology (plaques and CAA). Prior studies have shown that *Trem2* deficiency affects overall amyloid pathology in a disease stage-dependent manner, generally with an increase in total amyloid in the late stage but a decrease or no change in the earlier phase (Ulrich *et al*., 2014; Jay *et al*., 2015, 2017; Wang *et al*., 2015, 2016; Krasemann *et al*., 2017; Sheng *et al*., 2019). However, those studies were conducted in Aβ42-enriched, parenchymal plaques-dominant mouse models. In the present study, we used the Aβ40-enriched, CAA-prone SwDI/Trem2 mice at 16 months, when both amyloid plaques and CAA are usually well developed in SwDI mice at this age (Miao *et al*., 2005), to assess the function of TREM2.

Consistent with previous findings from other AD models at a late disease stage, loss of TREM2 led to a robust increase in overall amyloid load in the cortex, hippocampus, and thalamus in SwDI mice (**Fig. 1**). Interestingly, haplodeficiency of *Trem2* was not sufficient to modulate cerebral Aβ deposition in SwDI mice, which is consistent with the results in APPPS1-21 mice (Ulrich *et al*., 2014), although a dose-dependent effect of *Trem2* has been reported in 5XFAD mice (Wang *et al*., 2015). Because of the fibrillar nature of Aβ in CAA, in addition to Aβ immunostaining, methoxy-X04 staining was used to assess the amount of amyloid fibrils in SwDI/TREM2 mice. The results showed a significant increase of fibrillar Aβ in the three regions in SwDI/TKO, similar to total amyloid load assessed by Aβ immunostaining; however, unexpectedly, loss of TREM2 led to marked reduction of CAA, in particular in the thalamus where CAA is predominantly located in SwDI mice (**Fig. 3**). These intriguing findings, for the first time, reveal the differential impact of TREM2 deficiency on parenchymal and vascular amyloidosis.

Subsequent quantification of Aβ species was performed with Aβ40 and Aβ42 specific ELISA to understand their contributions to the discordant parenchymal Aβ deposition and CAA. The results showed a significant increase in both Aβ40 and Aβ42 in either soluble or insoluble fractions of the total brain homogenates in SwDI/TKO mice. Importantly, the Aβ40/Aβ42 ratio was also elevated, indicating that Aβ40 was increased more than Aβ42. Since the increase of Aβ40 promotes CAA, this would predict exacerbation rather than diminishment of CAA as observed in SwDI/TKO mice. This paradoxical finding might be the result of Aβ redistribution. As reported in a study with a bigenic model from crossing SwDI with 5XFAD, an aggressive plaque-rich model (Xu *et al*., 2014), early parenchymal fibrillar amyloid plaques originated from 5XFAD act as a scaffold to capture CAA composed by vasculotropic Dutch/Iowa mutant Aβ and promote its local assembly and deposition into parenchymal plaques, hence precluding microvascular amyloid formation. Based on the finding that loss of or dysfunctional TREM2 boosts amyloid seeding in the early stages (Parhizkar *et al*., 2019; Zhao *et al*., 2022), the shift from CAA to parenchymal plaques was likely caused by increased amyloid seeding in the brain parenchyma of SwDI/TKO mice. The overall increase of both total and fibrillar Aβ deposition in various brain regions observed in SwDI mice supports this notion. Interestingly, recent studies found that depletion of microglia led to a decrease in parenchymal plaques but an increase in CAA in 5XFAD mice (Spangenberg *et al*., 2019; Shabestari *et al*., 2022). These findings corroborate the role of microglia in the dynamics of parenchymal and vascular Aβ deposition, and in the meantime demonstrate that lack of microglia is not equivalent to lack of TREM2 and other microglial functions are likely responsible for the opposite outcome. In addition, the intrinsic differences between the mouse models (Aβ40-dominnat SwDI vs Aβ42-dominant 5XFAD) and the disease stages may also contribute to the discrepancy. How depletion of microglia modulates the parenchymal and vascular Aβ distribution in SwDI mice awaits further investigation.

In SwDI/TKO mice, CAA-associated IBA1+ microglia were reduced compared to SwDI/TWT, consistent with previous studies reporting reduced plaque-associated microglia in other *Trem2*-deficient AD mice (Ulrich *et al*., 2014; Jay *et al*., 2015, 2017; Wang *et al*., 2015, 2016; Krasemann *et al*., 2017; Parhizkar *et al*., 2019; Meilandt *et al*., 2020). Notably, in the brain cortex where diffused plaques were prevalent, there was no change in the extent of microgliosis by IBA1 staining (**Fig. 4D**) in the absence of TREM2. This could be attributed to the fact that the microglial activation in the cortex of SwDI mice is limited in the first place due to the scarce Aβ fibrillar pathology (Miao *et al*., 2005; Fan *et al*., 2007; Xu *et al*., 2007). Although *Trem2* deletion led to an increase in fibrillar Aβ, *Trem2*-deficient microglia failed to respond in SwDI/TKO mice, which is supported by snRNA-seq data.

Transcriptomic analysis showed that microglia displayed a partially reactive profile, trapped in the intermediate state of DAM Stage 1 in SwDI/TKO compared to SwDI/TWT, without being fully activated to the DAM Stage 2 as reported (Keren-Shaul *et al*., 2017; Krasemann *et al*., 2017; McQuade *et al*., 2020; Zhou *et al*., 2020; Zhao *et al*., 2022). These results are consistent with the notion that Stage 1 DAM genes are *Trem2* independent whereas Stage 2 DAM genes are *Trem2* dependent (Keren-Shaul *et al*., 2017). Pathway analysis also revealed a stressed state of microglia in SwDI/TKO, with overwhelming upregulation of protein synthesis, cell death, energy production, immune response, and lipid biosynthesis and metabolism pathways. Interestingly, it has been reported that *Trem2* deficiency leads to induced autophagy but curtails biosynthetic and energetic metabolism in 5XFAD mice (Ulland *et al*., 2017). Here, we found both autophagy and energy production pathways were upregulated in SwDI/TKO mice, indicating both overlapping and divergent responses in different AD models.

Further, we identified a PVM sub-cluster from the macrophage population through snRNA-seq analysis. The role of perivascular macrophages in cerebrovascular function is well recognized. In the homeostatic state, PVMs in the brain are beneficial, involving in the clearance of waste products from the cerebral parenchyma and in the regulation of the CSF flow through extracellular matrix (Kida *et al*., 1993; Drieu *et al*., 2022). However, under pathological conditions including AD, PVMs could be a double-edged sword, depending on the disease stage. On one hand, PVMs facilitate amyloid clearance at the vascular level (Hawkes and McLaurin, 2009; Thanopoulou *et al*., 2010). On the other hand, PVMs exacerbate Aβ-induced neurovascular dysfunction by providing reactive oxygen species(Park *et al*., 2017). In addition, PVMs contribute to the degradation of extracellular matrix that in turn regulates the diameter of the perivascular space and increases CSF flow (Ahn *et al*., 2019; Drieu *et al*., 2022).

Pathway analysis showed that PVMs were differentially activated in SwDI/TKO mice. In addition to upregulation of protein synthesis and immune function pathways as in microglia, cell junction pathways were upregulated in the absence of *Trem2* (**Fig. 7J**), indicating a potential impact on BBB integrity. In addition, snRNA-seq analyses in vascular cells showed that mural cells (smooth muscle cells and pericytes) were activated, whereas vascular-associated astrocytes were suppressed in SwDI/TKO mice. Since these cells are involved in the regulation of neurovascular structure and functions (Takano *et al*., 2005; Hill *et al*., 2015; Sweeney, Ayyadurai and Zlokovic, 2016), transcriptomic changes in each of the cell types likely contributed to the pathological phenotype in SwDI/TKO mice. The results also demonstrate that lack of TREM2 not only affects microglia but also modifies the response of various other cells types in the brain.

A limitation of the present study was the use of only one aged cohort of mice at the late stage of the disease. Thus, it remains unknown whether Trem2 deficiency affects amyloid pathology in SwDI mice in the age/stage-dependent manner as observed in other AD models (Ulrich *et al*., 2014; Jay *et al*., 2015, 2017; Wang *et al*., 2015, 2016; Krasemann *et al*., 2017; Sheng *et al*., 2019; Joshi *et al*., 2021). Another limitation was the focus on the pathological changes without functional assessments. Therefore, it is not clear whether the reduction of CAA, despite the increase of parenchymal Aβ deposition, leads to functional improvement in SwDI/TKO mice. Future studies including different age groups, along with behavioral and cerebrovascular function measurements, are required to address these questions.

Overall, our study provides the first evidence that TREM2 differentially modulates parenchymal and cerebrovascular amyloid pathology in a CAA-prone mouse model of AD. Lack of Trem2 exacerbates overall amyloid deposition but diminish CAA, the well-recognized culprit for cerebrovascular dysfunction, including the severe side effect of ARIA in anti-Aβ immunotherapy. Therefore, our findings may have significant implications for both TREM2- and Aβ-targeting therapies for AD.

## Acknowledgments

We thank Dr. Marco Colonna at Washington University for providing the original breeding pairs of Trem2-/- mice, Andrea Gram for maintaining and genotyping the experimental mice, and Stephen Martin for assisting in immunofluorescence imaging.

## Author contributions

RZ performed or participated in all experiments, analyzed the data, interpreted the results, and wrote the manuscript. YX performed the snRNA-seq data analysis and wrote relevant sections of the manuscript. JWW provided guidance on snRNA-seq data analysis and assisted data interpretation. LL conceived the study, supervised the progress of all experiments, and edited and finalized the manuscript. All authors reviewed and approved the manuscript.

