## supplemental material for "Loss of TREM2 exacerbates parenchymal amyloid pathology but diminishes CAA in Tg-SwDI mice"

### Supplementary Materials

**Fig. S1.** *Trem2* deletion exacerbates plaque deposition of various sizes in the cortex of SwDI mice.

**Fig. S2.** *Trem2* deletion increases amyloid fibril counts but decreases its density in the thalamus.

**Fig. S3.** *Trem2* deletion reduces microglial clustering in the CA1 stratum oriens region of SwDI mice.

**Fig. S4.** Functional analysis of microglia and macrophage subclusters.

**Table S1.** Classification of significant pathways

**Table S2.** DEGs in microglia and vascular cells, SwDI/TKO vs. SwDI/TWT

### Fig. S1

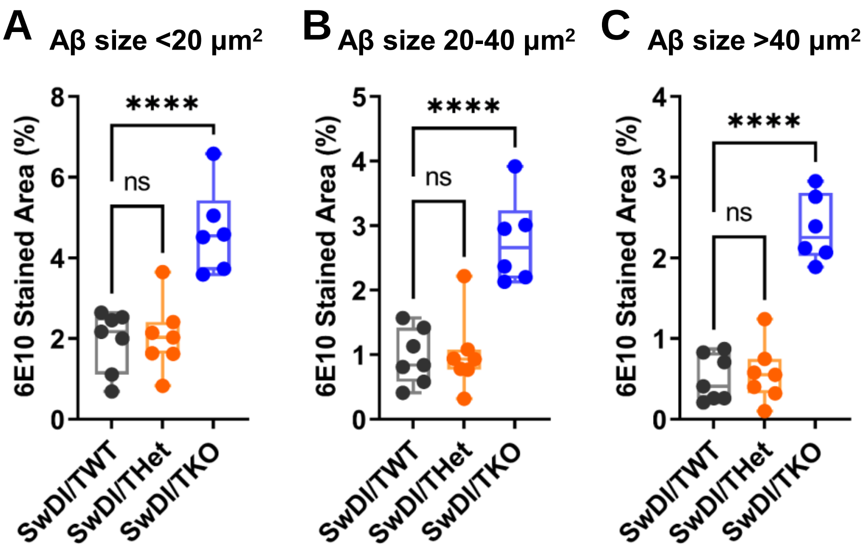

**Fig. S1.** *Trem2* deletion exacerbates plaque deposition of various sizes in the cortex of SwDI mice. **A-C** Quantification of cortical 6E10+ amyloid plaques in Fig. 1 in the sizes of **(A)** < 20 μm$^2$, **(B)** 20-40 μm$^2$, and **(C)** > 40 μm$^2$. One-way ANOVA and Tukey's post-hoc test. n=6-7/genotype. ****$p<0.0001$. ns, not significant.

### Fig. S2

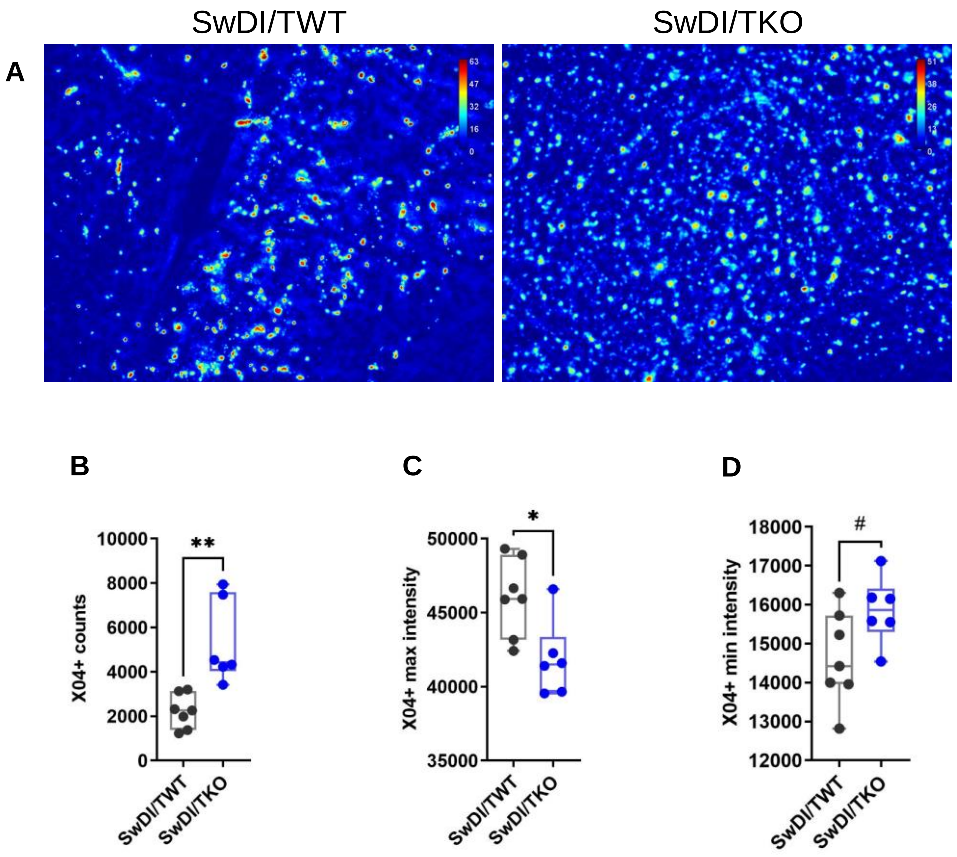

**Fig. S2.** *Trem2* deletion increases amyloid fibril counts but decreases its density in the thalamus. **A** Representative fluorescent intensity heatmap of $X04^+$ amyloid fibrils in the thalamus of SwDI/TWT and SwDI/TKO. The colored scale shows the gray values of the fluorescent staining. **B** Number of $X04^+$ staining in the thalamus. SwDI/TKO has significantly more $X04^+$ staining than SwDI/TWT. **C-D** Quantification of the **(C)** maximum, and **(D)** minimum fluorescent intensity of $X04^+$ fibrils based on individual puncta in SwDI/TWT and SwDI/TKO. n=6-7/genotype. #p<0.1, *p<0.05, **p<0.01.

### Fig. S3

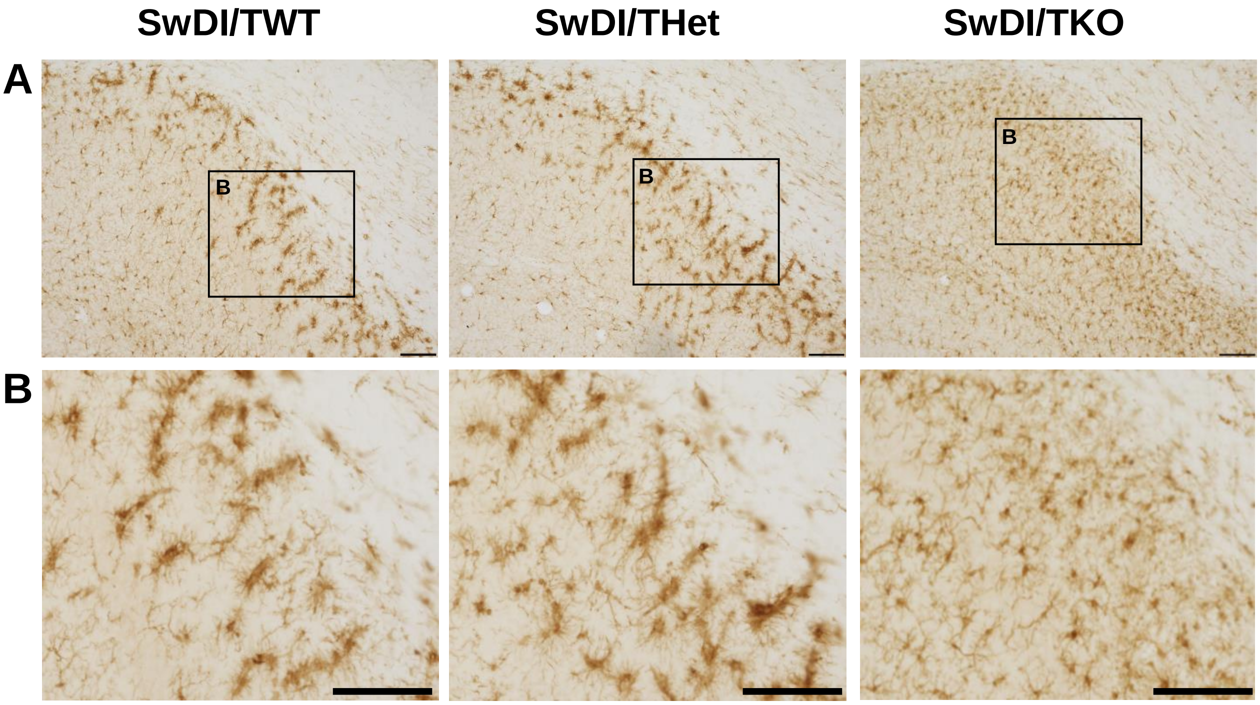

**Fig. S3.** *Trem2* deletion reduces microglial clustering in the CA1 stratum oriens region of SwDI mice. **A** Representative images of IBA stained brain sections focusing on the CA1 stratum oriens region. **B** Selected zoomed-in area from (**A**). Scale bars, 100 μm.

### Fig. S4

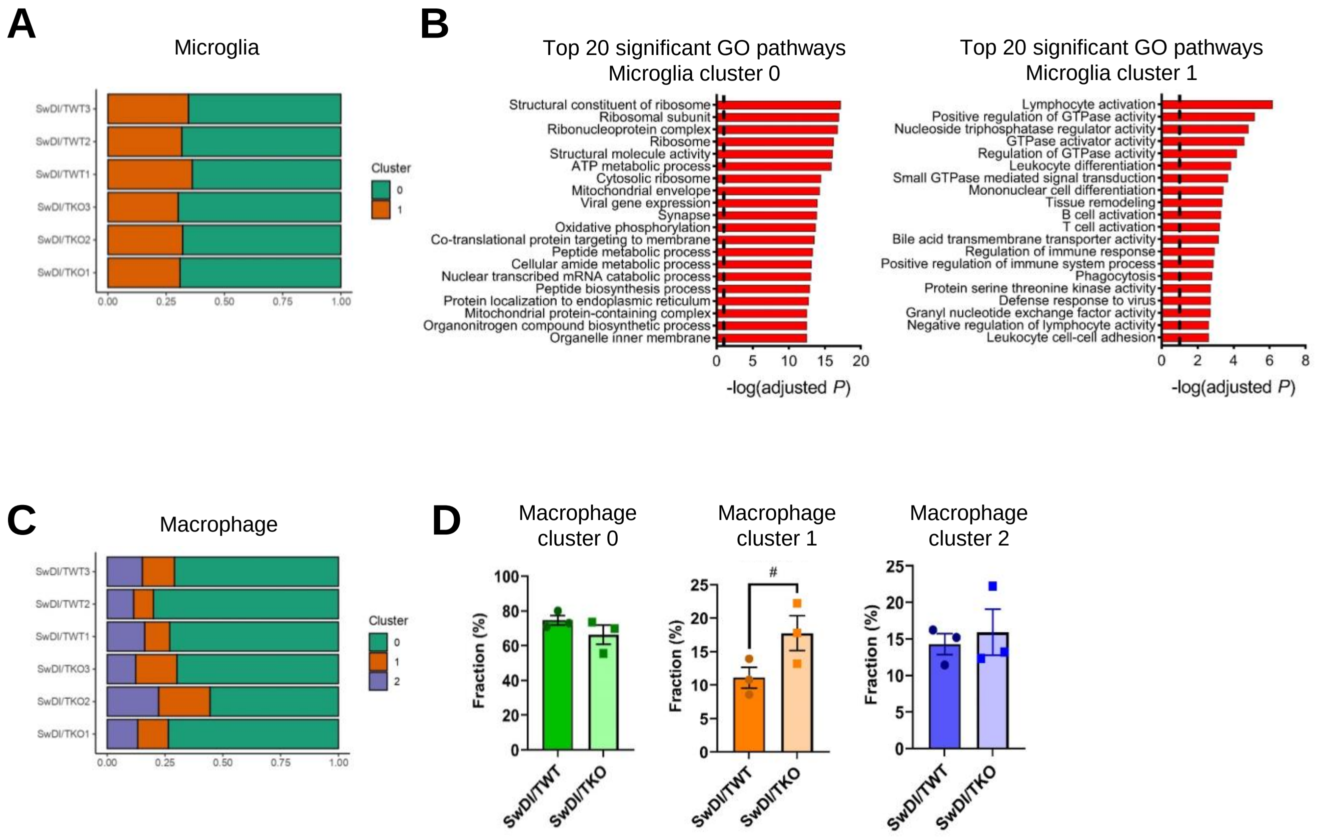

**Fig. S4.** Functional analysis of microglia and macrophage subclusters. **A** Bar graph showing the fractions of both microglia clusters 0 and 1 in individual samples. Both clusters are similarly represented among the total six mice analyzed. **B** Bar graphs showing the top 20 significant Gene Ontology (GO) signature pathways (adjusted *p* < 0.1) from cluster-specific DEGs in clusters 0 (left) and 1 (right). Dashed lines, adjusted p=0.1. **C** Bar graph showing the fractions of macrophage clusters 0, 1, and 2 in individual samples. **D** Macrophage cluster fractions comparing SwDI/TWT and SwDI/TKO groups. n=3 per genotype. #p<0.1.

### Table S1. Classification of significant pathways

#### Microglia.0

| pathway | pval | padj | log2err | ES | NES | size | dir | cell | class |
|---|---|---|---|---|---|---|---|---|---|
| GOMF_HYDROLASE_ACTIVITY_ACTING_ON_ACID_ANHYDRIDES | 9.38572E-05 | 0.005779917 | 0.538434 | 0.373666 | 1.340962 | 827 | UP | micro0 | Enzyme activity |
| GOBP_CYTOSKELETON_DEPENDENT_INTRACELLULAR_TRANSPORT | 0.000681723 | 0.02569922 | 0.477271 | 0.471689 | 1.513691 | 188 | UP | micro0 | Cytoskeleton |
| GOCC_MICROTUBULE_CYTOSKELETON | 2.82504E-05 | 0.002218035 | 0.57561 | 0.365113 | 1.335443 | 1201 | UP | micro0 | Cytoskeleton |
| GOBP_MICROTUBULE_BASED_MOVEMENT | 0.002668399 | 0.067289192 | 0.431708 | 0.410549 | 1.391506 | 335 | UP | micro0 | Cytoskeleton |
| GOBP_MICROTUBULE_BASED_PROCESS | 0.001267135 | 0.039877618 | 0.45506 | 0.35829 | 1.28509 | 787 | UP | micro0 | Cytoskeleton |
| GOBP_MICROTUBULE_BASED_TRANSPORT | 0.001403266 | 0.042759209 | 0.45506 | 0.449752 | 1.43546 | 181 | UP | micro0 | Cytoskeleton |
| GOBP_TRANSPORT_ALONG_MICROTUBULE | 0.001250184 | 0.039551177 | 0.45506 | 0.482027 | 1.510796 | 151 | UP | micro0 | Cytoskeleton |
| GOCC_MICROTUBULE | 0.000370109 | 0.016568583 | 0.498493 | 0.416113 | 1.432689 | 399 | UP | micro0 | Cytoskeleton |
| GOMF_BETA_TUBULIN_BINDING | 0.000172096 | 0.00930212 | 0.518848 | 0.710229 | 1.845906 | 38 | UP | micro0 | Cytoskeleton |
| GOMF_TUBULIN_BINDING | 0.001075182 | 0.035731264 | 0.45506 | 0.414262 | 1.414446 | 352 | UP | micro0 | Cytoskeleton |
| GOCC_SECRETORY_GRANULE | 1.12608E-08 | 2.06463E-06 | 0.74774 | 0.438978 | 1.571715 | 732 | UP | micro0 | Cell secretion |
| GOCC_SECRETORY_VESICLE | 1.16271E-07 | 1.79549E-05 | 0.704976 | 0.412948 | 1.48957 | 879 | UP | micro0 | Cell secretion |
| REACTOME_SENESCENCE_ASSOCIATED_SECRETORY_PHENOTYPE_SASP | 0.003008097 | 0.07276483 | 0.431708 | 0.58109 | 1.599861 | 52 | UP | micro0 | Cell secretion |
| GOBP_ARACHIDONIC_ACID_SECRETION | 0.002706157 | 0.067798901 | 0.431708 | -0.688149 | -1.68412 | 25 | DOWN | micro0 | Cell secretion |
| GOBP_SECRETION | 0.000530218 | 0.021467528 | 0.477271 | 0.341282 | 1.253549 | 1330 | UP | micro0 | Cell secretion |
| GOBP_VACUOLE_ORGANIZATION | 0.001061182 | 0.035338793 | 0.45506 | 0.471925 | 1.491656 | 166 | UP | micro0 | Cytosolic vesicles |
| GOCC_VACUOLE | 1.2553E-06 | 0.000150027 | 0.643552 | 0.412685 | 1.477356 | 738 | UP | micro0 | Cytosolic vesicles |
| GOBP_SYNAPTIC_VESICLE_LOCALIZATION | 0.004653144 | 0.095480228 | 0.407018 | 0.567311 | 1.561925 | 52 | UP | micro0 | Cytosolic vesicles |
| GOBP_SYNAPTIC_VESICLE_TRANSPORT | 0.000882182 | 0.030875423 | 0.477271 | 0.657492 | 1.727188 | 41 | UP | micro0 | Cytosolic vesicles |
| GOCC_VESICLE_LUMEN | 5.94427E-07 | 7.76581E-05 | 0.659444 | 0.500846 | 1.6713 | 289 | UP | micro0 | Cytosolic vesicles |
| GOCC_VESICLE_MEMBRANE | 0.000363384 | 0.016423042 | 0.498493 | 0.372705 | 1.33466 | 741 | UP | micro0 | Cytosolic vesicles |
| REACTOME_VESICLE_MEDIATED_TRANSPORT | 6.40476E-05 | 0.004217863 | 0.538434 | 0.393561 | 1.395103 | 635 | UP | micro0 | Cytosolic vesicles |
| GOCC_CLATHRIN_COMPLEX | 0.004094556 | 0.088854894 | 0.407018 | 0.940923 | 1.608203 | 6 | UP | micro0 | Cytosolic vesicles |
| GOBP_OXIDATIVE_PHOSPHORYLATION | 1.97885E-05 | 0.00162482 | 0.57561 | 0.557614 | 1.732064 | 137 | UP | micro0 | Oxidative phosphorylation |
| HALLMARK_OXIDATIVE_PHOSPHORYLATION | 6.4129E-09 | 1.22448E-06 | 0.761461 | 0.588915 | 1.902341 | 197 | UP | micro0 | Oxidative phosphorylation |
| KEGG_OXIDATIVE_PHOSPHORYLATION | 5.81264E-06 | 0.000549872 | 0.610527 | 0.58726 | 1.808256 | 121 | UP | micro0 | Oxidative phosphorylation |
| WP_OXIDATIVE_PHOSPHORYLATION | 0.002933662 | 0.071825749 | 0.431708 | 0.579236 | 1.615732 | 57 | UP | micro0 | Oxidative phosphorylation |
| REACTOME_THE_CITRIC_ACID_TCA_CYCLE_AND_RESPIRATORY_ELECTR | 2.02078E-06 | 0.000225635 | 0.627257 | 0.554041 | 1.757325 | 172 | UP | micro0 | Oxidative phosphorylation |
| WP_GLYCOLYSIS_AND_GLUCONEOGENESIS | 7.02279E-05 | 0.004523319 | 0.538434 | 0.710398 | 1.866169 | 41 | UP | micro0 | Glycolysis |
| CONCANNON_APOPTOSIS_BY_EPOXOMICIN_UP | 0.002502547 | 0.064552108 | 0.431708 | 0.433792 | 1.42363 | 231 | UP | micro0 | Cell death |
| GRAESSMANN_APOPTOSIS_BY_DOXORUBICIN_DN | 0.002675476 | 0.067291446 | 0.431708 | 0.320334 | 1.189096 | 1734 | UP | micro0 | Cell death |
| GRAESSMANN_APOPTOSIS_BY_DOXORUBICIN_UP | 0.000366113 | 0.016457084 | 0.498493 | 0.349377 | 1.274171 | 1107 | UP | micro0 | Cell death |
| GRAESSMANN_APOPTOSIS_BY_SERUM_DEPRIVATION_UP | 0.000294085 | 0.013976164 | 0.498493 | 0.395653 | 1.387492 | 538 | UP | micro0 | Cell death |
| HOLLMANN_APOPTOSIS_VIA_CD40_DN | 0.003595938 | 0.081947252 | 0.431708 | 0.413183 | 1.360928 | 243 | UP | micro0 | Cell death |
| FAN_OVARY_CL17_PUTATIVE_APOPTOTIC_SMOOTH_MUSCLE_CELL | 4.71017E-05 | 0.003268655 | 0.557332 | 0.49159 | 1.606382 | 222 | UP | micro0 | Cell death |
| FAN_OVARY_CL9_PUTATIVE_APOPTOTIC_ENDOTHELIAL_CELL | 1.2885E-05 | 0.001102878 | 0.593325 | 0.452635 | 1.544944 | 350 | UP | micro0 | Cell death |
| GOBP_APOPTOTIC_PROCESS | 1.70454E-07 | 2.57027E-05 | 0.690132 | 0.370413 | 1.374417 | 1673 | UP | micro0 | Cell death |
| GOBP_APOPTOTIC_SIGNALING_PATHWAY | 0.000281767 | 0.013541564 | 0.498493 | 0.395239 | 1.385985 | 535 | UP | micro0 | Cell death |
| GOBP_INTRINSIC_APOPTOTIC_SIGNALING_PATHWAY | 0.001671324 | 0.048369459 | 0.45506 | 0.424408 | 1.404698 | 264 | UP | micro0 | Cell death |
| GOBP_POSITIVE_REGULATION_OF_INTRINSIC_APOPTOTIC_SIGNALING_ | 0.003896521 | 0.086306046 | 0.431708 | 0.953448 | 1.563064 | 5 | UP | micro0 | Cell death |
| GOBP_REGULATION_OF_APOPTOTIC_SIGNALING_PATHWAY | 6.5491E-05 | 0.004269349 | 0.538434 | 0.442058 | 1.493058 | 321 | UP | micro0 | Cell death |
| GOBP_REGULATION_OF_INTRINSIC_APOPTOTIC_SIGNALING_PATHWAY | 0.001045478 | 0.035105663 | 0.45506 | 0.489259 | 1.529601 | 147 | UP | micro0 | Cell death |
| WP_SENESCENCE_AND_AUTOPHAGY_IN_CANCER | 0.002352656 | 0.062125897 | 0.431708 | 0.5068 | 1.508731 | 98 | UP | micro0 | Cell death |
| GOBP_NEGATIVE_REGULATION_OF_CELL_DEATH | 0.001903381 | 0.052948443 | 0.45506 | 0.349928 | 1.260855 | 856 | UP | micro0 | Cell death |
| GOBP_REGULATION_OF_CELL_DEATH | 3.01107E-05 | 0.002318952 | 0.57561 | 0.354279 | 1.305163 | 1428 | UP | micro0 | Cell death |
| GOBP_AUTOPHAGY_OF_MITOCHONDRION | 0.001096179 | 0.035984329 | 0.45506 | 0.570461 | 1.638806 | 72 | UP | micro0 | Autophagy |
| GOBP_MITOCHONDRION_ORGANIZATION | 1.11443E-06 | 0.000134687 | 0.643552 | 0.437862 | 1.532521 | 513 | UP | micro0 | Mitochondria regulation |
| GOCC_MITOCHONDRION | 0.000221963 | 0.011173966 | 0.518848 | 0.339509 | 1.253741 | 1524 | UP | micro0 | Mitochondria regulation |
| GOBP_ESTABLISHMENT_OF_PROTEIN_LOCALIZATION_TO_MITOCHOND | 0.003956767 | 0.087094754 | 0.407018 | 0.573285 | 1.578374 | 52 | UP | micro0 | Mitochondria regulation |
| GOBP_INNER_MITOCHONDRIAL_MEMBRANE_ORGANIZATION | 0.000219326 | 0.011082713 | 0.518848 | 0.651841 | 1.80751 | 54 | UP | micro0 | Mitochondria regulation |
| GOBP_MITOCHONDRIAL_MEMBRANE_ORGANIZATION | 0.000619059 | 0.024096985 | 0.477271 | 0.506553 | 1.575532 | 134 | UP | micro0 | Mitochondria regulation |
| GOBP_REGULATION_OF_RELEASE_OF_CYTOCHROME_C_FROM_MITOC | 0.000365864 | 0.016457084 | 0.498493 | 0.66468 | 1.780062 | 44 | UP | micro0 | Mitochondria regulation |
| GOBP_RELEASE_OF_CYTOCHROME_C_FROM_MITOCHONDRIA | 0.001348238 | 0.041741995 | 0.45506 | 0.614243 | 1.703255 | 54 | UP | micro0 | Mitochondria regulation |
| GOCC_INNER_MITOCHONDRIAL_MEMBRANE_PROTEIN_COMPLEX | 0.000128973 | 0.007366096 | 0.518848 | 0.532596 | 1.652297 | 133 | UP | micro0 | Mitochondria regulation |
| GOCC_INTRINSIC_COMPONENT_OF_MITOCHONDRIAL_INNER_MEMBR | 0.002378136 | 0.062526767 | 0.431708 | 0.588429 | 1.620069 | 52 | UP | micro0 | Mitochondria regulation |
| GOCC_MITOCHONDRIAL_ENVELOPE | 5.75162E-06 | 0.000549109 | 0.610527 | 0.400796 | 1.436132 | 735 | UP | micro0 | Mitochondria regulation |
| GOCC_MITOCHONDRIAL_PROTEIN_CONTAINING_COMPLEX | 0.000145505 | 0.008067549 | 0.518848 | 0.466603 | 1.536358 | 254 | UP | micro0 | Mitochondria regulation |
| HP_MITOCHONDRIAL_RESPIRATORY_CHAIN_DEFECTS | 0.003985744 | 0.087434385 | 0.407018 | -0.763594 | -1.728472 | 18 | DOWN | micro0 | Mitochondria regulation |
| MOOTHA_MITOCHONDRIA | 4.68187E-05 | 0.003263052 | 0.557332 | 0.426904 | 1.477015 | 431 | UP | micro0 | Mitochondria regulation |
| WONG_MITOCHONDRIA_GENE_MODULE | 7.64179E-05 | 0.004844655 | 0.538434 | 0.489914 | 1.594664 | 216 | UP | micro0 | Mitochondria regulation |
| WP_ELECTRON_TRANSPORT_CHAIN_OXPHOS_SYSTEM_IN_MITOCHON | 2.03724E-05 | 0.001664292 | 0.57561 | 0.601821 | 1.803544 | 100 | UP | micro0 | Oxidative phosphorylation |
| GOBP_CELL_GROWTH | 0.000675611 | 0.025528432 | 0.477271 | 0.397912 | 1.377637 | 444 | UP | micro0 | Cell survival/development/Proliferation |
| GOBP_DEVELOPMENTAL_CELL_GROWTH | 0.004856604 | 0.097887412 | 0.407018 | 0.420801 | 1.363662 | 208 | UP | micro0 | Cell survival/development/Proliferation |
| GOBP_CARDIAC_MUSCLE_MYOBLAST_PROLIFERATION | 0.00448361 | 0.093222695 | 0.407018 | -0.984796 | -1.440357 | 3 | DOWN | micro0 | Smooth muscle development |
| GOBP_CELL_POPULATION_PROLIFERATION | 0.001023865 | 0.034536331 | 0.45506 | 0.331692 | 1.228994 | 1644 | UP | micro0 | Cell survival/development/Proliferation |
| LEE_LIVER_CANCER_SURVIVAL_DN | 1.86895E-06 | 0.000210137 | 0.643552 | 0.548205 | 1.738492 | 169 | UP | micro0 | Cell survival/development/Proliferation |
| SHEDDEN_LUNG_CANCER_POOR_SURVIVAL_A6 | 0.00285082 | 0.070384925 | 0.431708 | 0.382386 | 1.326522 | 436 | UP | micro0 | Cell survival/development/Proliferation |
| BURTON_ADIPOGENESIS_6 | 0.000840647 | 0.030042767 | 0.477271 | 0.469307 | 1.500258 | 183 | UP | micro0 | Other |
| BURTON_ADIPOGENESIS_7 | 0.000159496 | 0.008708527 | 0.518848 | 0.684778 | 1.862784 | 49 | UP | micro0 | Other |
| GERHOLD_ADIPOGENESIS_UP | 0.000290106 | 0.013848286 | 0.498493 | 0.671835 | 1.80782 | 46 | UP | micro0 | Other |
| GOBP_CARDIAC_LEFT_VENTRICLE_MORPHOGENESIS | 0.004330179 | 0.091653159 | 0.407018 | 0.815771 | 1.643224 | 12 | UP | micro0 | Cell survival/development/Proliferation |
| GOBP_REGULATION_OF_MESENCHYMAL_TO_EPITHELIAL_TRANSITION_ | 0.004564477 | 0.0945993 | 0.407018 | 0.949275 | 1.556224 | 5 | UP | micro0 | Cell survival/development/Proliferation |
| GOBP_RIBONUCLEOPROTEIN_COMPLEX_BIOGENESIS | 2.55428E-07 | 3.72468E-05 | 0.674963 | 0.467916 | 1.617905 | 423 | UP | micro0 | Protein translation |
| GOBP_RIBOSOMAL_LARGE_SUBUNIT_BIOGENESIS | 8.3007E-05 | 0.005221348 | 0.538434 | 0.625346 | 1.786376 | 69 | UP | micro0 | Protein translation |
| GOBP_RIBOSOMAL_SMALL_SUBUNIT_BIOGENESIS | 0.00111151 | 0.036332288 | 0.45506 | 0.569982 | 1.637429 | 72 | UP | micro0 | Protein translation |
| GOBP_RIBOSOME_BIOGENESIS | 1.79296E-06 | 0.000203721 | 0.643552 | 0.489207 | 1.631431 | 292 | UP | micro0 | Protein translation |
| HP_RENAL_AGENESIS | 0.000641907 | 0.024688541 | 0.477271 | 0.489658 | 1.540859 | 160 | UP | micro0 | Other |
| REACTOME_GLUCONEOGENESIS | 0.00364338 | 0.082678086 | 0.431708 | 0.688132 | 1.698637 | 28 | UP | micro0 | Energy production |
| SCHLINGEMANN_SKIN_CARCINOGENESIS_TPA_UP | 0.001769894 | 0.050408393 | 0.45506 | 0.660798 | 1.703419 | 36 | UP | micro0 | Cell survival/development/Proliferation |
| WAKABAYASHI_ADIPOGENESIS_PPARG_RXRA_BOUND_8D | 0.001284216 | 0.040194352 | 0.45506 | 0.355239 | 1.275421 | 820 | UP | micro0 | Other |
| GOCC_AUTOPHAGOSOME | 0.000243165 | 0.012096751 | 0.518848 | 0.580131 | 1.713307 | 91 | UP | micro0 | Autophagy |
| GOBP_MACROAUTOPHAGY | 0.000432672 | 0.018263273 | 0.498493 | 0.4326 | 1.45218 | 300 | UP | micro0 | Autophagy |
| GOBP_REGULATION_OF_AUTOPHAGY | 0.003369299 | 0.077994202 | 0.431708 | 0.399673 | 1.349035 | 314 | UP | micro0 | Autophagy |
| REACTOME_AUTOPHAGY | 0.001976407 | 0.054278025 | 0.431708 | 0.48201 | 1.493861 | 140 | UP | micro0 | Autophagy |
| GOMF_CADHERIN_BINDING | 0.000940825 | 0.032437699 | 0.477271 | 0.418517 | 1.412692 | 313 | UP | micro0 | Cell junctions |
| GOCC_ANCHORING_JUNCTION | 3.14786E-05 | 0.00240138 | 0.557332 | 0.387166 | 1.387838 | 783 | UP | micro0 | Cell junctions |
| GOCC_CELL_SUBSTRATE_JUNCTION | 6.62947E-10 | 1.45528E-07 | 0.801216 | 0.501286 | 1.730509 | 416 | UP | micro0 | Cell junctions |
| REACTOME_GAP_JUNCTION_ASSEMBLY | 0.000734267 | 0.027140976 | 0.477271 | 0.742005 | 1.804091 | 25 | UP | micro0 | Cell junctions |
| REACTOME_GAP_JUNCTION_TRAFFICKING_AND_REGULATION | 0.000723461 | 0.026772203 | 0.477271 | 0.675009 | 1.744828 | 37 | UP | micro0 | Cell junctions |
| HALLMARK_CHOLESTEROL_HOMEOSTASIS | 0.001388863 | 0.042440363 | 0.45506 | 0.577243 | 1.666592 | 74 | UP | micro0 | Lipid biosynthesis & metabolism |
| GOBP_CELLULAR_MACROMOLECULE_LOCALIZATION | 1.23225E-12 | 4.14203E-10 | 0.91012 | 0.40103 | 1.493239 | 1849 | UP | micro0 | Protein localization |
| GOBP_ESTABLISHMENT_OF_ORGANELLE_LOCALIZATION | 0.000409847 | 0.017687067 | 0.498493 | 0.402083 | 1.386716 | 415 | UP | micro0 | Protein localization |

| | | | | | | | | | |
|---|---|---|---|---|---|---|---|---|---|
| GOBP_ESTABLISHMENT_OF_PROTEIN_LOCALIZATION | 2.34867E-14 | 9.24261E-12 | 0.975995 | 0.412202 | 1.534247 | 1838 | UP | micro0 | Protein localization |
| GOBP_ESTABLISHMENT_OF_PROTEIN_LOCALIZATION_TO_ENDOPLASM | 2.90631E-20 | 2.03878E-17 | 1.16018 | 0.810793 | 2.453225 | 110 | UP | micro0 | Protein localization |
| GOBP_ESTABLISHMENT_OF_PROTEIN_LOCALIZATION_TO_MEMBRANE | 2.39832E-16 | 1.12161E-13 | 1.037696 | 0.609127 | 2.058432 | 327 | UP | micro0 | Protein localization |
| GOBP_ESTABLISHMENT_OF_PROTEIN_LOCALIZATION_TO_ORGANELLE | 1.05365E-14 | 4.35902E-12 | 0.986546 | 0.527168 | 1.847911 | 536 | UP | micro0 | Protein localization |
| GOBP_ORGANELLE_LOCALIZATION | 0.000175853 | 0.009457641 | 0.518848 | 0.38187 | 1.353266 | 627 | UP | micro0 | Protein localization |
| GOBP_PROTEIN_LOCALIZATION_TO_ENDOPLASMIC_RETICULUM | 1.35835E-20 | 1.01936E-17 | 1.16907 | 0.776004 | 2.410427 | 137 | UP | micro0 | Protein localization |
| GOBP_PROTEIN_LOCALIZATION_TO_MEMBRANE | 6.30521E-12 | 1.93832E-09 | 0.887075 | 0.484895 | 1.717546 | 622 | UP | micro0 | Protein localization |
| GOBP_PROTEIN_LOCALIZATION_TO_ORGANELLE | 5.85815E-14 | 2.25044E-11 | 0.954542 | 0.457238 | 1.656569 | 941 | UP | micro0 | Protein localization |
| HP_ABNORMAL_LOCALIZATION_OF_KIDNEY | 0.003881804 | 0.086149875 | 0.431708 | 0.468218 | 1.475148 | 158 | UP | micro0 | Protein localization |
| GOBP_COTRANSLATIONAL_PROTEIN_TARGETING_TO_MEMBRANE | 1.0986E-22 | 1.47712E-19 | 1.229504 | 0.844056 | 2.512737 | 98 | UP | micro0 | Protein localization |
| GOBP_PROTEIN_TARGETING | 3.51087E-17 | 1.74296E-14 | 1.06721 | 0.570671 | 1.970033 | 416 | UP | micro0 | Protein localization |
| GOBP_PROTEIN_TARGETING_TO_MEMBRANE | 2.00691E-15 | 8.75149E-13 | 1.007318 | 0.67917 | 2.189435 | 190 | UP | micro0 | Protein localization |
| REACTOME_SRP_DEPENDENT_COTRANSLATIONAL_PROTEIN_TARGETIN | 2.33466E-22 | 2.717E-19 | 1.221054 | 0.838153 | 2.528964 | 107 | UP | micro0 | Protein localization |
| GOCC_ENDOPLASMIC_RETICULUM | 4.99468E-05 | 0.003421939 | 0.557332 | 0.343338 | 1.275732 | 1767 | UP | micro0 | Endoplasmic reticulum |
| GOCC_LUMENAL_SIDE_OF_ENDOPLASMIC_RETICULUM_MEMBRANE | 0.00430169 | 0.091377855 | 0.407018 | -0.687782 | -1.678982 | 24 | DOWN | micro0 | Endoplasmic reticulum |
| HELLER_HDAC_TARGETS_SILENCED_BY_METHYLATION_UP | 0.00164339 | 0.047603719 | 0.45506 | 0.395579 | 1.360209 | 397 | UP | micro0 | Post-translational modification |
| MISSIAGLIA_REGULATED_BY_METHYLATION_UP | 0.00086279 | 0.030591876 | 0.477271 | 0.528305 | 1.598497 | 110 | UP | micro0 | Post-translational modification |
| GOMF_PROTON_TRANSMEMBRANE_TRANSPORTER_ACTIVITY | 7.59487E-06 | 0.00070425 | 0.610527 | 0.601894 | 1.814035 | 108 | UP | micro0 | Solute transporter |
| GOBP_CELLULAR_MACROMOLECULE_CATABOLIC_PROCESS | 8.22374E-10 | 1.76608E-07 | 0.801216 | 0.414527 | 1.51277 | 1125 | UP | micro0 | Protein degradation |
| GOBP_LIPOPROTEIN_CATABOLIC_PROCESS | 0.004297498 | 0.091377855 | 0.407018 | 0.796944 | 1.701226 | 15 | UP | micro0 | Protein degradation |
| GOBP_MACROMOLECULE_CATABOLIC_PROCESS | 7.50326E-12 | 2.26283E-09 | 0.875325 | 0.415171 | 1.525562 | 1340 | UP | micro0 | Protein degradation |
| GOBP_NUCLEAR_TRANSCRIBED_MRNA_CATABOLIC_PROCESS | 1.68898E-14 | 6.81271E-12 | 0.975995 | 0.660019 | 2.132988 | 204 | UP | micro0 | RNA catabolism |
| GOBP_NUCLEAR_TRANSCRIBED_MRNA_CATABOLIC_PROCESS_NONSEN | 3.27519E-21 | 2.78124E-18 | 1.195344 | 0.801069 | 2.456729 | 118 | UP | micro0 | RNA catabolism |
| GOBP_ORGANIC_CYCLIC_COMPOUND_CATABOLIC_PROCESS | 1.16391E-11 | 3.35341E-09 | 0.875325 | 0.487963 | 1.725963 | 592 | UP | micro0 | Protein degradation |
| GOBP_ORGANONITROGEN_COMPOUND_CATABOLIC_PROCESS | 0.004430985 | 0.092486056 | 0.407018 | 0.328499 | 1.202147 | 1218 | UP | micro0 | Protein degradation |
| GOBP_PROTEIN_CATABOLIC_PROCESS | 0.002071832 | 0.056276055 | 0.431708 | 0.348512 | 1.256545 | 884 | UP | micro0 | Protein degradation |
| GOBP_REGULATION_OF_CATABOLIC_PROCESS | 0.000503984 | 0.020586152 | 0.477271 | 0.357616 | 1.29289 | 955 | UP | micro0 | Protein degradation |
| GOBP_REGULATION_OF_CELLULAR_CATABOLIC_PROCESS | 0.000219398 | 0.011082713 | 0.518848 | 0.368384 | 1.321881 | 806 | UP | micro0 | Protein degradation |
| GOBP_REGULATION_OF_PROTEIN_CATABOLIC_PROCESS | 0.003949377 | 0.086991421 | 0.407018 | 0.382489 | 1.308606 | 367 | UP | micro0 | Protein degradation |
| GOBP_RNA_CATABOLIC_PROCESS | 2.22261E-13 | 7.91064E-11 | 0.943632 | 0.546981 | 1.884528 | 403 | UP | micro0 | Protein degradation |
| GOMF_CELL_ADHESION_MOLECULE_BINDING | 0.003210723 | 0.07557025 | 0.431708 | 0.372775 | 1.305793 | 507 | UP | micro0 | Membrane adhesion |
| FAN_OVARY_CL14_MATURE_SMOOTH_MUSCLE_CELL | 1.01462E-08 | 1.8709E-06 | 0.74774 | 0.524558 | 1.753223 | 296 | UP | micro0 | Smooth muscle development |
| GOBP_CARDIAC_MUSCLE_CELL_ACTION_POTENTIAL_INVOLVED_IN_CC | 0.004313982 | 0.091403733 | 0.407018 | -0.575467 | -1.633367 | 49 | DOWN | micro0 | Muscle contraction |
| GOBP_MEMBRANE_REPOLARIZATION_DURING_ATRIAL_CARDIAC_MUS | 0.002082219 | 0.056510625 | 0.431708 | -0.949997 | -1.654167 | 6 | DOWN | micro0 | Muscle contraction |
| GOBP_REGULATION_OF_VENTRICULAR_CARDIAC_MUSCLE_CELL_ACTI( | 0.003232395 | 0.07596953 | 0.431708 | -0.81408 | -1.704418 | 12 | DOWN | micro0 | Muscle contraction |
| GOMF_VOLTAGE_GATED_POTASSIUM_CHANNEL_ACTIVITY_INVOLVED_ | 0.00397814 | 0.087330132 | 0.407018 | 0.805113 | 1.682546 | 14 | UP | micro0 | Muscle relaxation |
| RUBENSTEIN_SKELETAL_MUSCLE_B_CELLS | 2.57039E-20 | 1.8432E-17 | 1.16907 | 0.749571 | 2.367135 | 163 | UP | micro0 | Immune function |
| RUBENSTEIN_SKELETAL_MUSCLE_FAP_CELLS | 0.004809388 | 0.097483748 | 0.407018 | 0.432369 | 1.374637 | 173 | UP | micro0 | Smooth muscle development |
| RUBENSTEIN_SKELETAL_MUSCLE_FBN1_FAP_CELLS | 0.000112576 | 0.00659294 | 0.538434 | 0.457478 | 1.520544 | 276 | UP | micro0 | Smooth muscle development |
| RUBENSTEIN_SKELETAL_MUSCLE_MYELOID_CELLS | 1.79075E-12 | 5.8965E-10 | 0.91012 | 0.571571 | 1.931517 | 327 | UP | micro0 | Immune function |
| RUBENSTEIN_SKELETAL_MUSCLE_NK_CELLS | 1.87107E-11 | 5.16048E-09 | 0.863415 | 0.614873 | 1.991109 | 207 | UP | micro0 | Immune function |
| RUBENSTEIN_SKELETAL_MUSCLE_PCV_ENDOTHELIAL_CELLS | 1.24258E-09 | 2.5869E-07 | 0.788187 | 0.581651 | 1.892691 | 215 | UP | micro0 | Smooth muscle development |
| RUBENSTEIN_SKELETAL_MUSCLE_PERICYTES | 6.06317E-06 | 0.000568757 | 0.610527 | 0.571191 | 1.771401 | 138 | UP | micro0 | Smooth muscle development |
| RUBENSTEIN_SKELETAL_MUSCLE_SATELLITE_CELLS | 6.642E-29 | 3.06187E-25 | 1.395187 | 0.708934 | 2.37748 | 299 | UP | micro0 | Smooth muscle development |
| RUBENSTEIN_SKELETAL_MUSCLE_SMOOTH_MUSCLE_CELLS | 4.20731E-12 | 1.33104E-09 | 0.887075 | 0.529687 | 1.83387 | 428 | UP | micro0 | Smooth muscle development |
| RUBENSTEIN_SKELETAL_MUSCLE_T_CELLS | 5.21245E-26 | 1.20143E-22 | 1.318889 | 0.777815 | 2.466643 | 169 | UP | micro0 | Immune function |
| TRAVAGLINI_LUNG_AIRWAY_SMOOTH_MUSCLE_CELL | 1.82099E-09 | 3.62726E-07 | 0.788187 | 0.627892 | 1.972261 | 153 | UP | micro0 | Smooth muscle development |
| GOBP_CARDIAC_PACEMAKER_CELL_DIFFERENTIATION | 0.003865535 | 0.085906979 | 0.431708 | -0.954915 | -1.602308 | 5 | DOWN | micro0 | Cell differentiation |
| GOBP_MYELOID_CELL_DIFFERENTIATION | 0.003003577 | 0.072749267 | 0.431708 | 0.397889 | 1.357755 | 351 | UP | micro0 | Immune function |
| GOBP_OSTEOBLAST_DIFFERENTIATION | 0.002548962 | 0.065435525 | 0.431708 | 0.443012 | 1.433415 | 200 | UP | micro0 | Cell differentiation |
| GOBP_REGULATION_OF_EPITHELIAL_CELL_DIFFERENTIATION | 0.000261308 | 0.012763484 | 0.498493 | 0.536569 | 1.64628 | 117 | UP | micro0 | Cell differentiation |
| MATSUDA_NATURAL_KILLER_DIFFERENTIATION | 0.004055506 | 0.088423731 | 0.407018 | 0.375615 | 1.304676 | 460 | UP | micro0 | Immune function |
| MA_MYELOID_DIFFERENTIATION_UP | 3.93952E-05 | 0.002850325 | 0.557332 | 0.733646 | 1.891207 | 36 | UP | micro0 | Immune function |
| RIZ_ERYTHROID_DIFFERENTIATION | 0.001809222 | 0.050855206 | 0.45506 | 0.550608 | 1.597373 | 78 | UP | micro0 | Cell differentiation |
| GOCC_SYNAPSE | 0.002517182 | 0.064825976 | 0.431708 | 0.335869 | 1.229729 | 1228 | UP | micro0 | Synaptic function |
| REACTOME_PROTEIN_PROTEIN_INTERACTIONS_AT_SYNAPSES | 0.001470131 | 0.044164542 | 0.45506 | -0.524691 | -1.606866 | 85 | DOWN | micro0 | Synaptic function |
| REACTOME_HSP90_CHAPERONE_CYCLE_FOR_STEROID_HORMONE_RE( | 0.004589289 | 0.09487261 | 0.407018 | 0.600867 | 1.634522 | 49 | UP | micro0 | Hormone regulation |
| RHEIN_ALL_GLUCOCORTICOID_THERAPY_DN | 0.000190463 | 0.010032688 | 0.518848 | 0.434239 | 1.483068 | 358 | UP | micro0 | Hormone regulation |
| GOBP_CELLULAR_RESPONSE_TO_HORMONE_STIMULUS | 0.00201885 | 0.05506869 | 0.431708 | 0.369093 | 1.299979 | 554 | UP | micro0 | Hormone regulation |
| GOBP_RESPONSE_TO_HORMONE | 2.27567E-05 | 0.001826709 | 0.57561 | 0.387463 | 1.390423 | 789 | UP | micro0 | Hormone regulation |
| GOBP_RESPONSE_TO_PEPTIDE_HORMONE | 0.002254454 | 0.060222674 | 0.431708 | 0.391717 | 1.344763 | 396 | UP | micro0 | Hormone regulation |
| REACTOME_PEPTIDE_HORMONE_METABOLISM | 0.001227997 | 0.039117698 | 0.45506 | 0.567356 | 1.637663 | 75 | UP | micro0 | Hormone regulation |
| REACTOME_EICOSANOID_LIGAND_BINDING_RECEPTORS | 0.002470935 | 0.063941142 | 0.431708 | -0.851486 | -1.742129 | 11 | DOWN | micro0 | Hormone regulation |
| GOBP_ATP_METABOLIC_PROCESS | 3.85706E-06 | 0.000393872 | 0.610527 | 0.486014 | 1.620573 | 291 | UP | micro0 | Energy production |
| GOBP_ATP_SYNTHESIS_COUPLED_ELECTRON_TRANSPORT | 0.000784315 | 0.028565535 | 0.477271 | 0.541791 | 1.604614 | 94 | UP | micro0 | Energy production |
| GOCC_PROTON_TRANSPORTING_ATP_SYNTHASE_COMPLEX | 0.002320267 | 0.061471824 | 0.431708 | 0.767087 | 1.76708 | 20 | UP | micro0 | Energy production |
| GOCC_PROTON_TRANSPORTING_TWO_SECTOR_ATPASE_COMPLEX | 0.000282212 | 0.013541564 | 0.498493 | 0.672159 | 1.808692 | 46 | UP | micro0 | Energy production |
| GOMF_ATPASE_ACTIVITY | 0.000406377 | 0.017649235 | 0.498493 | 0.403921 | 1.400577 | 451 | UP | micro0 | Energy production |
| GOMF_PROTON_TRANSPORTING_ATP_SYNTHASE_ACTIVITY_ROTATION | 0.004367375 | 0.091991393 | 0.407018 | 0.796751 | 1.700815 | 15 | UP | micro0 | Energy production |
| REACTOME_FORMATION_OF_ATP_BY_CHEMIOSMOTIC_COUPLING | 0.003489357 | 0.080198041 | 0.431708 | 0.774886 | 1.70039 | 17 | UP | micro0 | Energy production |
| REACTOME_RESPIRATORY_ELECTRON_TRANSPORT_ATP_SYNTHESIS_B\ | 3.39323E-06 | 0.000353214 | 0.627257 | 0.581189 | 1.789773 | 124 | UP | micro0 | Energy production |
| BLANCO_MELO_RESPIRATORY_SYNCYTIAL_VIRUS_INFECTION_A594_CE | 0.001088089 | 0.035980125 | 0.45506 | 0.435339 | 1.435536 | 251 | UP | micro0 | Viral infection |
| DURANTE_ADULT_OLFACTORY_NEUROEPITHELIUM_RESPIRATORY_HOI | 0.004637999 | 0.095327128 | 0.407018 | 0.715134 | 1.68309 | 22 | UP | micro0 | Energy production |
| GOBP_RESPIRATORY_ELECTRON_TRANSPORT_CHAIN | 0.001498186 | 0.044763858 | 0.45506 | 0.513706 | 1.55989 | 111 | UP | micro0 | Energy production |
| GOCC_RESPIRATORY_CHAIN_COMPLEX | 0.004869787 | 0.097969543 | 0.407018 | 0.523521 | 1.518071 | 81 | UP | micro0 | Energy production |
| HP_ABNORMAL_RESPIRATORY_SYSTEM_MORPHOLOGY | 0.004215558 | 0.090446703 | 0.407018 | 0.338039 | 1.231569 | 1062 | UP | micro0 | Energy production |
| HP_ABNORMAL_RESPIRATORY_SYSTEM_PHYSIOLOGY | 0.004602036 | 0.09495084 | 0.407018 | 0.33003 | 1.202933 | 1140 | UP | micro0 | Energy production |
| REACTOME_RESPIRATORY_ELECTRON_TRANSPORT | 0.000576697 | 0.022979571 | 0.477271 | 0.543507 | 1.631985 | 101 | UP | micro0 | Energy production |
| GOBP_AEROBIC_RESPIRATION | 0.000246609 | 0.012223996 | 0.498493 | 0.585093 | 1.712573 | 83 | UP | micro0 | Oxidative phosphorylation |
| GOBP_CELLULAR_RESPIRATION | 1.14987E-05 | 0.00100014 | 0.593325 | 0.522883 | 1.662662 | 177 | UP | micro0 | Energy production |
| GOMF_DOUBLE_STRANDED_RNA_BINDING | 0.002386944 | 0.062614783 | 0.431708 | 0.549329 | 1.592459 | 77 | UP | micro0 | DNA/RNA synthesis & repair |
| GOBP_CELLULAR_RESPONSE_TO_DNA_DAMAGE_STIMULUS | 0.000507799 | 0.020715747 | 0.477271 | 0.363856 | 1.307533 | 821 | UP | micro0 | DNA/RNA synthesis & repair |
| GOBP_DNA_METABOLIC_PROCESS | 2.89714E-05 | 0.002252721 | 0.57561 | 0.378267 | 1.363711 | 870 | UP | micro0 | DNA/RNA synthesis & repair |
| GOBP_DNA_REPAIR | 0.000164751 | 0.008954126 | 0.518848 | 0.397395 | 1.393912 | 532 | UP | micro0 | DNA/RNA synthesis & repair |
| GOBP_REGULATION_OF_DNA_TEMPLATED_TRANSCRIPTION_IN_RESPC | 0.00369153 | 0.083243874 | 0.431708 | 0.488946 | 1.493536 | 115 | UP | micro0 | DNA/RNA synthesis & repair |
| KYNG_DNA_DAMAGE_DN | 0.001923662 | 0.053374577 | 0.45506 | 0.457205 | 1.451107 | 175 | UP | micro0 | DNA/RNA synthesis & repair |
| FERNANDEZ_BOUND_BY_MYC | 0.004579001 | 0.094778556 | 0.407018 | 0.434002 | 1.379829 | 173 | UP | micro0 | Other |
| GOBP_MRNA_METABOLIC_PROCESS | 3.35654E-10 | 7.57428E-08 | 0.814036 | 0.445966 | 1.600837 | 808 | UP | micro0 | RNA metabolism |
| GOBP_REGULATION_OF_TRANSCRIPTION_FROM_RNA_POLYMERASE_II | 0.003200777 | 0.075391148 | 0.431708 | 0.849099 | 1.668394 | 11 | UP | micro0 | DNA/RNA synthesis & repair |
| GOBP_RRNA_METABOLIC_PROCESS | 0.000644257 | 0.024721666 | 0.477271 | 0.448737 | 1.474796 | 229 | UP | micro0 | RNA metabolism |
| GOMF_MRNA_BINDING | 5.65276E-05 | 0.003792283 | 0.557332 | 0.465545 | 1.543921 | 272 | UP | micro0 | RNA processing & binding |
| GOMF_RNA_BINDING | 1.0888E-16 | 5.24393E-14 | 1.057464 | 0.435454 | 1.608604 | 1557 | UP | micro0 | RNA processing & binding |
| GOMF_RRNA_BINDING | 2.07419E-05 | 0.001690199 | 0.57561 | 0.672615 | 1.88541 | 61 | UP | micro0 | RNA processing & binding |

| | | | | | | | | | |
|---|---|---|---|---|---|---|---|---|---|
| GSE14000_TRANSLATED_RNA_VS_MRNA_DC_DN | 6.61455E-09 | 1.25556E-06 | 0.761461 | 0.59787 | 1.895993 | 169 | UP | micro0 | DNA/RNA synthesis & repair |
| GSE21360_NAIVE_VS_QUATERNARY_MEMORY_CD8_TCELL_DN | 0.000607028 | 0.023772079 | 0.477271 | 0.472227 | 1.522983 | 192 | UP | micro0 | Immune function |
| GSE21360_PRIMARY_VS_QUATERNARY_MEMORY_CD8_TCELL_UP | 9.66114E-05 | 0.005904455 | 0.538434 | 0.498956 | 1.576939 | 165 | UP | micro0 | Immune function |
| GSE32164_ALTERNATIVELY_ACT_M2_VS_CMYC_INHIBITED_MACROPHA | 0.003618066 | 0.082251957 | 0.431708 | 0.440667 | 1.421199 | 192 | UP | micro0 | Immune function |
| LI_DCP2_BOUND_MRNA | 0.002441898 | 0.063511731 | 0.431708 | 0.526275 | 1.551047 | 88 | UP | micro0 | Other |
| REACTOME_ACTIVATION_OF_THE_MRNA_UPON_BINDING_OF_THE_CA | 6.88245E-11 | 1.6825E-08 | 0.839089 | 0.815093 | 2.283262 | 59 | UP | micro0 | RNA processing & binding |
| REACTOME_CELLULAR_RESPONSES_TO_EXTERNAL_STIMULI | 1.11758E-20 | 8.58649E-18 | 1.177893 | 0.557731 | 1.973889 | 601 | UP | micro0 | Other |
| REACTOME_METABOLISM_OF_RNA | 8.30277E-12 | 2.43565E-09 | 0.875325 | 0.48388 | 1.719491 | 650 | UP | micro0 | RNA metabolism |
| REACTOME_RNA_POLYMERASE_II_TRANSCRIPTION | 0.000344009 | 0.015858324 | 0.498493 | 0.354197 | 1.291894 | 1116 | UP | micro0 | DNA/RNA synthesis & repair |
| REACTOME_RRNA_PROCESSING | 4.71194E-15 | 2.00065E-12 | 0.996986 | 0.676114 | 2.181127 | 195 | UP | micro0 | RNA processing & binding |
| BILANGES_SERUM_RESPONSE_TRANSLATION | 2.66343E-05 | 0.002105936 | 0.57561 | 0.766506 | 1.939681 | 32 | UP | micro0 | Protein translation |
| GOBP_CYTOPLASMIC_TRANSLATION | 7.54639E-11 | 1.83094E-08 | 0.839089 | 0.720537 | 2.142252 | 96 | UP | micro0 | Protein translation |
| GOBP_TRANSLATIONAL_ELONGATION | 0.003332009 | 0.077435968 | 0.431708 | 0.465671 | 1.441833 | 131 | UP | micro0 | Protein translation |
| GOBP_TRANSLATIONAL_INITIATION | 1.67E-18 | 1.01678E-15 | 1.114664 | 0.711985 | 2.272254 | 184 | UP | micro0 | Protein translation |
| GOMF_TRANSLATION_REGULATOR_ACTIVITY | 0.00415762 | 0.089680641 | 0.407018 | 0.462051 | 1.430626 | 131 | UP | micro0 | Protein translation |
| REACTOME_EUKARYOTIC_TRANSLATION_ELONGATION | 7.73684E-25 | 1.63096E-21 | 1.295123 | 0.881987 | 2.599405 | 88 | UP | micro0 | Protein translation |
| REACTOME_EUKARYOTIC_TRANSLATION_INITIATION | 2.10104E-22 | 2.60764E-19 | 1.229504 | 0.820644 | 2.506744 | 115 | UP | micro0 | Protein translation |
| REACTOME_TRANSLATION | 6.54793E-21 | 5.28238E-18 | 1.186651 | 0.660962 | 2.203737 | 286 | UP | micro0 | Protein translation |
| GOBP_RIBOSOME_ASSEMBLY | 5.82777E-06 | 0.000549872 | 0.610527 | 0.690301 | 1.929038 | 58 | UP | micro0 | Protein translation |
| GOCC_CYTOSOLIC_RIBOSOME | 6.58514E-24 | 1.06248E-20 | 1.270913 | 0.854064 | 2.542528 | 98 | UP | micro0 | Protein translation |
| GOCC_POLYSOMAL_RIBOSOME | 7.85178E-06 | 0.000724756 | 0.593325 | 0.800311 | 1.993555 | 30 | UP | micro0 | Protein translation |
| GOCC_RIBOSOME | 2.40744E-20 | 1.76558E-17 | 1.16907 | 0.703393 | 2.283366 | 211 | UP | micro0 | Protein translation |
| GOMF_RIBOSOME_BINDING | 0.004661112 | 0.095498042 | 0.407018 | 0.580496 | 1.614292 | 55 | UP | micro0 | Protein translation |
| GOMF_STRUCTURAL_CONSTITUENT_OF_RIBOSOME | 1.12859E-21 | 1.07113E-18 | 1.203975 | 0.769605 | 2.412138 | 151 | UP | micro0 | Protein translation |
| KEGG_RIBOSOME | 2.35755E-22 | 2.717E-19 | 1.221054 | 0.878868 | 2.572454 | 83 | UP | micro0 | Protein translation |
| GOBP_RIBOSOMAL_LARGE_SUBUNIT_ASSEMBLY | 0.003339813 | 0.077478374 | 0.431708 | 0.702435 | 1.72162 | 26 | UP | micro0 | Protein translation |
| GOCC_CYTOSOLIC_LARGE_RIBOSOMAL_SUBUNIT | 9.05626E-14 | 3.3981E-11 | 0.954542 | 0.86425 | 2.374046 | 51 | UP | micro0 | Protein translation |
| GOCC_CYTOSOLIC_SMALL_RIBOSOMAL_SUBUNIT | 9.67292E-11 | 2.32937E-08 | 0.839089 | 0.854864 | 2.277083 | 43 | UP | micro0 | Protein translation |
| GOCC_LARGE_RIBOSOMAL_SUBUNIT | 1.43222E-12 | 4.76457E-10 | 0.91012 | 0.727895 | 2.193787 | 108 | UP | micro0 | Protein translation |
| GOCC_RIBOSOMAL_SUBUNIT | 2.31504E-21 | 2.07511E-18 | 1.195344 | 0.73768 | 2.345675 | 177 | UP | micro0 | Protein translation |
| GOCC_SMALL_RIBOSOMAL_SUBUNIT | 1.60838E-09 | 3.28273E-07 | 0.788187 | 0.74904 | 2.151611 | 71 | UP | micro0 | Protein translation |
| WP_CYTOPLASMIC_RIBOSOMAL_PROTEINS | 2.86604E-21 | 2.49957E-18 | 1.195344 | 0.856374 | 2.517632 | 86 | UP | micro0 | Protein translation |

Microglia.1

| pathway | pval | padj | log2err | ES | NES | size | dir | cell | class |
|---|---|---|---|---|---|---|---|---|---|
| GOCC_SECRETORY_GRANULE | 9.37843E-10 | 3.51898E-07 | 0.788187 | 0.452173 | 1.606418 | 732 | UP | micro1 | Cell secretion |
| GOCC_SECRETORY_GRANULE_MEMBRANE | 5.66788E-05 | 0.005862074 | 0.557332 | 0.470892 | 1.57411 | 279 | UP | micro1 | Cell secretion |
| GOCC_SECRETORY_VESICLE | 3.11614E-09 | 1.03665E-06 | 0.774939 | 0.431739 | 1.547074 | 879 | UP | micro1 | Cell secretion |
| GOBP_SECRETION | 1.76435E-07 | 3.67315E-05 | 0.690132 | 0.383273 | 1.394608 | 1330 | UP | micro1 | Cell secretion |
| GOCC_VACUOLE | 0.000102324 | 0.009197489 | 0.538434 | 0.38885 | 1.382334 | 738 | UP | micro1 | Cytosolic vesicles |
| GOCC_ENDOCYTIC_VESICLE_LUMEN | 0.000321518 | 0.02202773 | 0.498493 | 0.858232 | 1.848514 | 16 | UP | micro1 | Cytosolic vesicles |
| GOCC_VESICLE_LUMEN | 6.03713E-05 | 0.006145491 | 0.557332 | 0.463439 | 1.553169 | 289 | UP | micro1 | Cytosolic vesicles |
| GOCC_VESICLE_MEMBRANE | 0.000145305 | 0.011870535 | 0.518848 | 0.38464 | 1.368394 | 741 | UP | micro1 | Cytosolic vesicles |
| GOCC_MULTIVESICULAR_BODY_LUMEN | 0.001261571 | 0.06121748 | 0.45506 | -0.992291 | -1.512113 | 4 | DOWN | micro1 | Cytosolic vesicles |
| HALLMARK_OXIDATIVE_PHOSPHORYLATION | 0.000133928 | 0.011138421 | 0.518848 | 0.492907 | 1.595252 | 197 | UP | micro1 | Oxidative phosphorylation |
| GRAESSMANN_APOPTOSIS_BY_DOXORUBICIN_UP | 7.25902E-07 | 0.00013234 | 0.659444 | 0.392907 | 1.416952 | 1107 | UP | micro1 | Cell death |
| GRAESSMANN_APOPTOSIS_BY_SERUM_DEPRIVATION_UP | 4.82887E-06 | 0.000721402 | 0.610527 | 0.430662 | 1.507047 | 538 | UP | micro1 | Cell death |
| HOLLMANN_APOPTOSIS_VIA_CD40_DN | 0.002713028 | 0.098699787 | 0.431708 | 0.432241 | 1.427946 | 243 | UP | micro1 | Cell death |
| GOBP_INTRINSIC_APOPTOTIC_SIGNALING_PATHWAY | 0.000758315 | 0.043081075 | 0.477271 | 0.439525 | 1.46303 | 264 | UP | micro1 | Cell death |
| GOBP_REGULATION_OF_INTRINSIC_APOPTOTIC_SIGNALING_P/ | 0.00159005 | 0.07146145 | 0.45506 | 0.488109 | 1.52854 | 147 | UP | micro1 | Cell death |
| WP_SENESCENCE_AND_AUTOPHAGY_IN_CANCER | 0.001659356 | 0.073250028 | 0.45506 | 0.532083 | 1.582207 | 98 | UP | micro1 | Cell death |
| GOBP_MITOCHONDRION_ORGANIZATION | 0.000875246 | 0.047388116 | 0.477271 | 0.385149 | 1.341237 | 513 | UP | micro1 | Mitochondria regulation |
| GOCC_MITOCHONDRION | 0.002159694 | 0.086572884 | 0.431708 | 0.331375 | 1.210257 | 1524 | UP | micro1 | Mitochondria regulation |
| GOCC_MITOCHONDRIAL_ENVELOPE | 0.000351567 | 0.023585707 | 0.498493 | 0.3742 | 1.329209 | 735 | UP | micro1 | Mitochondria regulation |
| HP_MITOCHONDRIAL_INHERITANCE | 0.001687305 | 0.07416557 | 0.45506 | -0.781498 | -1.767528 | 18 | DOWN | micro1 | Mitochondria regulation |
| MOOTHA_MITOCHONDRIA | 0.000230402 | 0.017250224 | 0.518848 | 0.41738 | 1.441543 | 431 | UP | micro1 | Mitochondria regulation |
| GOBP_CELL_GROWTH | 2.18445E-05 | 0.002610742 | 0.57561 | 0.433915 | 1.4986 | 444 | UP | micro1 | Cell survival/development/Proliferation |
| GOBP_DEVELOPMENTAL_CELL_GROWTH | 3.72272E-05 | 0.004081741 | 0.557332 | 0.504884 | 1.641825 | 208 | UP | micro1 | Cell survival/development/Proliferation |
| GOBP_REGULATION_OF_EXTENT_OF_CELL_GROWTH | 0.002287552 | 0.089583752 | 0.431708 | 0.511746 | 1.531586 | 102 | UP | micro1 | Cell survival/development/Proliferation |
| GOBP_NEGATIVE_REGULATION_OF_CELL_POPULATION_PROLI | 0.001693983 | 0.074270577 | 0.45506 | 0.370572 | 1.304343 | 613 | UP | micro1 | Cell survival/development/Proliferation |
| GERHOLD_ADIPOGENESIS_UP | 0.001008144 | 0.052983383 | 0.45506 | 0.640885 | 1.67682 | 46 | UP | micro1 | Other |
| GOBP_CELLULAR_COMPONENT_MORPHOGENESIS | 0.001440329 | 0.067067788 | 0.45506 | 0.36628 | 1.301214 | 726 | UP | micro1 | Cell survival/development/Proliferation |
| GOBP_CELL_MORPHOGENESIS | 0.002157758 | 0.086572884 | 0.431708 | 0.347879 | 1.249275 | 956 | UP | micro1 | Cell survival/development/Proliferation |
| GOBP_CELL_PART_MORPHOGENESIS | 0.001982 | 0.082328697 | 0.431708 | 0.369077 | 1.303431 | 651 | UP | micro1 | Cell survival/development/Proliferation |
| GOBP_CEREBELLAR_CORTEX_MORPHOGENESIS | 0.001564651 | 0.07065248 | 0.45506 | 0.698571 | 1.721441 | 32 | UP | micro1 | Neurogenesis |
| GOBP_DEVELOPMENTAL_GROWTH_INVOLVED_IN_MORPHOGE | 0.000120737 | 0.010529895 | 0.538434 | 0.482731 | 1.573847 | 216 | UP | micro1 | Cell survival/development/Proliferation |
| GOBP_NEUROGENESIS | 0.000201182 | 0.015493859 | 0.518848 | 0.347276 | 1.267841 | 1508 | UP | micro1 | Neurogenesis |
| GOBP_REGULATION_OF_MESENCHYMAL_TO_EPITHELIAL_TRAN | 1.00595E-06 | 0.000177382 | 0.643552 | 0.999744 | 1.596108 | 5 | UP | micro1 | Cell survival/development/Proliferation |
| GOBP_RIBOSOMAL_SMALL_SUBUNIT_BIOGENESIS | 0.001146005 | 0.05680558 | 0.45506 | 0.566168 | 1.610737 | 72 | UP | micro1 | Protein translation |
| GOBP_REGULATION_OF_AUTOPHAGY | 0.000964707 | 0.050949456 | 0.477271 | 0.422917 | 1.422148 | 314 | UP | micro1 | Autophagy |
| GOMF_CADHERIN_BINDING | 0.000180045 | 0.014135915 | 0.518848 | 0.448139 | 1.506563 | 313 | UP | micro1 | Cell junctions |
| GOCC_ANCHORING_JUNCTION | 3.42684E-05 | 0.003826326 | 0.557332 | 0.39165 | 1.39523 | 783 | UP | micro1 | Cell junctions |
| GOCC_CELL_SUBSTRATE_JUNCTION | 3.35589E-08 | 8.9497E-06 | 0.719513 | 0.486947 | 1.675947 | 416 | UP | micro1 | Cell junctions |
| GOBP_FATTY_ACID_TRANSPORT | 4.07491E-06 | 0.000620251 | 0.610527 | 0.564615 | 1.76453 | 145 | UP | micro1 | Lipid biosynthesis & metabolism |
| GOBP_CELLULAR_MACROMOLECULE_LOCALIZATION | 8.6363E-11 | 4.03891E-08 | 0.839089 | 0.390241 | 1.432838 | 1849 | UP | micro1 | Protein localization |
| GOBP_ESTABLISHMENT_OF_PROTEIN_LOCALIZATION | 1.5177E-13 | 1.2888E-10 | 0.943632 | 0.402666 | 1.478049 | 1838 | UP | micro1 | Protein localization |
| GOBP_ESTABLISHMENT_OF_PROTEIN_LOCALIZATION_TO_END | 2.57119E-16 | 4.88058E-13 | 1.037696 | 0.772537 | 2.338838 | 110 | UP | micro1 | Protein localization |
| GOBP_ESTABLISHMENT_OF_PROTEIN_LOCALIZATION_TO_MEN | 2.98471E-11 | 1.53869E-08 | 0.851339 | 0.556411 | 1.877628 | 327 | UP | micro1 | Protein localization |
| GOBP_ESTABLISHMENT_OF_PROTEIN_LOCALIZATION_TO_ORG | 7.60396E-13 | 5.22068E-10 | 0.921426 | 0.514959 | 1.802336 | 536 | UP | micro1 | Protein localization |
| GOBP_LIPID_LOCALIZATION | 0.001725102 | 0.074821671 | 0.45506 | 0.396405 | 1.369558 | 451 | UP | micro1 | Protein localization |
| GOBP_PROTEIN_LOCALIZATION_TO_ENDOPLASMIC_RETICULUI | 2.72046E-12 | 1.79156E-09 | 0.898671 | 0.689361 | 2.142609 | 137 | UP | micro1 | Protein localization |
| GOBP_PROTEIN_LOCALIZATION_TO_MEMBRANE | 3.43123E-10 | 1.41952E-07 | 0.814036 | 0.472735 | 1.666014 | 622 | UP | micro1 | Protein localization |
| GOBP_PROTEIN_LOCALIZATION_TO_ORGANELLE | 4.5778E-09 | 1.49213E-06 | 0.761461 | 0.424209 | 1.524126 | 941 | UP | micro1 | Protein localization |
| GOBP_REGULATION_OF_CELLULAR_LOCALIZATION | 0.00212523 | 0.086013412 | 0.431708 | 0.357769 | 1.273351 | 770 | UP | micro1 | Protein localization |
| GOBP_REGULATION_OF_LIPID_LOCALIZATION | 0.000415669 | 0.026456062 | 0.498493 | 0.502954 | 1.589526 | 158 | UP | micro1 | Protein localization |
| GOBP_REGULATION_OF_PROTEIN_LOCALIZATION | 0.000124904 | 0.010713471 | 0.518848 | 0.379782 | 1.354722 | 827 | UP | micro1 | Protein localization |
| GOBP_COTRANSLATIONAL_PROTEIN_TARGETING_TO_MEMBR/ | 9.90873E-17 | 2.66454E-13 | 1.057464 | 0.802086 | 2.385092 | 98 | UP | micro1 | Protein localization |
| GOBP_PROTEIN_TARGETING | 2.05202E-11 | 1.10361E-08 | 0.863415 | 0.529982 | 1.824064 | 416 | UP | micro1 | Protein localization |
| GOBP_PROTEIN_TARGETING_TO_MEMBRANE | 8.53624E-12 | 5.00829E-09 | 0.875325 | 0.638891 | 2.057295 | 190 | UP | micro1 | Protein localization |
| REACTOME_SRP_DEPENDENT_COTRANSLATIONAL_PROTEIN_T/ | 2.25901E-16 | 4.85974E-13 | 1.037696 | 0.790279 | 2.379505 | 107 | UP | micro1 | Protein localization |
| GOCC_ENDOPLASMIC_RETICULUM | 0.00012425 | 0.010713471 | 0.518848 | 0.341743 | 1.252985 | 1767 | UP | micro1 | Endoplasmic reticulum |
| KINNEY_DNMT1_METHYLATION_TARGETS | 0.001619396 | 0.072187116 | 0.45506 | 0.955914 | 1.673163 | 7 | UP | micro1 | Post-translational modification |
| LIANG_SILENCED_BY_METHYLATION_2 | 7.4733E-05 | 0.007113747 | 0.538434 | 0.736343 | 1.886185 | 40 | UP | micro1 | Post-translational modification |
| SATO_SILENCED_BY_METHYLATION_IN_PANCREATIC_CANCER_ | 0.001539871 | 0.069986068 | 0.45506 | 0.406435 | 1.384123 | 352 | UP | micro1 | Post-translational modification |
| GOMF_TRANSPORTER_ACTIVITY | 0.002010381 | 0.082957786 | 0.431708 | 0.345549 | 1.242736 | 1047 | UP | micro1 | Solute transporter |
| GOBP_CELLULAR_MACROMOLECULE_CATABOLIC_PROCESS | 1.40226E-07 | 3.09929E-05 | 0.690132 | 0.393308 | 1.419817 | 1125 | UP | micro1 | Protein degradation |
| GOBP_MACROMOLECULE_CATABOLIC_PROCESS | 2.79535E-09 | 9.39615E-07 | 0.774939 | 0.400299 | 1.456391 | 1340 | UP | micro1 | Protein degradation |
| GOBP_NUCLEAR_TRANSCRIBED_MRNA_CATABOLIC_PROCESS | 1.12924E-09 | 4.04882E-07 | 0.788187 | 0.595087 | 1.935539 | 204 | UP | micro1 | RNA catabolism |
| GOBP_NUCLEAR_TRANSCRIBED_MRNA_CATABOLIC_PROCESS_ | 1.15857E-13 | 1.0385E-10 | 0.943632 | 0.734708 | 2.243798 | 118 | UP | micro1 | RNA catabolism |
| GOBP_ORGANIC_CYCLIC_COMPOUND_CATABOLIC_PROCESS | 4.63453E-07 | 8.69487E-05 | 0.674963 | 0.43602 | 1.534396 | 592 | UP | micro1 | Protein degradation |
| GOBP_PROTEIN_CATABOLIC_PROCESS | 0.001984931 | 0.082328697 | 0.431708 | 0.351307 | 1.259944 | 884 | UP | micro1 | Protein degradation |
| GOBP_REGULATION_OF_CATABOLIC_PROCESS | 1.81627E-05 | 0.002254203 | 0.57561 | 0.383552 | 1.377269 | 955 | UP | micro1 | Protein degradation |
| GOBP_REGULATION_OF_CELLULAR_CATABOLIC_PROCESS | 5.05211E-05 | 0.005345128 | 0.557332 | 0.386485 | 1.378043 | 806 | UP | micro1 | Protein degradation |
| GOBP_RNA_CATABOLIC_PROCESS | 6.75043E-09 | 2.07457E-06 | 0.761461 | 0.503614 | 1.731458 | 403 | UP | micro1 | RNA catabolism |
| GOMF_CELL_ADHESION_MOLECULE_BINDING | 2.4693E-06 | 0.000398409 | 0.627257 | 0.440874 | 1.535624 | 507 | UP | micro1 | Membrane adhesion |
| DESCARTES_FETAL_MUSCLE_SCHWANN_CELLS | 0.000434613 | 0.027475213 | 0.498493 | 0.554423 | 1.667898 | 106 | UP | micro1 | Smooth muscle development |
| FAN_OVARY_CL14_MATURE_SMOOTH_MUSCLE_CELL | 4.55867E-05 | 0.004838942 | 0.557332 | 0.462504 | 1.552208 | 296 | UP | micro1 | Smooth muscle development |
| HP_PECTORAL_MUSCLE_HYPOPLASIA_APLASIA | 0.000856876 | 0.046706957 | 0.477271 | 0.974369 | 1.630692 | 6 | UP | micro1 | Smooth muscle development |
| RUBENSTEIN_SKELETAL_MUSCLE_B_CELLS | 3.98555E-15 | 4.76332E-12 | 0.996986 | 0.698703 | 2.217261 | 163 | UP | micro1 | Immune function |
| RUBENSTEIN_SKELETAL_MUSCLE_MYELOID_CELLS | 2.00903E-08 | 5.63735E-06 | 0.733762 | 0.517992 | 1.747979 | 327 | UP | micro1 | Immune function |
| RUBENSTEIN_SKELETAL_MUSCLE_NK_CELLS | 5.72643E-08 | 1.39588E-05 | 0.719513 | 0.561971 | 1.827077 | 207 | UP | micro1 | Immune function |
| RUBENSTEIN_SKELETAL_MUSCLE_PCV_ENDOTHELIAL_CELLS | 6.76884E-08 | 1.61795E-05 | 0.704976 | 0.557712 | 1.818479 | 215 | UP | micro1 | Smooth muscle development |
| RUBENSTEIN_SKELETAL_MUSCLE_SATELLITE_CELLS | 8.92838E-22 | 2.8811E-17 | 1.203975 | 0.67083 | 2.255043 | 299 | UP | micro1 | Smooth muscle development |
| RUBENSTEIN_SKELETAL_MUSCLE_SMOOTH_MUSCLE_CELLS | 4.66583E-08 | 1.17626E-05 | 0.719513 | 0.480921 | 1.659292 | 428 | UP | micro1 | Smooth muscle development |
| RUBENSTEIN_SKELETAL_MUSCLE_T_CELLS | 9.35569E-15 | 9.57716E-12 | 0.986546 | 0.68915 | 2.189357 | 169 | UP | micro1 | Immune function |
| TRAVAGLINI_LUNG_AIRWAY_SMOOTH_MUSCLE_CELL | 6.15758E-05 | 0.006228808 | 0.538434 | 0.532889 | 1.67785 | 153 | UP | micro1 | Smooth muscle development |
| GOBP_NEURON_DIFFERENTIATION | 0.000269109 | 0.019427029 | 0.498493 | 0.349051 | 1.269164 | 1268 | UP | micro1 | Cell differentiation |
| RIZ_ERYTHROID_DIFFERENTIATION_6HR | 0.001956374 | 0.081563612 | 0.431708 | 0.741074 | 1.778005 | 26 | UP | micro1 | Cell differentiation |
| GOCC_NEURON_TO_NEURON_SYNAPSE | 0.00101794 | 0.053232712 | 0.45506 | 0.417886 | 1.416174 | 336 | UP | micro1 | Synaptic function |
| GOCC_POSTSYNAPSE | 0.000570932 | 0.033680823 | 0.477271 | 0.385366 | 1.355998 | 594 | UP | micro1 | Synaptic function |
| GOCC_SYNAPSE | 1.12377E-06 | 0.00019392 | 0.643552 | 0.383676 | 1.391813 | 1228 | UP | micro1 | Synaptic function |
| GOBP_CATECHOLAMINE_UPTAKE_INVOLVED_IN_SYNAPTIC_TR | 0.000590361 | 0.034574179 | 0.477271 | 0.883767 | 1.767957 | 12 | UP | micro1 | Synaptic function |
| GOBP_REGULATION_OF_SYNAPTIC_TRANSMISSION_GLUTAMA | 0.002279419 | 0.089373706 | 0.431708 | 0.584614 | 1.644749 | 67 | UP | micro1 | Synaptic function |
| GOBP_SYNAPTIC_SIGNALING | 0.00137781 | 0.064700946 | 0.45506 | 0.367113 | 1.298533 | 677 | UP | micro1 | Synaptic function |

| | | | | | | | | | |
|---|---|---|---|---|---|---|---|---|---|
| GOBP_SYNAPTIC_TRANSMISSION_DOPAMINERGIC | 0.001298661 | 0.062453809 | 0.45506 | 0.758016 | 1.779683 | 24 | UP | micro1 | Synaptic function |
| MANNO_MIDBRAIN_NEUROTYPES_HGABA | 0.000153083 | 0.012389007 | 0.518848 | 0.362865 | 1.305894 | 1038 | UP | micro1 | Synaptic function |
| MANNO_MIDBRAIN_NEUROTYPES_HNBGABA | 1.2767E-05 | 0.001661202 | 0.593325 | 0.40216 | 1.421 | 654 | UP | micro1 | Synaptic function |
| GOBP_REGULATION_OF_STEROID_METABOLIC_PROCESS | 0.002350656 | 0.090761889 | 0.431708 | 0.511399 | 1.530547 | 102 | UP | micro1 | Hormone regulation |
| GOBP_REGULATION_OF_HORMONE_LEVELS | 0.001084861 | 0.055216697 | 0.45506 | 0.399522 | 1.380498 | 438 | UP | micro1 | Hormone regulation |
| GOBP_REGULATION_OF_SYSTEMIC_ARTERIAL_BLOOD_PRESSU | 0.002446759 | 0.093106674 | 0.431708 | 0.722747 | 1.743477 | 27 | UP | micro1 | Hormone regulation |
| GOBP_ATP_METABOLIC_PROCESS | 0.001083739 | 0.055216697 | 0.45506 | 0.4339 | 1.453654 | 291 | UP | micro1 | Energy production |
| BLANCO_MELO_RESPIRATORY_SYNCYTIAL_VIRUS_INFECTION_ | 3.63262E-08 | 9.53017E-06 | 0.719513 | 0.541728 | 1.798472 | 251 | UP | micro1 | Energy production |
| GOBP_MRNA_METABOLIC_PROCESS | 4.17961E-08 | 1.07041E-05 | 0.719513 | 0.423251 | 1.508418 | 808 | UP | micro1 | RNA metabolism |
| GOMF_MRNA_BINDING | 0.000582819 | 0.034304549 | 0.477271 | 0.444249 | 1.48344 | 272 | UP | micro1 | RNA processing & binding |
| GOMF_RNA_BINDING | 1.10505E-10 | 5.09412E-08 | 0.839089 | 0.399732 | 1.459365 | 1557 | UP | micro1 | RNA processing & binding |
| GOMF_RRNA_BINDING | 0.000313243 | 0.021691095 | 0.498493 | 0.628643 | 1.746681 | 61 | UP | micro1 | RNA processing & binding |
| GSE14000_TRANSLATED_RNA_VS_MRNA_DC_DN | 0.000826151 | 0.045571072 | 0.477271 | 0.479948 | 1.524746 | 169 | UP | micro1 | DNA/RNA synthesis & repair |
| GSE21360_NAIVE_VS_QUATERNARY_MEMORY_CD8_TCELL_DN | 7.51235E-07 | 0.000134676 | 0.659444 | 0.551083 | 1.775178 | 192 | UP | micro1 | Immune function |
| GSE21360_NAIVE_VS_QUATERNARY_MEMORY_CD8_TCELL_UP | 1.41434E-07 | 3.10473E-05 | 0.690132 | 0.563692 | 1.815524 | 193 | UP | micro1 | Immune function |
| GSE21360_PRIMARY_VS_QUATERNARY_MEMORY_CD8_TCELL_ | 2.73836E-05 | 0.00321324 | 0.57561 | 0.534143 | 1.696917 | 165 | UP | micro1 | Immune function |
| GSE21360_TERTIARY_VS_QUATERNARY_MEMORY_CD8_TCELL_ | 0.000835409 | 0.045846621 | 0.477271 | 0.478234 | 1.526409 | 174 | UP | micro1 | Immune function |
| HP_HYPERNATRIURIA | 0.001038705 | 0.053838093 | 0.45506 | 0.908945 | 1.706783 | 9 | UP | micro1 | Other |
| REACTOME_ACTIVATION_OF_THE_MRNA_UPON_BINDING_OF_ | 7.35759E-06 | 0.001045913 | 0.610527 | 0.689475 | 1.900933 | 59 | UP | micro1 | RNA processing & binding |
| REACTOME_CELLULAR_RESPONSES_TO_EXTERNAL_STIMULI | 5.40003E-13 | 3.8723E-10 | 0.921426 | 0.500837 | 1.763172 | 601 | UP | micro1 | Other |
| REACTOME_METABOLISM_OF_RNA | 4.33998E-07 | 8.28679E-05 | 0.674963 | 0.428261 | 1.512288 | 650 | UP | micro1 | RNA metabolism |
| REACTOME_RRNA_PROCESSING | 1.70886E-10 | 7.4518E-08 | 0.826657 | 0.614375 | 1.982227 | 195 | UP | micro1 | RNA processing & binding |
| GOBP_CYTOPLASMIC_TRANSLATION | 1.46494E-07 | 3.18275E-05 | 0.690132 | 0.655182 | 1.943869 | 96 | UP | micro1 | Protein translation |
| GOBP_TRANSLATIONAL_INITIATION | 7.81859E-12 | 4.76034E-09 | 0.875325 | 0.638004 | 2.044407 | 184 | UP | micro1 | Protein translation |
| REACTOME_EUKARYOTIC_TRANSLATION_ELONGATION | 4.92814E-19 | 3.18053E-15 | 1.123915 | 0.844665 | 2.487001 | 88 | UP | micro1 | Protein translation |
| REACTOME_EUKARYOTIC_TRANSLATION_INITIATION | 3.60674E-15 | 4.65543E-12 | 0.996986 | 0.753089 | 2.300204 | 115 | UP | micro1 | Protein translation |
| REACTOME_POST_TRANSLATIONAL_PROTEIN_MODIFICATION | 3.15862E-05 | 0.003576337 | 0.557332 | 0.359906 | 1.308735 | 1291 | UP | micro1 | Protein translation |
| REACTOME_TRANSLATION | 3.70441E-11 | 1.86778E-08 | 0.851339 | 0.573904 | 1.919022 | 286 | UP | micro1 | Protein translation |
| GOBP_RIBOSOME_ASSEMBLY | 0.000148894 | 0.012132951 | 0.518848 | 0.658333 | 1.808156 | 58 | UP | micro1 | Protein translation |
| GOCC_CYTOSOLIC_RIBOSOME | 1.46822E-16 | 3.38414E-13 | 1.047626 | 0.80009 | 2.379158 | 98 | UP | micro1 | Protein translation |
| GOCC_POLYSOMAL_RIBOSOME | 1.24834E-05 | 0.001637509 | 0.593325 | 0.811567 | 1.983976 | 30 | UP | micro1 | Protein translation |
| GOCC_RIBOSOME | 2.14422E-13 | 1.64743E-10 | 0.943632 | 0.640482 | 2.084008 | 211 | UP | micro1 | Protein translation |
| GOMF_STRUCTURAL_CONSTITUENT_OF_RIBOSOME | 9.38562E-15 | 9.57716E-12 | 0.986546 | 0.709471 | 2.231852 | 151 | UP | micro1 | Protein translation |
| KEGG_RIBOSOME | 4.05156E-18 | 2.179E-14 | 1.095929 | 0.846702 | 2.456284 | 83 | UP | micro1 | Protein translation |
| GOBP_RIBOSOMAL_SMALL_SUBUNIT_ASSEMBLY | 0.001128774 | 0.056210495 | 0.45506 | 0.804524 | 1.770321 | 18 | UP | micro1 | Protein translation |
| GOCC_CYTOSOLIC_LARGE_RIBOSOMAL_SUBUNIT | 2.8759E-10 | 1.23737E-07 | 0.814036 | 0.83344 | 2.230172 | 51 | UP | micro1 | Protein translation |
| GOCC_CYTOSOLIC_SMALL_RIBOSOMAL_SUBUNIT | 1.46961E-07 | 3.18275E-05 | 0.690132 | 0.794679 | 2.054613 | 43 | UP | micro1 | Protein translation |
| GOCC_LARGE_RIBOSOMAL_SUBUNIT | 5.02198E-08 | 1.25623E-05 | 0.719513 | 0.655257 | 1.981262 | 108 | UP | micro1 | Protein translation |
| GOCC_RIBOSOMAL_SUBUNIT | 1.41386E-13 | 1.23308E-10 | 0.943632 | 0.668654 | 2.133794 | 177 | UP | micro1 | Protein translation |
| GOCC_SMALL_RIBOSOMAL_SUBUNIT | 5.40937E-06 | 0.000797055 | 0.610527 | 0.678214 | 1.92365 | 71 | UP | micro1 | Protein translation |
| WP_CYTOPLASMIC_RIBOSOMAL_PROTEINS | 6.03115E-17 | 1.76926E-13 | 1.057464 | 0.829539 | 2.412618 | 86 | UP | micro1 | Protein translation |

Macrophage.0

| pathway | pval | padj | log2err | ES | NES | size | dir | cell | class |
|---|---|---|---|---|---|---|---|---|---|
| GOBP_MUSCLE_ORGAN_MORPHOGENESIS | 0.000758574 | 0.095246792 | 0.477271 | -0.634084 | -1.725984 | 57 | DOWN | mac0 | Smooth muscle development |
| GOBP_RIBONUCLEOPROTEIN_COMPLEX_BIOGENESIS | 0.00029071 | 0.049114704 | 0.498493 | 0.410723 | 1.383646 | 423 | UP | mac0 | Protein translation |
| GOBP_RIBOSOME_BIOGENESIS | 0.000730002 | 0.093850358 | 0.477271 | 0.427878 | 1.401158 | 292 | UP | mac0 | Protein translation |
| GOCC_CELL_SUBSTRATE_JUNCTION | 6.7636E-05 | 0.017023546 | 0.538434 | 0.425928 | 1.435123 | 416 | UP | mac0 | Cell junctions |
| GOBP_CELLULAR_MACROMOLECULE_LOCALIZATION | 1.65643E-06 | 0.000703305 | 0.643552 | 0.353119 | 1.283023 | 1849 | UP | mac0 | Protein localization |
| GOBP_ESTABLISHMENT_OF_PROTEIN_LOCALIZATION | 9.34284E-07 | 0.000432739 | 0.659444 | 0.355449 | 1.291419 | 1838 | UP | mac0 | Protein localization |
| GOBP_ESTABLISHMENT_OF_PROTEIN_LOCALIZATION_TO_ENDOPLASMIC_RETICUL | 1.14465E-07 | 9.45204E-05 | 0.704976 | 0.65781 | 1.942945 | 110 | UP | mac0 | Protein localization |
| GOBP_ESTABLISHMENT_OF_PROTEIN_LOCALIZATION_TO_MEMBRANE | 1.24655E-07 | 9.57736E-05 | 0.690132 | 0.504046 | 1.665534 | 327 | UP | mac0 | Protein localization |
| GOBP_ESTABLISHMENT_OF_PROTEIN_LOCALIZATION_TO_ORGANELLE | 3.51029E-07 | 0.000205952 | 0.674963 | 0.44228 | 1.51571 | 536 | UP | mac0 | Protein localization |
| GOBP_PROTEIN_LOCALIZATION_TO_ENDOPLASMIC_RETICULUM | 9.61512E-07 | 0.000437 | 0.643552 | 0.598611 | 1.814905 | 137 | UP | mac0 | Protein localization |
| GOBP_PROTEIN_LOCALIZATION_TO_MEMBRANE | 2.2778E-05 | 0.00727745 | 0.57561 | 0.405035 | 1.402189 | 622 | UP | mac0 | Protein localization |
| GOBP_PROTEIN_LOCALIZATION_TO_ORGANELLE | 1.14841E-06 | 0.000507645 | 0.643552 | 0.392867 | 1.38693 | 941 | UP | mac0 | Protein localization |
| GOBP_COTRANSLATIONAL_PROTEIN_TARGETING_TO_MEMBRANE | 7.09215E-08 | 6.18531E-05 | 0.704976 | 0.680856 | 1.98324 | 98 | UP | mac0 | Protein localization |
| GOBP_PROTEIN_TARGETING | 4.2244E-08 | 4.5439E-05 | 0.719513 | 0.484778 | 1.63341 | 416 | UP | mac0 | Protein localization |
| GOBP_PROTEIN_TARGETING_TO_MEMBRANE | 1.68318E-09 | 5.43145E-06 | 0.788187 | 0.606782 | 1.905563 | 190 | UP | mac0 | Protein localization |
| REACTOME_SRP_DEPENDENT_COTRANSLATIONAL_PROTEIN_TARGETING_TO_MEM | 1.74003E-07 | 0.000119466 | 0.690132 | 0.645921 | 1.896857 | 107 | UP | mac0 | Protein localization |
| GOCC_ENDOPLASMIC_RETICULUM | 0.000202255 | 0.037945152 | 0.518848 | 0.337702 | 1.226088 | 1767 | UP | mac0 | Endoplasmic reticulum |
| GOMF_WATER_TRANSMEMBRANE_TRANSPORTER_ACTIVITY | 0.000364575 | 0.056938283 | 0.498493 | -0.990927 | -1.739013 | 8 | DOWN | mac0 | Solute transporter |
| GOBP_CELLULAR_MACROMOLECULE_CATABOLIC_PROCESS | 1.74418E-05 | 0.005802372 | 0.57561 | 0.3646 | 1.296816 | 1125 | UP | mac0 | Protein degradation |
| GOBP_MACROMOLECULE_CATABOLIC_PROCESS | 6.68634E-07 | 0.000322032 | 0.659444 | 0.372096 | 1.334732 | 1340 | UP | mac0 | Protein degradation |
| GOBP_NUCLEAR_TRANSCRIBED_MRNA_CATABOLIC_PROCESS | 8.75841E-06 | 0.003216148 | 0.593325 | 0.516182 | 1.642493 | 204 | UP | mac0 | RNA catabolism |
| GOBP_NUCLEAR_TRANSCRIBED_MRNA_CATABOLIC_PROCESS_NONSENSE_MEDIAT | 2.48098E-06 | 0.001000732 | 0.627257 | 0.612396 | 1.815921 | 118 | UP | mac0 | RNA catabolism |
| GOBP_ORGANIC_CYCLIC_COMPOUND_CATABOLIC_PROCESS | 0.000144844 | 0.030548873 | 0.518848 | 0.39373 | 1.357353 | 592 | UP | mac0 | Protein degradation |
| GOBP_RNA_CATABOLIC_PROCESS | 2.93336E-05 | 0.008846412 | 0.57561 | 0.438863 | 1.476097 | 403 | UP | mac0 | RNA catabolism |
| GOBP_REGULATION_OF_SKELETAL_MUSCLE_CONTRACTION_BY_CALCIUM_ION_SI | 0.000571389 | 0.08031884 | 0.477271 | -0.996422 | -1.531347 | 5 | DOWN | mac0 | Muscle contraction |
| GOBP_VENTRICULAR_CARDIAC_MUSCLE_TISSUE_DEVELOPMENT | 0.000271848 | 0.047417576 | 0.498493 | -0.698605 | -1.821606 | 45 | DOWN | mac0 | Smooth muscle development |
| RUBENSTEIN_SKELETAL_MUSCLE_B_CELLS | 4.11955E-07 | 0.000221556 | 0.674963 | 0.577334 | 1.793967 | 163 | UP | mac0 | Immune function |
| RUBENSTEIN_SKELETAL_MUSCLE_MYELOID_CELLS | 4.96665E-05 | 0.013582095 | 0.557332 | 0.452878 | 1.496456 | 327 | UP | mac0 | Immune function |
| RUBENSTEIN_SKELETAL_MUSCLE_NK_CELLS | 6.55962E-05 | 0.016667105 | 0.538434 | 0.483263 | 1.539367 | 207 | UP | mac0 | Immune function |
| RUBENSTEIN_SKELETAL_MUSCLE_PCV_ENDOTHELIAL_CELLS | 5.03358E-05 | 0.013584974 | 0.557332 | 0.49595 | 1.578204 | 215 | UP | mac0 | Smooth muscle development |
| RUBENSTEIN_SKELETAL_MUSCLE_SATELLITE_CELLS | 3.9279E-10 | 1.81071E-06 | 0.814036 | 0.548852 | 1.803573 | 299 | UP | mac0 | Smooth muscle development |
| RUBENSTEIN_SKELETAL_MUSCLE_SMOOTH_MUSCLE_CELLS | 8.77068E-06 | 0.003216148 | 0.593325 | 0.435418 | 1.469039 | 428 | UP | mac0 | Smooth muscle development |
| RUBENSTEIN_SKELETAL_MUSCLE_T_CELLS | 4.15866E-08 | 4.5439E-05 | 0.719513 | 0.598466 | 1.864191 | 169 | UP | mac0 | Immune function |
| GOBP_REGULATION_OF_MYOBLAST_DIFFERENTIATION | 0.000438545 | 0.065515828 | 0.498493 | -0.671643 | -1.74867 | 44 | DOWN | mac0 | Smooth muscle development |
| GOBP_MRNA_METABOLIC_PROCESS | 3.44527E-05 | 0.009926373 | 0.557332 | 0.382359 | 1.338187 | 808 | UP | mac0 | RNA metabolism |
| GOBP_NCRNA_METABOLIC_PROCESS | 0.000334104 | 0.053906016 | 0.498493 | 0.403465 | 1.368593 | 465 | UP | mac0 | RNA metabolism |
| GOBP_POSITIVE_REGULATION_OF_TRANSCRIPTION_BY_RNA_POLYMERASE_II | 0.000192999 | 0.037070787 | 0.518848 | 0.358213 | 1.266688 | 1029 | UP | mac0 | DNA/RNA synthesis & repair |
| GOMF_RNA_BINDING | 5.82893E-07 | 0.000289375 | 0.659444 | 0.365408 | 1.319006 | 1557 | UP | mac0 | RNA processing & binding |
| REACTOME_CELLULAR_RESPONSES_TO_EXTERNAL_STIMULI | 1.59353E-08 | 2.23572E-05 | 0.733762 | 0.451893 | 1.559607 | 601 | UP | mac0 | Other |
| REACTOME_METABOLISM_OF_RNA | 1.39079E-05 | 0.004825753 | 0.593325 | 0.40402 | 1.400176 | 650 | UP | mac0 | RNA metabolism |
| REACTOME_RRNA_PROCESSING | 1.56689E-07 | 0.00011234 | 0.690132 | 0.563128 | 1.776756 | 195 | UP | mac0 | RNA processing & binding |
| GOBP_CYTOPLASMIC_TRANSLATION | 0.00028217 | 0.048176453 | 0.498493 | 0.560541 | 1.632917 | 96 | UP | mac0 | Protein translation |
| GOBP_TRANSLATIONAL_INITIATION | 1.35354E-07 | 0.000101575 | 0.690132 | 0.581824 | 1.815817 | 184 | UP | mac0 | Protein translation |
| REACTOME_EUKARYOTIC_TRANSLATION_ELONGATION | 3.42071E-09 | 7.65739E-06 | 0.774939 | 0.732105 | 2.09051 | 88 | UP | mac0 | Protein translation |
| REACTOME_EUKARYOTIC_TRANSLATION_INITIATION | 9.0975E-08 | 7.72545E-05 | 0.704976 | 0.65579 | 1.944117 | 115 | UP | mac0 | Protein translation |
| REACTOME_TRANSLATION | 0.000177269 | 0.035093758 | 0.518848 | 0.449367 | 1.468377 | 286 | UP | mac0 | Protein translation |
| GOCC_CYTOSOLIC_RIBOSOME | 9.33377E-10 | 3.34657E-06 | 0.788187 | 0.718173 | 2.091938 | 98 | UP | mac0 | Protein translation |
| GOCC_RIBOSOME | 2.93991E-07 | 0.000175681 | 0.674963 | 0.544929 | 1.733015 | 211 | UP | mac0 | Protein translation |
| GOMF_STRUCTURAL_CONSTITUENT_OF_RIBOSOME | 1.23193E-06 | 0.000537206 | 0.643552 | 0.576844 | 1.770508 | 151 | UP | mac0 | Protein translation |
| KEGG_RIBOSOME | 1.40894E-08 | 2.08985E-05 | 0.74774 | 0.721611 | 2.045531 | 83 | UP | mac0 | Protein translation |
| GOCC_CYTOSOLIC_LARGE_RIBOSOMAL_SUBUNIT | 1.64434E-05 | 0.005538406 | 0.57561 | 0.73264 | 1.92002 | 51 | UP | mac0 | Protein translation |
| GOCC_CYTOSOLIC_SMALL_RIBOSOMAL_SUBUNIT | 0.00010984 | 0.024111856 | 0.538434 | 0.716785 | 1.818695 | 43 | UP | mac0 | Protein translation |
| GOCC_RIBOSOMAL_SUBUNIT | 4.33597E-07 | 0.000229373 | 0.674963 | 0.566284 | 1.762305 | 177 | UP | mac0 | Protein translation |
| GOCC_SMALL_RIBOSOMAL_SUBUNIT | 0.000366392 | 0.056938283 | 0.498493 | 0.604766 | 1.670006 | 71 | UP | mac0 | Protein translation |
| WP_CYTOPLASMIC_RIBOSOMAL_PROTEINS | 3.55948E-09 | 7.65739E-06 | 0.774939 | 0.732668 | 2.090141 | 86 | UP | mac0 | Protein translation |

Smooth muscle cells (SMCs)

| pathway | pval | padj | log2err | ES | NES | size | dir | cell | class |
|---|---|---|---|---|---|---|---|---|---|
| GOCC_ACTIN_CYTOSKELETON | 3.60411E-08 | 0.000582 | 0.719513 | 0.446804 | 1.619023 | 471 | UP | smc | Cytoskeleton |
| GOMF_ACTIN_BINDING | 9.1639E-08 | 0.000739 | 0.704976 | 0.454405 | 1.635699 | 417 | UP | smc | Cytoskeleton |
| GOCC_MICROTUBULE_CYTOSKELETON | 7.16652E-05 | 0.057433 | 0.538434 | 0.334118 | 1.276349 | 1201 | UP | smc | Cytoskeleton |
| GOMF_TUBULIN_BINDING | 0.000156833 | 0.078475 | 0.518848 | 0.404676 | 1.447453 | 352 | UP | smc | Cytoskeleton |
| GOBP_SECRETION | 0.000147599 | 0.078317 | 0.518848 | 0.324109 | 1.248969 | 1330 | UP | smc | Cell secretion |
| GOBP_CELL_MORPHOGENESIS | 6.39603E-05 | 0.052921 | 0.538434 | 0.346064 | 1.312163 | 956 | UP | smc | Cell survival/development/Proliferation |
| GOBP_CELL_MORPHOGENESIS_INVOLVED_IN_DIFFERENTIAT | 3.95889E-05 | 0.039678 | 0.557332 | 0.368701 | 1.374081 | 690 | UP | smc | Cell survival/development/Proliferation |
| GOBP_REGULATION_OF_CELLULAR_COMPONENT_BIOGENES | 0.000124171 | 0.073138 | 0.518848 | 0.348829 | 1.319335 | 889 | UP | smc | Cell survival/development/Proliferation |
| GOBP_ESTABLISHMENT_OF_ORGANELLE_LOCALIZATION | 7.29727E-05 | 0.057433 | 0.538434 | 0.404927 | 1.45678 | 415 | UP | smc | Protein localization |
| GOBP_ORGANELLE_LOCALIZATION | 9.74453E-05 | 0.06391 | 0.538434 | 0.370554 | 1.376913 | 627 | UP | smc | Protein localization |
| GOBP_MUSCLE_CONTRACTION | 0.000202575 | 0.084895 | 0.518848 | 0.41138 | 1.454926 | 323 | UP | smc | Smooth muscle contraction |
| GOBP_MUSCLE_SYSTEM_PROCESS | 0.000150474 | 0.078317 | 0.518848 | 0.395619 | 1.422349 | 398 | UP | smc | Smooth muscle contraction |
| GOBP_REGULATION_OF_CARDIAC_MUSCLE_CONTRACTION_ | 0.000217539 | 0.087945 | 0.518848 | 0.782374 | 1.913309 | 24 | UP | smc | Smooth muscle contraction |
| RUBENSTEIN_SKELETAL_MUSCLE_FAP_CELLS | 9.2014E-05 | 0.063174 | 0.538434 | -0.491311 | -1.614526 | 173 | DOWN | smc | Smooth muscle development |
| RUBENSTEIN_SKELETAL_MUSCLE_FBN1_FAP_CELLS | 8.95004E-06 | 0.018051 | 0.593325 | -0.464012 | -1.597716 | 276 | DOWN | smc | Smooth muscle development |
| GOCC_SYNAPSE | 6.64801E-06 | 0.014302 | 0.610527 | 0.34399 | 1.314831 | 1228 | UP | smc | Synaptic function |
| REACTOME_METABOLISM_OF_AMINE_DERIVED_HORMONE | 0.000169627 | 0.078647 | 0.518848 | 0.877336 | 1.961243 | 15 | UP | smc | Hormone regulation |
| GOCC_EXTERNAL_ENCAPSULATING_STRUCTURE | 3.37868E-05 | 0.036342 | 0.557332 | -0.405735 | -1.454902 | 483 | DOWN | smc | DNA/RNA synthesis & repair |
| GOCC_COLLAGEN_CONTAINING_EXTRACELLULAR_MATRIX | 6.61089E-06 | 0.014302 | 0.610527 | -0.435214 | -1.531558 | 367 | DOWN | smc | Extracellular matrix |
| AACTTT_UNKNOWN | 1.07782E-05 | 0.020459 | 0.593325 | 0.32654 | 1.273352 | 1779 | UP | smc | other |
| BUSSLINGER_GASTRIC_IMMUNE_CELLS | 5.62645E-05 | 0.050433 | 0.557332 | 0.32792 | 1.26827 | 1415 | UP | smc | Immune function |
| CHEN_METABOLIC_SYNDROM_NETWORK | 0.000270004 | 0.095744 | 0.498493 | -0.328412 | -1.240412 | 1196 | DOWN | smc | other |
| CREBP1CJUN_01 | 0.00016113 | 0.078475 | 0.518848 | 0.439725 | 1.514153 | 246 | UP | smc | other |
| CREBP1_Q2 | 6.51461E-06 | 0.014302 | 0.610527 | 0.48028 | 1.652068 | 240 | UP | smc | other |
| CREB_01 | 0.00024709 | 0.094921 | 0.498493 | 0.440604 | 1.520296 | 251 | UP | smc | other |
| CREB_02 | 0.000193915 | 0.08456 | 0.518848 | 0.437447 | 1.504315 | 248 | UP | smc | other |
| CUI_DEVELOPING_HEART_C3_FIBROBLAST_LIKE_CELL | 0.000266885 | 0.095744 | 0.498493 | -0.531304 | -1.639665 | 108 | DOWN | smc | Smooth muscle development |
| CUI_TCF21_TARGETS_2_DN | 3.05836E-05 | 0.034031 | 0.557332 | 0.361255 | 1.359368 | 825 | UP | smc | other |
| DUTERTRE_ESTRADIOL_RESPONSE_24HR_DN | 9.51903E-05 | 0.06391 | 0.538434 | 0.386763 | 1.405953 | 493 | UP | smc | other |
| FALVELLA_SMOKERS_WITH_LUNG_CANCER | 0.000162938 | 0.078475 | 0.518848 | 0.602951 | 1.775777 | 70 | UP | smc | other |
| GCM2_TARGET_GENES | 7.3724E-07 | 0.002974 | 0.659444 | 0.347194 | 1.346254 | 1469 | UP | smc | other |
| GNF2_SPINK1 | 0.000264439 | 0.095744 | 0.498493 | 0.897107 | 1.914409 | 13 | UP | smc | other |
| GOBP_CELL_PROJECTION_ORGANIZATION | 0.000148579 | 0.078317 | 0.518848 | 0.322762 | 1.253089 | 1485 | UP | smc | Cell projection |
| GOBP_CYTOSKELETON_ORGANIZATION | 3.57729E-07 | 0.001924 | 0.674963 | 0.351014 | 1.349909 | 1277 | UP | smc | other |
| GOBP_POSITIVE_REGULATION_OF_MOLECULAR_FUNCTION | 0.000230182 | 0.089491 | 0.518848 | 0.316138 | 1.226949 | 1593 | UP | smc | other |
| GOBP_PROTEIN_COMPLEX_OLIGOMERIZATION | 0.00010336 | 0.065398 | 0.538434 | 0.461429 | 1.579465 | 217 | UP | smc | other |
| GOBP_PROTEIN_CONTAINING_COMPLEX_SUBUNIT_ORGANI | 1.74577E-06 | 0.005633 | 0.643552 | 0.331493 | 1.294035 | 1733 | UP | smc | other |
| GOBP_REGULATION_OF_ION_TRANSPORT | 8.45532E-05 | 0.061724 | 0.538434 | 0.333158 | 1.273311 | 1189 | UP | smc | other |
| GOBP_REGULATION_OF_TRANSPORT | 1.96322E-07 | 0.001267 | 0.690132 | 0.344674 | 1.339098 | 1570 | UP | smc | other |
| GOCC_NEURON_PROJECTION | 1.54829E-05 | 0.026296 | 0.57561 | 0.342202 | 1.311779 | 1252 | UP | smc | Cell projection |
| GOMF_CYTOSKELETAL_PROTEIN_BINDING | 6.83726E-10 | 2.21E-05 | 0.801216 | 0.402182 | 1.524415 | 928 | UP | smc | Cytoskeleton |
| GOMF_ENZYME_BINDING | 2.68885E-05 | 0.034031 | 0.57561 | 0.322012 | 1.255809 | 1807 | UP | smc | other |
| GOMF_IDENTICAL_PROTEIN_BINDING | 0.00016234 | 0.078475 | 0.518848 | 0.310198 | 1.210363 | 1746 | UP | smc | other |
| GSE14769_UNSTIM_VS_40MIN_LPS_BMDM_DN | 8.6486E-05 | 0.061724 | 0.538434 | 0.471257 | 1.577129 | 192 | UP | smc | other |
| GSE14769_UNSTIM_VS_80MIN_LPS_BMDM_DN | 0.000189068 | 0.083576 | 0.518848 | 0.467987 | 1.571521 | 188 | UP | smc | other |
| GSE23925_LIGHT_ZONE_VS_NAIVE_BCELL_UP | 2.40403E-05 | 0.034031 | 0.57561 | 0.497241 | 1.667613 | 189 | UP | smc | other |
| GSE36891_POLYIC_TLR3_VS_PAM_TLR2_STIM_PERITONEAL_ | 0.000132168 | 0.074823 | 0.518848 | 0.502319 | 1.633407 | 142 | UP | smc | other |
| GSE36891_UNSTIM_VS_POLYIC_TLR3_STIM_PERITONEAL_M | 5.18231E-05 | 0.047779 | 0.557332 | 0.524837 | 1.689251 | 132 | UP | smc | other |
| GSE37605_C57BL6_VS_NOD_FOXP3_FUSION_GFP_TREG_DN | 0.000219854 | 0.087945 | 0.518848 | 0.465148 | 1.554723 | 182 | UP | smc | other |
| GSE37605_FOXP3_FUSION_GFP_VS_IRES_GFP_TREG_C57BL( | 8.79886E-05 | 0.061724 | 0.538434 | 0.488511 | 1.629757 | 180 | UP | smc | other |
| GSE37605_TREG_VS_TCONV_NOD_FOXP3_FUSION_GFP_UP | 0.000218593 | 0.087945 | 0.518848 | 0.516521 | 1.67214 | 133 | UP | smc | other |
| GTGACGY_E4F1_Q6 | 1.95186E-06 | 0.005726 | 0.627257 | 0.393556 | 1.462241 | 630 | UP | smc | other |
| GTGCAAT_MIR25_MIR32_MIR92_MIR363_MIR367 | 3.55827E-05 | 0.037039 | 0.557332 | 0.436965 | 1.531181 | 292 | UP | smc | other |
| HMGB1_TARGET_GENES | 4.05772E-05 | 0.039678 | 0.557332 | 0.330281 | 1.277787 | 1388 | UP | smc | other |
| HP_ABNORMALITY_OF_COORDINATION | 0.000123871 | 0.073138 | 0.518848 | 0.339921 | 1.289323 | 955 | UP | smc | other |
| HP_ABNORMAL_ORAL_PHYSIOLOGY | 0.000264695 | 0.095744 | 0.498493 | 0.444932 | 1.520422 | 220 | UP | smc | other |
| HP_NEOPLASM_OF_THE_GASTROINTESTINAL_TRACT | 0.000226155 | 0.088997 | 0.518848 | -0.442388 | -1.5039 | 242 | DOWN | smc | other |
| HP_PANCREATIC_PSEUDOCYST | 0.000170607 | 0.078647 | 0.518848 | 0.998893 | 1.638395 | 5 | UP | smc | other |
| KIM_GLIS2_TARGETS_UP | 6.09386E-05 | 0.051748 | 0.557332 | -0.591194 | -1.765753 | 85 | DOWN | smc | other |
| LAKE_ADULT_KIDNEY_C13_THICK_ASCENDING_LIMB | 0.000200381 | 0.084895 | 0.518848 | 0.517092 | 1.636459 | 118 | UP | smc | other |
| LAKE_ADULT_KIDNEY_C14_DISTAL_CONVOLUTED_TUBULE | 5.88066E-08 | 0.000633 | 0.719513 | 0.556235 | 1.862047 | 191 | UP | smc | other |
| LAKE_ADULT_KIDNEY_C15_CONNECTING_TUBULE | 1.33906E-05 | 0.024006 | 0.593325 | 0.514033 | 1.704231 | 166 | UP | smc | other |
| LAKE_ADULT_KIDNEY_C17_COLLECTING_SYSTEM_PCS_STRE: | 5.02776E-05 | 0.047718 | 0.557332 | 0.467071 | 1.604088 | 230 | UP | smc | other |
| LAKE_ADULT_KIDNEY_C3_PROXIMAL_TUBULE_EPITHELIAL_( | 1.55404E-06 | 0.005572 | 0.643552 | 0.508206 | 1.719569 | 201 | UP | smc | other |
| LAKE_ADULT_KIDNEY_C4_PROXIMAL_TUBULE_EPITHELIAL_( | 5.09915E-06 | 0.013712 | 0.610527 | 0.523122 | 1.733479 | 165 | UP | smc | other |
| MIR19A_3P | 0.000158034 | 0.078475 | 0.518848 | 0.367595 | 1.365991 | 628 | UP | smc | other |
| MIR19B_3P | 9.90263E-05 | 0.06391 | 0.538434 | 0.366723 | 1.362543 | 630 | UP | smc | other |
| MIR25_3P | 0.000125202 | 0.073138 | 0.518848 | 0.402745 | 1.45165 | 405 | UP | smc | other |
| MIR32_5P | 1.85757E-05 | 0.029971 | 0.57561 | 0.404856 | 1.464477 | 460 | UP | smc | other |
| MIR363_3P | 0.000110198 | 0.068385 | 0.538434 | 0.398384 | 1.435929 | 410 | UP | smc | other |
| MIR367_3P | 0.000167818 | 0.078647 | 0.518848 | 0.397733 | 1.431159 | 411 | UP | smc | other |
| MIR4470 | 0.000126925 | 0.073138 | 0.518848 | 0.448511 | 1.542787 | 240 | UP | smc | other |
| MIR8063 | 2.72734E-05 | 0.034031 | 0.57561 | 0.415453 | 1.489761 | 391 | UP | smc | other |
| MIR92A_3P | 2.93394E-05 | 0.034031 | 0.57561 | 0.405584 | 1.463441 | 461 | UP | smc | other |
| MIR92B_3P | 2.57611E-05 | 0.034031 | 0.57561 | 0.404342 | 1.463493 | 458 | UP | smc | other |
| MIR9985 | 0.00026879 | 0.095744 | 0.498493 | 0.363017 | 1.343454 | 598 | UP | smc | other |
| MURARO_PANCREAS_MESENCHYMAL_STROMAL_CELL | 0.000264844 | 0.095744 | 0.498493 | -0.360082 | -1.318436 | 659 | DOWN | smc | other |
| NAGASHIMA_NRG1_SIGNALING_UP | 2.79082E-05 | 0.034031 | 0.57561 | 0.51591 | 1.699795 | 158 | UP | smc | other |
| NAKAYA_PLASMACYTOID_DENDRITIC_CELL_FLUMIST_AGE_1 | 5.90906E-05 | 0.051535 | 0.557332 | 0.338189 | 1.291476 | 1186 | UP | smc | other |
| PAX3_TARGET_GENES | 3.04123E-05 | 0.034031 | 0.57561 | 0.340077 | 1.300211 | 1222 | UP | smc | other |
| PEREZ_TP53_TARGETS | 8.09648E-05 | 0.060759 | 0.538434 | 0.341218 | 1.297252 | 1055 | UP | smc | other |
| PID_ERBB1_DOWNSTREAM_PATHWAY | 2.2498E-05 | 0.034031 | 0.57561 | 0.5725 | 1.778682 | 104 | UP | smc | other |
| REACTOME_NERVOUS_SYSTEM_DEVELOPMENT | 0.000143543 | 0.078317 | 0.518848 | 0.373722 | 1.373829 | 558 | UP | smc | Cell survival/development/Proliferation |

| | | | | | | | | | |
|---|---|---|---|---|---|---|---|---|---|
| REACTOME_SIGNALING_BY_RHO_GTPASES_MIRO_GTPASES_ | 0.000220754 | 0.087945 | 0.518848 | 0.363431 | 1.35306 | 639 | UP | smc | other |
| TGACATY_UNKNOWN | 0.000196795 | 0.084672 | 0.518848 | 0.365781 | 1.356906 | 608 | UP | smc | other |
| TGACCTTG_SF1_Q6 | 0.000266541 | 0.095744 | 0.498493 | 0.435908 | 1.49729 | 238 | UP | smc | other |
| TGACCTY_ERR1_Q2 | 7.70464E-05 | 0.059196 | 0.538434 | 0.34561 | 1.312482 | 977 | UP | smc | other |
| TGACGTCA_ATF3_Q6 | 0.00017522 | 0.079636 | 0.518848 | 0.455374 | 1.556908 | 219 | UP | smc | other |
| TGAYRTCA_ATF3_Q6 | 6.79886E-07 | 0.002974 | 0.659444 | 0.430054 | 1.560616 | 488 | UP | smc | other |
| TTANTCA_UNKNOWN | 0.000139425 | 0.077571 | 0.518848 | 0.347194 | 1.30876 | 852 | UP | smc | other |
| YAMASHITA_LIVER_CANCER_STEM_CELL_DN | 0.000183557 | 0.082267 | 0.518848 | 0.625162 | 1.794577 | 59 | UP | smc | other |

#### Pericytes

| pathway | pval | padj | log2err | ES | NES | size | dir | cell | class |
|---|---|---|---|---|---|---|---|---|---|
| GOBP_ESTABLISHMENT_OF_PROTEIN_LOCALIZATION_TO_ENDOPLASMIC | 1.16088E-05 | 0.024974 | 0.593325 | 0.638169 | 1.726248 | 110 | UP | pericyte | Protein localization |
| GOBP_PROTEIN_LOCALIZATION_TO_ENDOPLASMIC_RETICULUM | 6.07017E-05 | 0.043626 | 0.557332 | 0.590854 | 1.643418 | 137 | UP | pericyte | Protein localization |
| GOBP_COTRANSLATIONAL_PROTEIN_TARGETING_TO_MEMBRANE | 3.37365E-05 | 0.036438 | 0.557332 | 0.646027 | 1.723065 | 98 | UP | pericyte | Protein localization |
| REACTOME_SRP_DEPENDENT_COTRANSLATIONAL_PROTEIN_TARGETING | 2.27684E-05 | 0.031944 | 0.57561 | 0.641771 | 1.730014 | 107 | UP | pericyte | Protein localization |
| GOBP_CELLULAR_MACROMOLECULE_CATABOLIC_PROCESS | 0.000153528 | 0.07741 | 0.518848 | 0.39753 | 1.275408 | 1125 | UP | pericyte | Protein degradation |
| GOBP_NUCLEAR_TRANSCRIBED_MRNA_CATABOLIC_PROCESS_NONSENS | 1.47933E-05 | 0.029835 | 0.593325 | 0.631264 | 1.729451 | 118 | UP | pericyte | RNA catabolism |
| GOBP_RNA_CATABOLIC_PROCESS | 5.74869E-05 | 0.043626 | 0.557332 | 0.471588 | 1.437274 | 403 | UP | pericyte | RNA catabolism |
| RUBENSTEIN_SKELETAL_MUSCLE_SATELLITE_CELLS | 4.34189E-07 | 0.003043 | 0.674963 | 0.55099 | 1.647685 | 299 | UP | pericyte | Smooth muscle development |
| RUBENSTEIN_SKELETAL_MUSCLE_T_CELLS | 1.36351E-06 | 0.006286 | 0.643552 | 0.609369 | 1.731393 | 169 | UP | pericyte | Immune function |
| GOBP_TRANSLATIONAL_INITIATION | 2.13884E-05 | 0.031944 | 0.57561 | 0.55311 | 1.581139 | 184 | UP | pericyte | Protein translation |
| REACTOME_EUKARYOTIC_TRANSLATION_ELONGATION | 2.80475E-05 | 0.032324 | 0.57561 | 0.678167 | 1.77956 | 88 | UP | pericyte | Protein translation |
| REACTOME_EUKARYOTIC_TRANSLATION_INITIATION | 5.30612E-05 | 0.043626 | 0.557332 | 0.616737 | 1.678404 | 115 | UP | pericyte | Protein translation |
| REACTOME_POST_TRANSLATIONAL_PROTEIN_MODIFICATION | 0.000104162 | 0.062244 | 0.538434 | 0.390633 | 1.256707 | 1291 | UP | pericyte | Protein translation |
| REACTOME_TRANSLATION | 6.21897E-05 | 0.043626 | 0.538434 | 0.504123 | 1.49947 | 286 | UP | pericyte | Protein translation |
| GOCC_CYTOSOLIC_RIBOSOME | 0.000115886 | 0.064714 | 0.538434 | 0.625493 | 1.668298 | 98 | UP | pericyte | Protein translation |
| KEGG_RIBOSOME | 3.76558E-05 | 0.037972 | 0.557332 | 0.671361 | 1.743577 | 83 | UP | pericyte | Protein translation |
| BUSSLINGER_DUODENAL_DIFFERENTIATING_STEM_CELLS | 2.83952E-07 | 0.003043 | 0.674963 | 0.555949 | 1.655554 | 282 | UP | pericyte | Cell differentiation |
| BUSSLINGER_DUODENAL_STEM_CELLS | 0.000208042 | 0.098725 | 0.518848 | 0.484561 | 1.445489 | 294 | UP | pericyte | Cell differentiation |
| BUSSLINGER_DUODENAL_TRANSIT_AMPLIFYING_CELLS | 1.88965E-05 | 0.030489 | 0.57561 | 0.569277 | 1.625563 | 182 | UP | pericyte | other |
| DESCARTES_FETAL_HEART_VASCULAR_ENDOTHELIAL_CELLS | 7.38921E-05 | 0.050732 | 0.538434 | -0.98326 | -1.87319 | 13 | DOWN | pericyte | Cell junction |
| GENTLES_LEUKEMIC_STEM_CELL_DN | 7.57457E-05 | 0.050922 | 0.538434 | 0.971707 | 1.836428 | 15 | UP | pericyte | other |
| GNF2_CD7 | 2.27495E-05 | 0.031944 | 0.57561 | -0.88226 | -1.96933 | 26 | DOWN | pericyte | other |
| GNF2_MKI67 | 7.93561E-05 | 0.051756 | 0.538434 | 0.854883 | 1.862357 | 27 | UP | pericyte | other |
| GOBP_AMIDE_BIOSYNTHETIC_PROCESS | 4.8715E-05 | 0.042486 | 0.557332 | 0.426835 | 1.345472 | 761 | UP | pericyte | other |
| GOBP_BIOLOGICAL_PROCESS_INVOLVED_IN_SYMBIOTIC_INTERACTION | 2.07025E-07 | 0.003043 | 0.690132 | 0.4474 | 1.419169 | 926 | UP | pericyte | other |
| GOBP_CELLULAR_AMIDE_METABOLIC_PROCESS | 2.8021E-05 | 0.032324 | 0.57561 | 0.413093 | 1.317002 | 1024 | UP | pericyte | other |
| GOBP_INTRACELLULAR_TRANSPORT | 5.88904E-05 | 0.043626 | 0.557332 | 0.38325 | 1.24226 | 1617 | UP | pericyte | Solute transporter |
| GOBP_PEPTIDE_BIOSYNTHETIC_PROCESS | 0.000140579 | 0.073692 | 0.518848 | 0.431539 | 1.347682 | 630 | UP | pericyte | Protein translation |
| GOBP_PEPTIDE_METABOLIC_PROCESS | 4.35715E-05 | 0.039778 | 0.557332 | 0.425195 | 1.341808 | 765 | UP | pericyte | other |
| GOBP_VIRAL_GENE_EXPRESSION | 2.02227E-06 | 0.008157 | 0.627257 | 0.58268 | 1.673528 | 194 | UP | pericyte | Viral infection |
| GOCC_CYTOSOLIC_LARGE_RIBOSOMAL_SUBUNIT | 8.57577E-06 | 0.019767 | 0.593325 | 0.77889 | 1.892426 | 51 | UP | pericyte | Protein translation |
| GOCC_ENVELOPE | 6.11983E-05 | 0.043626 | 0.538434 | 0.403176 | 1.291672 | 1140 | UP | pericyte | other |
| GSE2405_0H_VS_24H_A_PHAGOCYTOPHILUM_STIM_NEUTROPHIL_UP | 9.41001E-05 | 0.058395 | 0.538434 | 0.534309 | 1.536713 | 198 | UP | pericyte | other |
| GSE2405_0H_VS_9H_A_PHAGOCYTOPHILUM_STIM_NEUTROPHIL_DN | 0.00013983 | 0.073692 | 0.518848 | 0.528482 | 1.519954 | 198 | UP | pericyte | other |
| GSE42088_UNINF_VS_LEISHMANIA_INF_DC_2H_DN | 5.89783E-05 | 0.043626 | 0.557332 | 0.55425 | 1.582212 | 183 | UP | pericyte | other |
| HSIAO_HOUSEKEEPING_GENES | 1.69583E-05 | 0.030489 | 0.57561 | 0.493392 | 1.498929 | 385 | UP | pericyte | other |
| KIM_ALL_DISORDERS_CALB1_CORR_UP | 0.000144339 | 0.073931 | 0.518848 | 0.441246 | 1.363105 | 538 | UP | pericyte | other |
| KIM_ALL_DISORDERS_OLIGODENDROCYTE_NUMBER_CORR_UP | 4.11122E-06 | 0.013111 | 0.610527 | 0.445173 | 1.402978 | 742 | UP | pericyte | other |
| LAKE_ADULT_KIDNEY_C10_THIN_ASCENDING_LIMB | 4.34402E-05 | 0.039778 | 0.557332 | 0.487505 | 1.469088 | 347 | UP | pericyte | other |
| LAKE_ADULT_KIDNEY_C12_THICK_ASCENDING_LIMB | 1.76689E-05 | 0.030489 | 0.57561 | 0.490409 | 1.48264 | 367 | UP | pericyte | other |
| LAKE_ADULT_KIDNEY_C18_COLLECTING_DUCT_PRINCIPAL_CELLS_MEDU | 3.47376E-06 | 0.012455 | 0.627257 | 0.518928 | 1.55601 | 313 | UP | pericyte | other |
| LAKE_ADULT_KIDNEY_C8_DECENDING_THIN_LIMB | 3.3876E-05 | 0.036438 | 0.557332 | 0.517831 | 1.531144 | 270 | UP | pericyte | other |
| LAKE_ADULT_KIDNEY_C9_THIN_ASCENDING_LIMB | 1.05885E-06 | 0.005695 | 0.643552 | 0.551878 | 1.631815 | 270 | UP | pericyte | other |
| LI_AMPLIFIED_IN_LUNG_CANCER | 9.93766E-05 | 0.060505 | 0.538434 | 0.559629 | 1.586528 | 168 | UP | pericyte | other |
| MENON_FETAL_KIDNEY_1_EMBRYONIC_RED_BLOOD_CELLS | 0.000213583 | 0.099886 | 0.518848 | 0.619741 | 1.649642 | 96 | UP | pericyte | other |
| MODULE_114 | 2.70448E-05 | 0.032324 | 0.57561 | 0.498107 | 1.499373 | 336 | UP | pericyte | other |
| MODULE_151 | 2.45167E-05 | 0.032324 | 0.57561 | 0.502445 | 1.506698 | 315 | UP | pericyte | other |
| MODULE_83 | 5.48427E-05 | 0.043626 | 0.557332 | 0.498429 | 1.496259 | 318 | UP | pericyte | other |
| MORF_ACTG1 | 0.00016831 | 0.083557 | 0.518848 | 0.576082 | 1.602331 | 137 | UP | pericyte | other |
| MORF_CSNK2B | 0.000116316 | 0.064714 | 0.538434 | 0.502097 | 1.495189 | 282 | UP | pericyte | other |
| MORF_NME2 | 0.000141589 | 0.073692 | 0.518848 | 0.564628 | 1.588732 | 153 | UP | pericyte | other |
| MORF_NPM1 | 4.43774E-05 | 0.039778 | 0.557332 | 0.584105 | 1.656018 | 160 | UP | pericyte | other |
| MORF_SOD1 | 0.000173393 | 0.084776 | 0.518848 | 0.49635 | 1.472466 | 274 | UP | pericyte | other |
| MORF_TPT1 | 8.0194E-05 | 0.051756 | 0.538434 | 0.630481 | 1.685091 | 101 | UP | pericyte | other |
| REACTOME_INFLUENZA_INFECTION | 8.32164E-05 | 0.052653 | 0.538434 | 0.580089 | 1.625964 | 147 | UP | pericyte | Viral infection |
| REACTOME_METABOLISM_OF_AMINO_ACIDS_AND_DERIVATIVES | 6.11762E-05 | 0.043626 | 0.538434 | 0.481287 | 1.449645 | 351 | UP | pericyte | other |
| REACTOME_NONSENSE_MEDIATED_DECAY_NMD | 6.33661E-06 | 0.015729 | 0.610527 | 0.646647 | 1.749182 | 110 | UP | pericyte | other |
| REACTOME_REGULATION_OF_EXPRESSION_OF_SLITS_AND_ROBOS | 4.80092E-06 | 0.013111 | 0.610527 | 0.591871 | 1.678572 | 165 | UP | pericyte | Cell survival/development/Proliferation |
| REACTOME_SELENOAMINO_ACID_METABOLISM | 0.000123334 | 0.067456 | 0.518848 | 0.61226 | 1.654751 | 109 | UP | pericyte | other |
| REACTOME_SIGNALING_BY_ROBO_RECEPTORS | 4.87558E-06 | 0.013111 | 0.610527 | 0.567971 | 1.638005 | 209 | UP | pericyte | Cell survival/development/Proliferation |
| REACTOME_TACHYKININ_RECEPTORS_BIND_TACHYKININS | 4.29059E-05 | 0.039778 | 0.557332 | 0.999276 | 1.444181 | 5 | UP | pericyte | other |
| SANA_RESPONSE_TO_IFNG_DN | 0.000180485 | 0.086927 | 0.518848 | 0.647988 | 1.684105 | 82 | UP | pericyte | other |
| TRAVAGLINI_LUNG_BRONCHIAL_VESSEL_1_CELL | 4.7153E-07 | 0.003043 | 0.674963 | 0.580294 | 1.682504 | 215 | UP | pericyte | other |
| TRAVAGLINI_LUNG_CD4_NAIVE_T_CELL | 0.000106141 | 0.062274 | 0.538434 | 0.60532 | 1.649451 | 116 | UP | pericyte | Immune function |
| TRAVAGLINI_LUNG_MESOTHELIAL_CELL | 3.3995E-08 | 0.001097 | 0.719513 | 0.491276 | 1.529007 | 596 | UP | pericyte | Immune function |
| TRAVAGLINI_LUNG_PROLIFERATING_BASAL_CELL | 3.71083E-05 | 0.037972 | 0.557332 | 0.420777 | 1.333011 | 834 | UP | pericyte | other |
| TRAVAGLINI_LUNG_PROXIMAL_BASAL_CELL | 1.85626E-05 | 0.030489 | 0.57561 | 0.450589 | 1.403746 | 589 | UP | pericyte | other |
| WP_CYTOPLASMIC_RIBOSOMAL_PROTEINS | 0.000112542 | 0.064714 | 0.538434 | 0.641572 | 1.678812 | 86 | UP | pericyte | Protein translation |
| ZHONG_PFC_C2_UNKNOWN_NPC | 2.75367E-05 | 0.032324 | 0.57561 | 0.688861 | 1.768894 | 73 | UP | pericyte | other |

Vascular astrocytes

| pathway | pval | padj | log2err | ES | NES | size | dir | cell | class |
|---|---|---|---|---|---|---|---|---|---|
| GOBP_NEUROGENESIS | 8.34739E-08 | 0.000336702 | 0.704976 | -0.386086 | -1.338054 | 1508 | DOWN | hybrid | Cell survival/development/Proliferation |
| GOBP_CELL_JUNCTION_ORGANIZATION | 3.91492E-05 | 0.03509184 | 0.557332 | -0.411275 | -1.376823 | 661 | DOWN | hybrid | Cell junctions |
| GOCC_ANCHORED_COMPONENT_OF_MEMBRANE | 3.25399E-05 | 0.0334384 | 0.557332 | -0.580982 | -1.694988 | 134 | DOWN | hybrid | Protein localization |
| GOBP_BIOLOGICAL_ADHESION | 1.63797E-08 | 0.000105711 | 0.733762 | -0.405487 | -1.394753 | 1313 | DOWN | hybrid | Membrane adhesion |
| GOBP_CELL_CELL_ADHESION | 1.51023E-06 | 0.0041251 | 0.643552 | -0.419283 | -1.413584 | 771 | DOWN | hybrid | Membrane adhesion |
| GOBP_CELL_CELL_ADHESION_VIA_PLASMA_MEMBRA | 7.57226E-07 | 0.002714991 | 0.659444 | -0.551466 | -1.715645 | 239 | DOWN | hybrid | Membrane adhesion |
| GOBP_HOMOPHILIC_CELL_ADHESION_VIA_PLASMA_I | 2.59252E-06 | 0.005761212 | 0.627257 | -0.603888 | -1.791732 | 152 | DOWN | hybrid | Membrane adhesion |
| DESCARTES_FETAL_MUSCLE_SCHWANN_CELLS | 1.66185E-06 | 0.0041251 | 0.643552 | -0.667207 | -1.891537 | 106 | DOWN | hybrid | Smooth muscle development |
| GOBERT_OLIGODENDROCYTE_DIFFERENTIATION_DN | 1.26797E-05 | 0.016366456 | 0.593325 | -0.388896 | -1.32938 | 1054 | DOWN | hybrid | Cell differentiation |
| GOBP_NEURON_DIFFERENTIATION | 6.64869E-06 | 0.011291923 | 0.610527 | -0.382979 | -1.317441 | 1268 | DOWN | hybrid | Cell differentiation |
| GOBP_SPINAL_CORD_ASSOCIATION_NEURON_DIFFER | 5.89917E-05 | 0.041560138 | 0.557332 | -0.989136 | -1.818496 | 12 | DOWN | hybrid | Cell differentiation |
| GOBP_SYNAPSE_ASSEMBLY | 0.000168096 | 0.093522348 | 0.518848 | -0.518898 | -1.558891 | 167 | DOWN | hybrid | Synaptic function |
| GOBP_SYNAPSE_ORGANIZATION | 4.83934E-05 | 0.039622059 | 0.557332 | -0.444926 | -1.433643 | 400 | DOWN | hybrid | Synaptic function |
| GOCC_SYNAPSE | 1.93626E-05 | 0.022314683 | 0.57561 | -0.378457 | -1.298436 | 1228 | DOWN | hybrid | Synaptic function |
| GOCC_POSTSYNAPTIC_MEMBRANE | 8.88796E-05 | 0.055126433 | 0.538434 | -0.487437 | -1.525612 | 257 | DOWN | hybrid | Synaptic function |
| GOCC_SYNAPTIC_MEMBRANE | 3.73793E-05 | 0.034462654 | 0.557332 | -0.464837 | -1.487098 | 358 | DOWN | hybrid | Synaptic function |
| GOMF_NEUROTRANSMITTER_RECEPTOR_ACTIVITY_IN | 1.62787E-05 | 0.019455401 | 0.57561 | -0.782756 | -1.933419 | 46 | DOWN | hybrid | Synaptic function |
| GOMF_POSTSYNAPTIC_NEUROTRANSMITTER_RECEPT | 3.3028E-06 | 0.006661124 | 0.627257 | -0.749748 | -1.9456 | 59 | DOWN | hybrid | Synaptic function |
| MANNO_MIDBRAIN_NEUROTYPES_HGABA | 9.31428E-06 | 0.013661925 | 0.593325 | -0.391619 | -1.339493 | 1038 | DOWN | hybrid | Synaptic function |
| MANNO_MIDBRAIN_NEUROTYPES_HNBGABA | 3.72729E-05 | 0.034462654 | 0.557332 | -0.414005 | -1.38188 | 654 | DOWN | hybrid | Synaptic function |
| GOBP_MRNA_METABOLIC_PROCESS | 6.99021E-05 | 0.046993131 | 0.538434 | 0.40215 | 1.315792 | 808 | UP | hybrid | RNA metabolism |
| BENPORATH_ES_WITH_H3K27ME3 | 2.67806E-06 | 0.005761212 | 0.627257 | -0.406719 | -1.381623 | 925 | DOWN | hybrid | other |
| BENPORATH_PRC2_TARGETS | 7.93508E-06 | 0.012802851 | 0.593325 | -0.451099 | -1.480553 | 508 | DOWN | hybrid | other |
| BENPORATH_SUZ12_TARGETS | 1.58482E-06 | 0.0041251 | 0.643552 | -0.410469 | -1.392415 | 855 | DOWN | hybrid | other |
| DESCARTES_FETAL_ADRENAL_SCHWANN_CELLS | 8.21577E-11 | 9.64622E-07 | 0.839089 | -0.761664 | -2.15939 | 107 | DOWN | hybrid | other |
| DESCARTES_FETAL_HEART_SCHWANN_CELLS | 7.70757E-05 | 0.049743112 | 0.538434 | -0.729621 | -1.839712 | 52 | DOWN | hybrid | other |
| DESCARTES_FETAL_INTESTINE_ENS_GLIA | 5.45058E-06 | 0.00977137 | 0.610527 | -0.776831 | -1.930719 | 48 | DOWN | hybrid | other |
| DESCARTES_FETAL_LUNG_VISCERAL_NEURONS | 0.000138243 | 0.079660164 | 0.518848 | -0.521627 | -1.574389 | 180 | DOWN | hybrid | other |
| DESCARTES_FETAL_PANCREAS_ENS_GLIA | 4.60761E-08 | 0.000247805 | 0.719513 | -0.813215 | -2.086833 | 56 | DOWN | hybrid | other |
| DESCARTES_FETAL_STOMACH_ENS_GLIA | 5.53369E-05 | 0.040583302 | 0.557332 | -0.771969 | -1.888526 | 44 | DOWN | hybrid | other |
| DESCARTES_FETAL_STOMACH_MUC13_DMBT1_POSIT | 1.21344E-05 | 0.01631525 | 0.593325 | 0.716553 | 1.830149 | 64 | UP | hybrid | other |
| DURANTE_ADULT_OLFACTORY_NEUROEPITHELIUM_C | 9.91784E-06 | 0.013914728 | 0.593325 | -0.634208 | -1.797991 | 108 | DOWN | hybrid | other |
| FAN_EMBRYONIC_CTX_BIG_GROUPS_GLIAL | 3.31597E-05 | 0.0334384 | 0.557332 | -0.586461 | -1.707968 | 132 | DOWN | hybrid | other |
| GOBP_CELL_PROJECTION_ORGANIZATION | 5.35008E-05 | 0.040149266 | 0.557332 | -0.356146 | -1.232968 | 1485 | DOWN | hybrid | Cell projection |
| GOBP_DORSAL_SPINAL_CORD_DEVELOPMENT | 4.62859E-05 | 0.039622059 | 0.557332 | -0.931433 | -1.906985 | 18 | DOWN | hybrid | Neuronal development |
| GOBP_G_PROTEIN_COUPLED_RECEPTOR_SIGNALING_ | 0.000156842 | 0.088791568 | 0.518848 | -0.383737 | -1.299148 | 840 | DOWN | hybrid | other |
| GOBP_NEURON_DEVELOPMENT | 3.61524E-05 | 0.034462654 | 0.557332 | -0.382556 | -1.307708 | 1054 | DOWN | hybrid | Neuronal development |
| GOMF_CALCIUM_ION_BINDING | 8.22726E-05 | 0.052055989 | 0.538434 | -0.408642 | -1.359315 | 620 | DOWN | hybrid | other |
| GOMF_EXTRACELLULAR_LIGAND_GATED_ION_CHANN | 7.21093E-05 | 0.047487625 | 0.538434 | -0.693367 | -1.816356 | 65 | DOWN | hybrid | other |
| GOMF_NEUROTRANSMITTER_RECEPTOR_ACTIVITY | 0.000119659 | 0.070204945 | 0.538434 | -0.613919 | -1.70362 | 92 | DOWN | hybrid | Synaptic function |
| GOMF_TRANSMITTER_GATED_CHANNEL_ACTIVITY | 2.81579E-05 | 0.03133198 | 0.57561 | -0.734312 | -1.870003 | 54 | DOWN | hybrid | Synaptic function |
| GSE46606_IRF4_KO_VS_WT_UNSTIM_BCELL_DN | 5.92447E-05 | 0.041560138 | 0.557332 | -0.533468 | -1.619443 | 189 | DOWN | hybrid | other |
| LOPEZ_MBD_TARGETS | 1.57059E-05 | 0.019455401 | 0.57561 | 0.404537 | 1.330566 | 902 | UP | hybrid | other |
| MANNO_MIDBRAIN_NEUROTYPES_HDA | 0.000173833 | 0.095074995 | 0.518848 | -0.424196 | -1.38268 | 472 | DOWN | hybrid | other |
| MANNO_MIDBRAIN_NEUROTYPES_HDA2 | 4.65603E-06 | 0.008837971 | 0.610527 | -0.456083 | -1.494618 | 488 | DOWN | hybrid | other |
| MANNO_MIDBRAIN_NEUROTYPES_HOPC | 2.14941E-11 | 6.93593E-07 | 0.863415 | -0.579279 | -1.838511 | 347 | DOWN | hybrid | other |
| MANNO_MIDBRAIN_NEUROTYPES_HRGL2A | 5.41452E-09 | 4.36803E-05 | 0.761461 | -0.486657 | -1.60461 | 551 | DOWN | hybrid | other |
| MANNO_MIDBRAIN_NEUROTYPES_HRGL2B | 1.20409E-06 | 0.003885476 | 0.643552 | -0.484529 | -1.564904 | 417 | DOWN | hybrid | other |
| MANNO_MIDBRAIN_NEUROTYPES_HRGL2C | 5.99444E-08 | 0.000276335 | 0.704976 | -0.541352 | -1.709699 | 305 | DOWN | hybrid | other |
| MIKKELSEN_MEF_HCP_WITH_H3K27ME3 | 3.11188E-05 | 0.0334384 | 0.557332 | -0.437608 | -1.434007 | 493 | DOWN | hybrid | other |
| MIR4261 | 5.04238E-05 | 0.039622059 | 0.557332 | -0.449889 | -1.451394 | 413 | DOWN | hybrid | other |
| MIR6867_5P | 9.0542E-05 | 0.055126433 | 0.538434 | -0.390664 | -1.321782 | 826 | DOWN | hybrid | other |
| MODULE_64 | 0.000109953 | 0.065704785 | 0.538434 | -0.436262 | -1.418507 | 454 | DOWN | hybrid | other |
| SABATES_COLORECTAL_ADENOMA_DN | 4.74492E-05 | 0.039622059 | 0.557332 | -0.502164 | -1.567377 | 251 | DOWN | hybrid | other |
| SETD7_TARGET_GENES | 0.00017797 | 0.095715272 | 0.518848 | 0.408788 | 1.333465 | 751 | UP | hybrid | other |
| SOBOLEV_PBMC_PANDEMRIX_AGE_18_64YO_MEDIU | 8.42793E-06 | 0.012950515 | 0.593325 | 0.828855 | 1.905849 | 35 | UP | hybrid | other |
| ZHONG_PFC_C9_ORG_OTHER | 6.80039E-05 | 0.046689713 | 0.538434 | -0.652978 | -1.780707 | 82 | DOWN | hybrid | other |
| ZHONG_PFC_MAJOR_TYPES_ASTROCYTES | 8.96794E-11 | 9.64622E-07 | 0.839089 | -0.577304 | -1.828897 | 327 | DOWN | hybrid | other |
| ZHONG_PFC_MAJOR_TYPES_OPC | 5.13891E-05 | 0.039622059 | 0.557332 | -0.644952 | -1.771767 | 89 | DOWN | hybrid | other |
| ZNF711_TARGET_GENES | 5.15704E-05 | 0.039622059 | 0.557332 | 0.376913 | 1.267744 | 1448 | UP | hybrid | other |

Table S2. DEGs in microglia and vascular cells, SwDI/TKO vs. SwDI/TWT

Microglia

| p_val | avg_log2FC | pct.1 | pct.2 | p_val_adj | genes |
|---|---|---|---|---|---|
| 6.26275E-37 | -0.26116 | 1 | 1 | 1.47074E-32 | Malat1 |
| 4.82696E-25 | 1.318413 | 0.263 | 0.09 | 1.13356E-20 | Gm26917 |
| 5.29992E-24 | -0.96446 | 0.33 | 0.515 | 1.24463E-19 | Trem2 |
| 1.05338E-23 | 0.500452 | 0.994 | 0.983 | 2.47376E-19 | Gm42418 |
| 7.42319E-19 | 0.559898 | 0.86 | 0.732 | 1.74326E-14 | Ctsd |
| 8.74602E-15 | 0.754721 | 0.325 | 0.177 | 2.05391E-10 | Cpeb2 |
| 1.34178E-13 | 0.599053 | 0.374 | 0.218 | 3.15104E-09 | Serpine2 |
| 2.98826E-13 | 0.546269 | 0.464 | 0.313 | 7.01764E-09 | Sipa1l2 |
| 5.60705E-12 | 0.537993 | 0.504 | 0.353 | 1.31676E-07 | Ctsb |
| 6.14304E-12 | 0.574138 | 0.374 | 0.232 | 1.44263E-07 | Gnas |
| 8.80118E-12 | 0.518538 | 0.488 | 0.337 | 2.06687E-07 | Ints6l |
| 1.0317E-11 | 0.573259 | 0.162 | 0.063 | 2.42285E-07 | Coro2a |
| 2.4351E-11 | -0.39148 | 0.684 | 0.767 | 5.71858E-07 | Lysmd4 |
| 8.00394E-11 | -0.47303 | 0.429 | 0.561 | 1.87965E-06 | Cacnb2 |
| 1.45691E-10 | 0.614078 | 0.183 | 0.085 | 3.42142E-06 | Gcnt2 |
| 1.64283E-10 | 0.487473 | 0.357 | 0.221 | 3.85803E-06 | Plek |
| 1.74494E-10 | 0.287482 | 0.972 | 0.942 | 4.09781E-06 | Plxdc2 |
| 9.7233E-10 | -0.23573 | 0.961 | 0.959 | 2.28342E-05 | Tanc2 |
| 1.17683E-09 | 0.54311 | 0.304 | 0.187 | 2.76366E-05 | Anapc15 |
| 1.55344E-09 | 0.388643 | 0.073 | 0.016 | 3.64809E-05 | Gm26714 |
| 1.94559E-09 | -0.59673 | 0.071 | 0.156 | 4.56902E-05 | Gm31243 |
| 2.60702E-09 | 0.560125 | 0.429 | 0.304 | 6.12231E-05 | Arhgap24 |
| 3.089E-09 | 0.43529 | 0.559 | 0.446 | 7.25421E-05 | Fkbp5 |
| 4.44285E-09 | 0.297657 | 0.937 | 0.918 | 0.000104336 | Csmd3 |
| 6.44601E-09 | 0.534509 | 0.146 | 0.065 | 0.000151378 | Adamts16 |
| 6.82736E-09 | -0.35948 | 0.741 | 0.808 | 0.000160334 | Cx3cr1 |
| 7.07592E-09 | 0.361316 | 0.661 | 0.556 | 0.000166171 | Jmjd1c |
| 7.567E-09 | 0.433044 | 0.554 | 0.435 | 0.000177703 | Snx24 |
| 1.82912E-08 | 0.513476 | 0.138 | 0.062 | 0.000429552 | Rftn1 |
| 2.42111E-08 | 0.475717 | 0.216 | 0.121 | 0.000568573 | Timp2 |
| 2.91803E-08 | -0.49125 | 0.419 | 0.508 | 0.00068527 | Prkca |
| 3.79363E-08 | 0.41423 | 0.298 | 0.192 | 0.000890897 | Dnajc13 |
| 3.94636E-08 | -0.48529 | 0.372 | 0.476 | 0.000926764 | Plxna4 |
| 4.48302E-08 | 0.359022 | 0.726 | 0.633 | 0.001052793 | Ly86 |
| 5.54175E-08 | 0.386656 | 0.484 | 0.367 | 0.001301424 | Dgkd |
| 8.42447E-08 | 0.40104 | 0.512 | 0.403 | 0.001978403 | Cux1 |
| 8.8179E-08 | 0.495756 | 0.271 | 0.173 | 0.002070796 | Hspa8 |
| 1.22639E-07 | -0.3495 | 0.592 | 0.659 | 0.002880063 | Lpcat2 |
| 1.28034E-07 | 0.529111 | 0.219 | 0.128 | 0.003006761 | Lyz2 |
| 1.29964E-07 | 0.218495 | 0.84 | 0.749 | 0.003052085 | Cmss1 |
| 1.58588E-07 | 0.383109 | 0.447 | 0.337 | 0.00372429 | Acer3 |
| 1.58655E-07 | 0.48162 | 0.345 | 0.237 | 0.003725857 | Fau |
| 1.75043E-07 | 0.426551 | 0.123 | 0.055 | 0.004110715 | Sp100 |
| 1.89116E-07 | 0.654351 | 0.564 | 0.455 | 0.004441209 | Apoe |
| 1.92271E-07 | 0.468393 | 0.163 | 0.085 | 0.004515302 | Stat1 |
| 2.1188E-07 | 0.380088 | 0.107 | 0.044 | 0.00497578 | Ptchd1 |

| | | | | | |
|---|---|---|---|---|---|
| 3.54937E-07 | 0.478218 | 0.243 | 0.153 | 0.00833535 | Rpl21 |
| 4.96201E-07 | -0.02363 | 0.348 | 0.236 | 0.011652793 | Nrxn3 |
| 5.19609E-07 | 0.378818 | 0.095 | 0.038 | 0.012202495 | Ifi204 |
| 7.02737E-07 | 0.375259 | 0.337 | 0.234 | 0.016503082 | Atp6v0c |
| 8.42908E-07 | 0.435482 | 0.321 | 0.226 | 0.019794854 | Cables1 |
| 1.0657E-06 | 0.360776 | 0.268 | 0.172 | 0.025026898 | Mpeg1 |
| 1.14975E-06 | 0.418955 | 0.248 | 0.157 | 0.027000657 | Rpl37a |
| 1.18579E-06 | 0.454975 | 0.191 | 0.112 | 0.027847038 | Rpl3 |
| 1.30133E-06 | 0.419254 | 0.29 | 0.195 | 0.030560527 | Ppia |
| 1.58365E-06 | 0.348244 | 0.216 | 0.131 | 0.037190541 | Klhl6 |
| 1.81055E-06 | 0.386755 | 0.091 | 0.038 | 0.042518891 | Per1 |
| 2.11992E-06 | 0.371655 | 0.143 | 0.075 | 0.049784256 | Rps5 |
| 2.31438E-06 | 0.337424 | 0.471 | 0.378 | 0.054350862 | Pvt1 |
| 2.50448E-06 | 0.429413 | 0.124 | 0.062 | 0.058815128 | Rpl7 |
| 2.51401E-06 | 0.397811 | 0.031 | 0.003 | 0.059038982 | Igf1 |
| 2.58068E-06 | 0.347313 | 0.525 | 0.421 | 0.060604736 | Tgfbr2 |
| 2.6651E-06 | 0.372747 | 0.15 | 0.082 | 0.062587156 | Pkib |
| 2.92022E-06 | -0.3216 | 0.474 | 0.563 | 0.06857838 | Zfp710 |
| 3.11812E-06 | 0.47385 | 0.206 | 0.128 | 0.073226006 | Ftl1 |
| 3.26976E-06 | 0.266424 | 0.028 | 0.002 | 0.076787001 | Baiap2l2 |
| 3.4537E-06 | -0.25548 | 0.708 | 0.775 | 0.081106729 | Frmd4a |
| 3.63148E-06 | 0.445782 | 0.185 | 0.111 | 0.08528163 | Acaca |
| 3.69939E-06 | 0.401028 | 0.053 | 0.015 | 0.086876496 | Ccl3 |
| 3.88017E-06 | 0.37594 | 0.546 | 0.454 | 0.091122013 | Cd9 |
| 3.9604E-06 | 0.270579 | 0.617 | 0.51 | 0.093006002 | Sirpa |
| 6.07891E-06 | 0.417464 | 0.218 | 0.14 | 0.142757127 | Dgkz |
| 6.16106E-06 | 0.343273 | 0.053 | 0.016 | 0.144686289 | AY512915 |
| 7.09174E-06 | -0.44122 | 0.324 | 0.402 | 0.166542384 | Dapp1 |
| 7.57865E-06 | 0.426084 | 0.202 | 0.126 | 0.177976977 | Rpsa |
| 7.96275E-06 | 0.391676 | 0.248 | 0.166 | 0.186997325 | Tpt1 |
| 8.15779E-06 | 0.42987 | 0.237 | 0.158 | 0.191577491 | Rpl17 |
| 8.20666E-06 | 0.369573 | 0.163 | 0.095 | 0.192725262 | Stx5a |
| 8.23138E-06 | 0.30663 | 0.443 | 0.338 | 0.193305641 | Fth1 |
| 8.60438E-06 | 0.499599 | 0.329 | 0.247 | 0.202065225 | Cacna1a |
| 9.81137E-06 | 0.33254 | 0.367 | 0.274 | 0.230410259 | Dhx9 |
| 9.85415E-06 | 0.323836 | 0.14 | 0.077 | 0.231414796 | Eif2ak2 |
| 1.01875E-05 | 0.342697 | 0.495 | 0.404 | 0.239242727 | Rcbtb2 |
| 1.02839E-05 | 0.374084 | 0.062 | 0.022 | 0.2415067 | Xaf1 |
| 1.03983E-05 | 0.350293 | 0.304 | 0.215 | 0.244193559 | St6gal1 |
| 1.16032E-05 | 0.384136 | 0.305 | 0.217 | 0.272489615 | Rpl13 |
| 1.17459E-05 | 0.360618 | 0.243 | 0.161 | 0.275841159 | Lars2 |
| 1.19002E-05 | 0.303676 | 0.563 | 0.463 | 0.279463251 | Hdac9 |
| 1.19666E-05 | 0.406346 | 0.189 | 0.118 | 0.28102332 | Rpl13a |
| 1.19768E-05 | -0.44709 | 0.061 | 0.115 | 0.281263484 | Clec4a3 |
| 1.21474E-05 | 0.257553 | 0.69 | 0.578 | 0.285270224 | Snhg11 |
| 1.24197E-05 | 0.293399 | 0.064 | 0.023 | 0.291664157 | Ifih1 |
| 1.28867E-05 | -0.36128 | 0.467 | 0.535 | 0.30263237 | P3h2 |

Vascular cells

| p_val | avg_log2FC | pct.1 | pct.2 | p_val_adj | genes |
|---|---|---|---|---|---|
| 9.75E-11 | 1.602088 | 0.331 | 0.128 | 2.29E-06 | Gm26917 |
| 3.06E-08 | 0.87871 | 0.392 | 0.199 | 0.000719 | Unc13c |
| 8.98E-08 | -1.29531 | 0.003 | 0.101 | 0.002108 | Mybpc1 |
| 1.97E-07 | -0.99494 | 0.024 | 0.14 | 0.004622 | Npy |
| 5.63E-07 | -1.40513 | 0.007 | 0.098 | 0.013213 | Fabp7 |
| 6.56E-07 | -0.88052 | 0.003 | 0.089 | 0.015398 | Gm10863 |
| 1.01E-06 | 0.591903 | 0.195 | 0.062 | 0.023811 | Armh4 |
| 1.13E-06 | 0.703743 | 0.563 | 0.39 | 0.026511 | Fkbp5 |
| 1.24E-06 | 0.660478 | 0.805 | 0.696 | 0.029206 | Ptgds |
| 1.35E-06 | 0.907687 | 0.608 | 0.461 | 0.03164 | Rbfox1 |
| 1.4E-06 | -1.3825 | 0.143 | 0.289 | 0.032781 | Nkain2 |
| 2.13E-06 | 0.688857 | 0.307 | 0.143 | 0.050137 | Aifm3 |
| 3.15E-06 | -1.05342 | 0.003 | 0.08 | 0.073902 | Gdpd4 |
| 4.41E-06 | -0.67216 | 0.007 | 0.086 | 0.103517 | Mfap3l |
| 4.43E-06 | 0.425655 | 0.355 | 0.182 | 0.104062 | Nsa2 |
| 6.15E-06 | -1.80988 | 0.041 | 0.143 | 0.144395 | Brinp3 |
| 6.51E-06 | 0.880013 | 0.355 | 0.211 | 0.152935 | Cpeb2 |
| 6.81E-06 | -0.67307 | 0.007 | 0.083 | 0.159958 | Prima1 |
| 7.41E-06 | -1.0343 | 0.007 | 0.083 | 0.173956 | Gm29521 |
| 8.73E-06 | -0.60256 | 0.003 | 0.074 | 0.204934 | 6030407O03Rik |
| 9.25E-06 | 0.519742 | 0.307 | 0.158 | 0.217135 | Tpm4 |
| 9.25E-06 | 0.632944 | 0.304 | 0.149 | 0.217268 | Ppp1r1a |
| 1.23E-05 | 0.537491 | 0.085 | 0.012 | 0.289411 | Cftr |
| 1.26E-05 | -1.22204 | 0.208 | 0.345 | 0.295409 | Zfhx3 |
| 1.32E-05 | -0.23417 | 1 | 1 | 0.309261 | Malat1 |
| 1.32E-05 | -1.04104 | 0.041 | 0.14 | 0.311004 | Ptprz1 |
| 1.5E-05 | -1.08469 | 0.038 | 0.134 | 0.351225 | Cdh19 |
| 1.53E-05 | 0.527433 | 0.304 | 0.152 | 0.358854 | Acsl1 |
| 1.61E-05 | 0.65044 | 0.253 | 0.119 | 0.378642 | Ube4a |
| 1.64E-05 | -1.18016 | 0.092 | 0.208 | 0.383967 | Ptprt |
| 1.97E-05 | -0.46317 | 0.782 | 0.827 | 0.461872 | Cmss1 |
| 2.03E-05 | 0.972532 | 0.372 | 0.238 | 0.477512 | Btg2 |
| 2.04E-05 | 0.771839 | 0.621 | 0.515 | 0.480016 | Trpm3 |
| 2.18E-05 | -1.18788 | 0.058 | 0.164 | 0.512411 | Sorcs1 |
| 2.72E-05 | -1.3838 | 0.184 | 0.312 | 0.63959 | Sgcd |
| 2.79E-05 | 0.313315 | 0.106 | 0.024 | 0.655529 | Usp38 |
| 3.05E-05 | -1.27871 | 0.044 | 0.14 | 0.716861 | Luzp2 |
| 3.63E-05 | -0.87382 | 0.034 | 0.122 | 0.853171 | Col11a1 |
| 3.64E-05 | -0.85232 | 0.003 | 0.065 | 0.855036 | Gm34838 |
| 3.82E-05 | 0.529855 | 0.358 | 0.205 | 0.897481 | Flt1 |
| 3.91E-05 | -0.70871 | 0.055 | 0.155 | 0.918939 | Fut9 |
| 3.92E-05 | -1.02865 | 0.14 | 0.259 | 0.921363 | Frmd4a |
| 3.98E-05 | 0.490888 | 0.177 | 0.068 | 0.934819 | Ddx1 |
| 4.54E-05 | 0.560175 | 0.461 | 0.295 | 1 | Kcnk2 |
| 4.95E-05 | -0.77018 | 0.027 | 0.11 | 1 | Cdh8 |
| 5.23E-05 | -0.69731 | 0.01 | 0.077 | 1 | Gsap |
